# Variation in Ubiquitin System Genes Creates Substrate-Specific Effects on Proteasomal Protein Degradation

**DOI:** 10.1101/2021.05.05.442832

**Authors:** Mahlon A. Collins, Gemechu Mekonnen, Frank W. Albert

**Affiliations:** Department of Genetics, Cell Biology, and Development University of Minnesota Minneapolis, MN, USA; Department of Biology Johns Hopkins University Baltimore, MD, USA

## Abstract

Ubiquitin-proteasome system (UPS) protein degradation regulates protein abundance and eliminates mis-folded and damaged proteins from eukaryotic cells. Variation in UPS activity influences numerous cellular and organismal phenotypes. However, to what extent such variation results from individual genetic differences is almost entirely unknown. Here, we developed a statistically powerful mapping approach to characterize the genetic basis of variation in UPS activity. Using the yeast *Saccharomyces cerevisiae*, we systematically mapped genetic influences on the N-end rule, a UPS pathway that recognizes N-degrons, degradation-promoting signals in protein N-termini. We identified 149 genomic loci that influence UPS activity across the complete set of N-degrons. Resolving four loci to individual causal nucleotides identified regulatory and missense variants in ubiquitin system genes whose products process (*NTA1*), recognize (*UBR1* and *DOA10*), and ubiquitinate (*UBC6*) cellular proteins. Each of these genes contained multiple causal variants and several individual variants had substrate-specific effects on UPS activity. A *cis*-acting promoter variant that modulates UPS activity by altering *UBR1* expression also alters the abundance of 36 proteins without affecting levels of the corresponding mRNAs. Our results demonstrate that natural genetic variation shapes the full sequence of molecular events in protein ubiquitination and implicate genetic influences on the UPS as a prominent source of post-translational variation in gene expression.

## Introduction

Protein degradation is an essential biological process that occurs continuously throughout the life of a cell. Degradative protein turnover helps maintain protein homeostasis by regulating protein abundance and eliminating misfolded and damaged proteins from cells^1–3^. In eukaryotes, most protein degradation occurs through the concerted actions of the ubiquitin system and the proteasome, together known as the ubiquitin-proteasome system (UPS)^2, 4–7^. Ubiquitin system enzymes bind degradation-promoting signal sequences, termed degrons^8^, in cellular proteins and mark them for degradation by covalently attaching chains of the small protein ubiquitin^5, 9, 10^. The proteasome binds poly-ubiquitinated proteins, then processively unfolds, deubiquitinates, and degrades them to small peptides^11^. The UPS degrades a wide array of proteins spanning diverse biological functions and subcellular localizations. By controlling the turnover of a large fraction of the cellular proteome, the UPS regulates numerous aspects of cellular physiology and function, including gene expression, protein homeostasis, cell growth and division, stress responses, and energy metabolism^1, 3, 5, 12^.

Because of the central role of UPS protein degradation in regulating protein abundance, variation in UPS activity can influence a variety of cellular and organismal phenotypes^1, 3, 13, 14^. Physiological variation in UPS activity enables cells to respond to changes in their internal and external environments. For example, UPS activity increases when misfolded or oxidatively damaged proteins accumulate, preventing these molecules from damaging the cell^15–17^. Conversely, UPS activity decreases during nutrient deprivation, when the energetic demands of UPS protein degradation would be costly to the cell^18–20^. Variation in UPS activity may also create discrepancies between protein degradation and the proteolytic needs of the cell, leading to adverse phenotypic outcomes. For example, age-related declines in UPS activity exacerbate the accumulation of damaged and misfolded proteins that occurs during aging^21–23^. This leads to the formation of protein aggregates and inclusions in a host of age-related neurodegenerative diseases^24–27^. Understanding the sources of variation in UPS activity thus has considerable implications for our understanding of traits influenced by protein degradation, including the many human diseases marked by aberrant UPS activity.

A handful of examples have shown that variation in UPS activity can be caused by individual genetic differences. Rare mutations that ablate or diminish the function of UPS genes impair UPS protein degradation and cause a variety of incurable syndromes. For example, nonsense and frameshift mutations in *UBR1*, an E3 ubiquitin ligase that targets proteins for proteasomal degradation, cause Johansson-Blizzard Syndrome, a developmental disorder that produces morphological abnormalities, cognitive disability, and pancreatic insufficiency^28, 29^. Missense mutations in proteasome genes can cause proteasome-associated autoinflammatory syndrome, a spectrum of disorders associated with lipodystrophy, recurrent fever, and muscle weakness resulting from reduced proteasome activity^30–32^. Genome-wide association studies have linked variants in ubiquitin system^33, 34^ and proteasome genes^35^ to a variety of disorders, but have neither established the specific causal variants nor shown that they alter UPS activity.

These limited examples have left us with a biased, incomplete view of the genetics of UPS activity. In particular, we do not know what fraction of genetic effects on the UPS are due to large-effect variants, such as those in UPS genes that cause rare Mendelian diseases. Most traits are genetically complex, shaped by many loci of small effect and few loci of large effect throughout the genome^36, 37^, suggesting that large-effect variants in UPS genes may represent only one extreme of a continuum of genetic effects on UPS activity. Moreover, variation in UPS activity can differentially affect the degradation of distinct UPS sub-strates^38, 39^. Whether genetic effects on UPS activity affect the turnover of distinct proteins consistently or in a substrate-specific manner remains a fundamentally open question. Finally, we do not know how genetic effects on UPS activity influence other traits. For example, many genetic effects on gene expression influence protein levels without altering mRNA abundance for the same gene^40–48^. These protein-specific effects could arise through differences in UPS activity, but there have been no efforts to understand how natural variation that alters UPS activity influences global protein abundance.

Technical challenges have precluded a comprehensive view of the genetics of UPS activity. Systematically mapping genetic influences on a trait with high statistical power requires assaying large, genetically diverse populations of thousands of individuals^49^. At this scale, *in vitro* biochemical assays of UPS activity are impractical. Several synthetic reporter systems can measure UPS activity with high-throughput *in vivo*^50–52^. However, these systems use genetically encoded fluorescent proteins coupled to degrons to measure UPS activity. When deployed in genetically diverse populations, their output is likely confounded from widespread genetic effects on reporter expression.

Here, we leveraged advances in synthetic reporter design to obtain high-throughput, reporter expression-independent measurements of UPS activity in millions of live, single cells. We use these measurements to map genetic influences on the N-end rule, a UPS pathway that targets degrons in the N-termini (N-degrons^53^) of thousands of endogeneous proteins^1, 54, 55^ and that is linked to diverse physiological functions, including cell growth and division, nutrient sensing and metabolism, meiosis, development, and a host of human diseases^1, 56^. Systematic, statistically powerful genetic mapping revealed the complex, polygenic genetic architecture of UPS activity. We identified 149 loci influencing UPS activity across the complete set of 20 N-degrons. Resolving causal nucleotides at four loci identified regulatory and missense variants in ubiquitin system genes whose products process, recognize, and ubiquitinate cellular proteins. By measuring the effect of a causal variant in the *UBR1* promoter, we implicate genetic influences on UPS activity as a prominent source of post-translational variation in gene expression.

## Results

### Single-Cell Measurements Identify Heritable Variation in UPS Activity

To understand how genetic variation influences UPS activity, we focused on the N-end rule, in which a protein’s N-terminal amino acid functions as an N-degron that results in a protein’s ubiquitination and proteasomal degradation (Figure 1A). The UPS N-end rule can be subdivided into the Arg/N-end and Ac/N-end pathways based on the molecular properties and recognition mechanisms of each pathway’s constituent N-degrons (Figure 1A)^1^. We reasoned that the breadth of degradation signals and recognition mechanisms encompassed in the N-end rule would allow us to identify diverse genetic influences on UPS activity and that the well-characterized effectors of the N-end rule would aid in defining the molecular mechanisms of variant effects. We used a previously-described approach^57^ to generate constructs containing each of the 20 possible N-degrons and appended these sequences to tandem fluorescent timers (TFTs^58^, Figure 1A). TFTs are fusions of a rapidly maturing green fluorescent protein (GFP) and a slower maturing red fluorescent protein (RFP)^58, 59^. The TFT’s ouput, expressed as the -log_2_ RFP / GFP ratio, is directly proportional to its degradation rate and, when fused to N-degrons, measures UPS N-end rule activity^39, 55, 58^. Because the TFT is expressed as a single protein construct, the output of the TFT is also independent of its expression level^39, 55, 58, 60^, enabling its use in genetically diverse populations.

**Figure 1:**
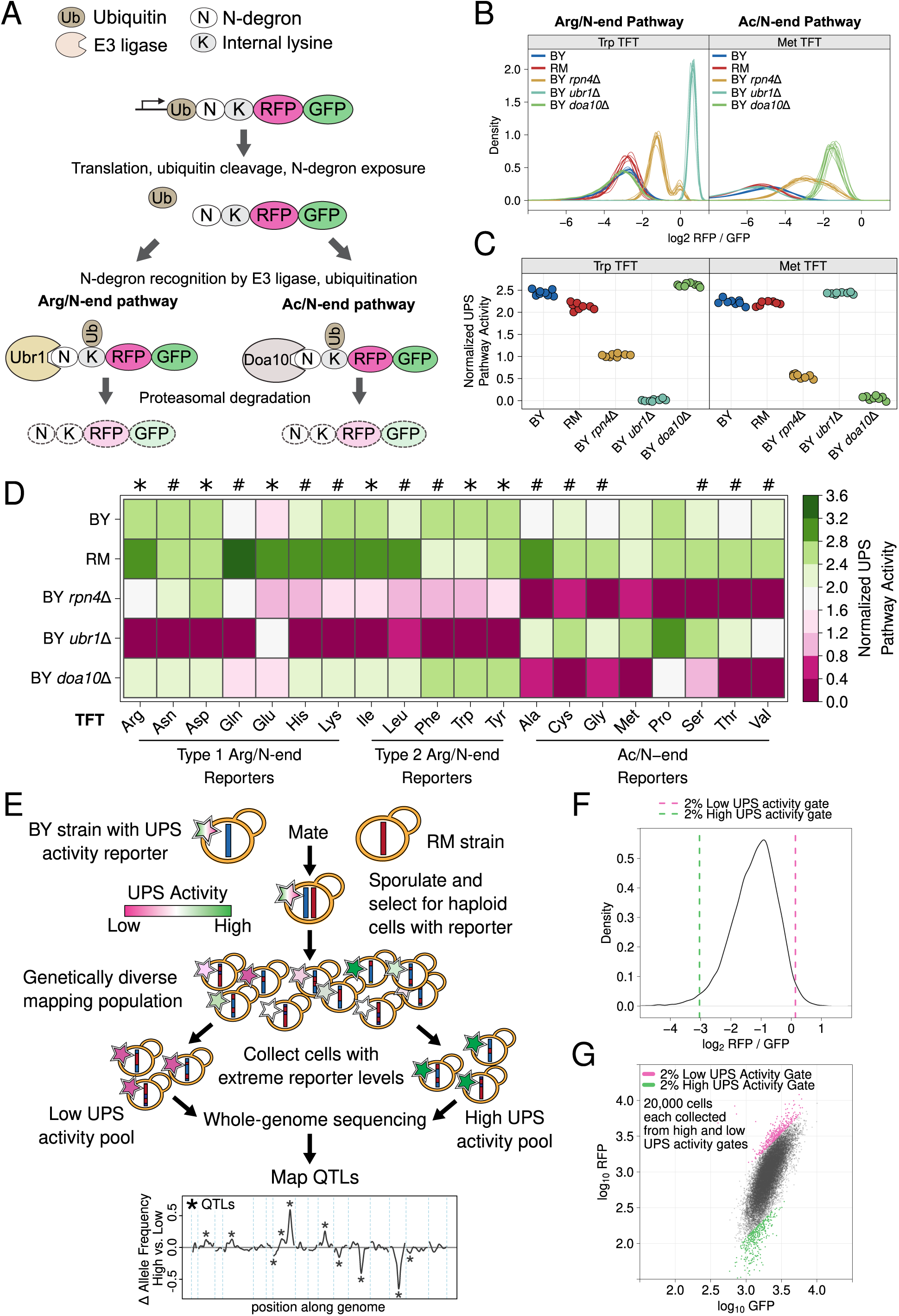
*UPS N-end rule activity reporters and genetic mapping method. A. Schematic of the production and degradation of UPS activity reporters according to the UPS N-end rule. B. Density plots of the log*_2_ *RFP / GFP ratio from 10,000 cells for each of 8 independent biological replicates per strain per reporter for representative Arg/N-end and Ac/N-end UPS activity reporters. “BY” and “RM” are genetically divergent yeast strains. “BY* rpn4Δ*”, “BY* ubr1Δ *”, and “BY* doa10Δ*” carry the indicated gene deletions in the BY background and were used as reporter control strains. C. The median from each biological replicate in B. was scaled, normalized, and plotted as a stripchart such that y axis values are directly proportional to UPS activity. D. Heatmap for all strains and N-degrons using data generated as in C. Symbols above the heatmap denote significant UPS activity differences between BY and RM. “*” indicates 0.05 > Tukey HSD* p *> 1e-6; “#” indicates Tukey HSD* p *< 1e-6. E. Schematic of the bulk segregant analysis genetic mapping method used to identify UPS activity QTLs. F. Density plot of the UPS activity distribution for a genetically diverse mapping population harboring the tryptophan (Trp) N-degron reporter. Dashed vertical lines show the thresholds used to collect cells with extreme UPS activity, which correspond to the high and low UPS activity pools denoted in E. G. Backplot of the cells collected in F. onto a scatter plot of GFP and RFP.*

We characterized the performance of our TFTs by measuring their output in yeast strains with gene deletions that alter UPS activity towards N-end rule substrates. As expected, deleting the E3 ubiquitin ligases of the Arg/N-end (*UBR1*) and the Ac/N-end (*DOA10*) pathways specifically stabilized N-degron TFTs from these pathways (corrected *p <* 0.05 vs. the BY strain, Figure 1B-D, Supplementary Figure 1, Supplementary Table 1). Deleting *RPN4*, which encodes a transcription factor for proteasome genes, reduces proteasome activity^61^ and stabilized reporters from both the Arg/N-end and Ac/N-end pathways (corrected *p <* 0.05 vs. the BY strain, Figure 1B-D, Supplementary Figure 1, Supplementary Table 1). These results show that our TFTs provide sensitive, quantitative, substrate-specific measures of UPS N-end rule activity in multiple genetic backgrounds.

To understand how natural genetic variation influences UPS activity, we compared two genetically divergent *S. cerevisiae* strains, the “BY” laboratory strain and the “RM” vineyard strain^37^. RM had higher UPS activity than BY for 9 of 12 Arg/N-degrons and 6 of 8 Ac/N-degrons (corrected *p <* 0.05, Figure 1D, Supplmentary Figure 1, Supplementary Table 1). BY had higher UPS activity than RM for the phenlyalanine, tryptophan, and tyrosine Arg/N-degrons (corrected *p <* 0.05, Figure 1D, Supplementary Figure 1, Supplementary Table 1). BY and RM had similar activity towards the methionine and proline Ac/N-degrons (corrected *p >* 0.05, Figure 1D, Supplementary Figure 1, Supplementary Table 1). Together, these results show that individual genetic differences create heritable, substrate-specific variation in UPS activity.

### Genetic Mapping Reveals the Complex, Polygenic Genetic Architecture of UPS Activity

We mapped quantitative trait loci (QTLs) for UPS activity using bulk segregant analysis (Figure 1E)^43, 62, 63^. In our implementation, this approach attains high statistical power by comparing whole-genome sequence data from pools of thousands of single cells with extreme UPS activity selected from a large population of haploid, recombinant progeny obtained by crossing BY and RM (Figure 1E-G)^43, 63^. Using this method, we reproducibly identified 149 UPS activity QTLs across the set of 20 N-degron reporters at a false discovery rate of 0.5% (Figure 2A/B, Supplementary Table 2, Supplementary File 1). The number of QTLs per reporter ranged from 1 (for the Ile N-degron) to 15 (for the Ala N-degron) with a median of 7 (Figure 2B, Supplementary Table 2). The distribution of QTL effect sizes (as measured by the allele frequency difference between high and low UPS activity populations), is continuous and consists of many loci of small effect and few loci of large effect (Figure 2C). Thus, UPS activity is a complex, polygenic trait, shaped by variation throughout the genome.

**Figure 2:**
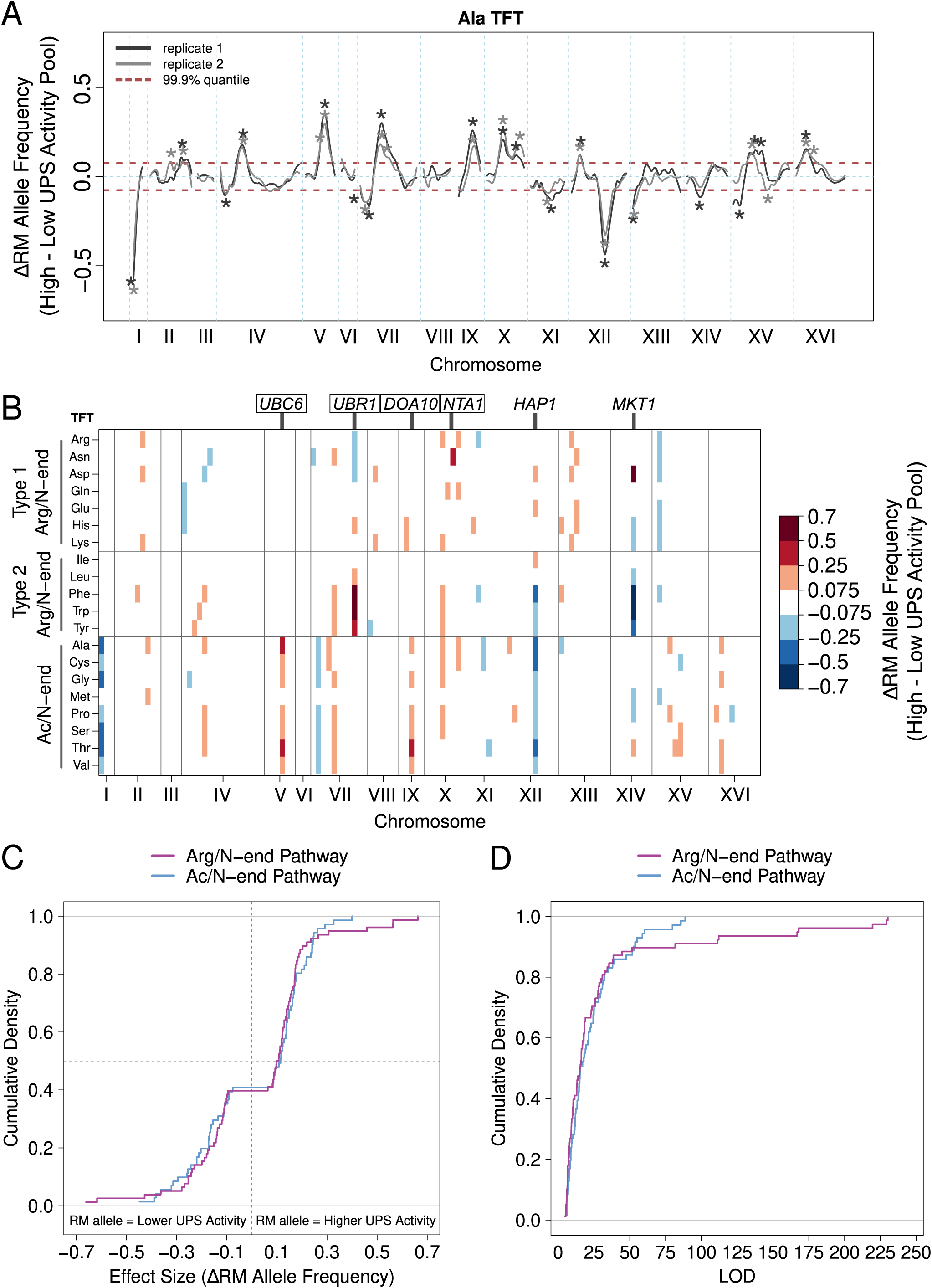
*UPS activity QTL mapping results. A. Results from the alanine (Ala) N-degron reporter illustrate the results and reproducibility of the method. Asterisks denote QTLs, colored by biological replicate. B. QTL mapping results for the 20 N-degrons. Colored blocks of 100 kb denote QTLs detected in each of two independent biological replicates, colored according to the direction and magnitude of the effect size (allele frequency difference between high and low UPS activity pools). Experimentally validated (boxed) and candidate (unboxed) causal genes for select QTLs are annotated above the plot. C. Cumulative distributions of the effect size and direction for Arg/N-end and Ac/N-end QTLs. D. Cumulative distribution of LOD scores for Arg/N-end and Ac/N-end QTLs.*

Analysis of the set of UPS QTLs revealed several patterns. First, many QTLs were found only for individual pathways or N-degrons (Figure 2B, Supplementary Notes 1/2, Supplementary Figure 2), revealing the diversity and specificity of genetic influences on UPS activity. Second, the RM allele was associated with higher UPS activity in a significant majority of UPS QTLs (89 out of 149 [60%; binominal test *p* = 0.021]; Figure 2C). These results are consistent with our observation that RM had higher UPS activity for 15 of 20 N-degrons (Figure 1D, Supplementary Table 1), suggesting that the QTLs we have mapped underlie a substantial portion of the heritable UPS activity difference between BY and RM. Third, the number and patterns of QTLs differed between the Ac/N-end and Arg/N-end pathways (Figure 2B, Supplementary Note 2). The Ac/N-end pathway was affected by a significantly higher number of QTLs per reporter than the Arg/N-end pathway (9 versus 7, respectively, Wilcoxon test *p* = 0.021), while the QTLs with the largest effect sizes were found for the Arg/N-end pathway (Figure 2C/D). The numerous pathway- and N-degron-specific QTLs we identified highlight the considerable complexity of genetic effects on UPS activity.

### Multiple Causal DNA Variants in *UBR1* Create Substrate-Specific Effects on UPS Activity

A QTL on chromosome VII detected with 8 of 12 Arg/N-end reporters (Figure 2B) was centered on *UBR1*, the E3 ligase that recognizes Arg/N-degrons and targets them for UPS protein degradation (Figure 3A). To determine whether *UBR1* contains causal DNA variants for UPS activity towards Arg/N-degrons, we used the CRISPR-swap allelic engineering method^64^ to create BY strains with RM *UBR1* alleles (see “Methods”). Arg/N-end degrons are classified as Type 1 or 2 depending on their Ubr1 binding site^1, 65^. The RM allele at the chromosome VII QTL was associated with decreased UPS activity towards Type 1 Arg/N-degrons and increased UPS activity towards Type 2 Arg/N-degrons (Figure 2B). We therefore tested the effects of the RM *UBR1* alleles on two Type 1 (asparagine [Asn] and aspartate [Asp]) and two Type 2 Arg/N-degrons (tryptophan [Trp] and phenylalanine [Phe]).

**Figure 3:**
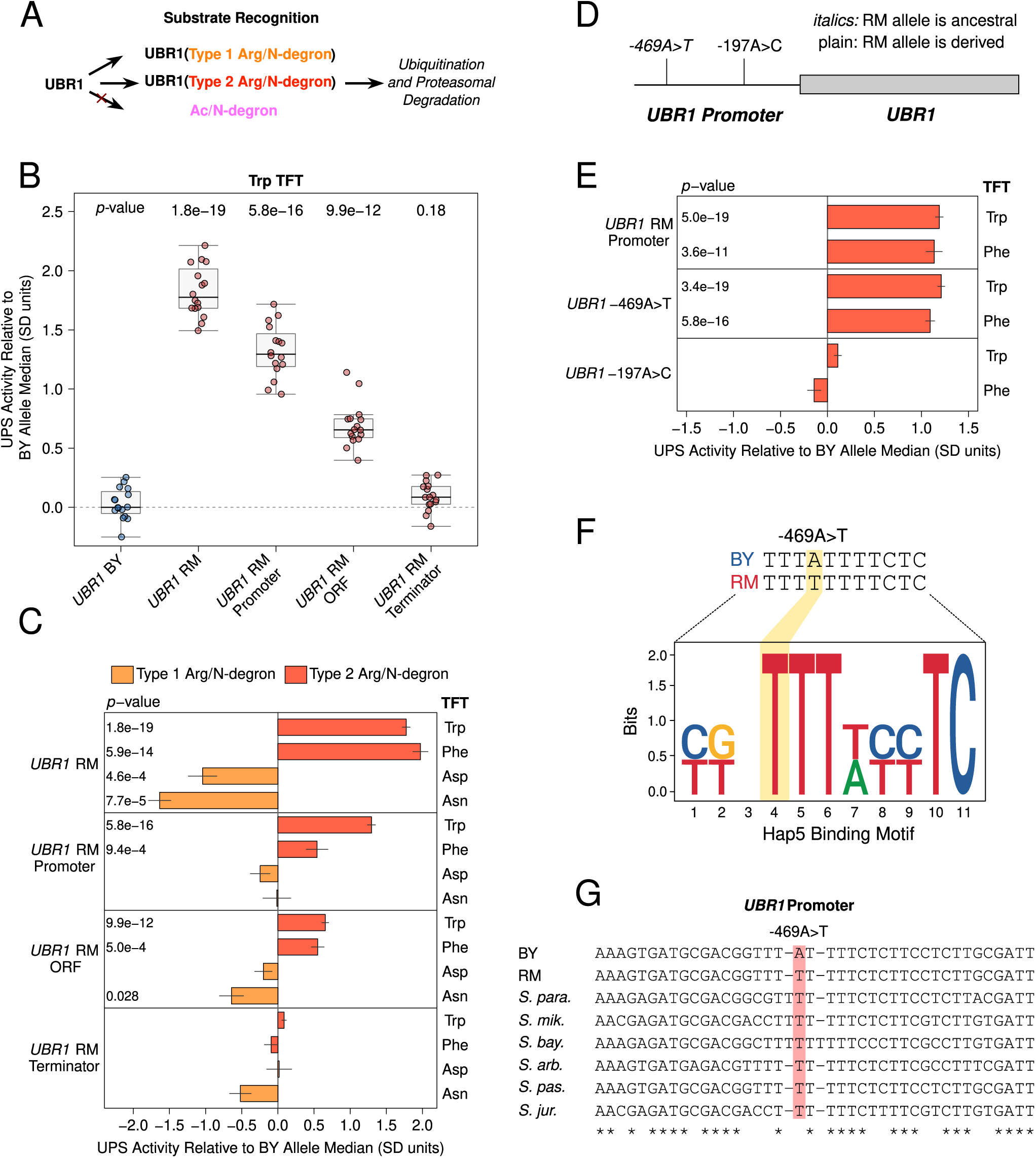
*Substrate-specific effects of* UBR1 *variants on the degradation of Arg/N-degrons. A. Schematic illustrating UBR1’s role in Arg/N-degron recognition. B. Multiple causal DNA variants in UBR1 shape UPS activity towards the Trp N-degron. The BY strain was engineered to contain full or partial RM UBR1 alleles as indicated and UPS activity towards the Trp N-degron TFT was measured by flow cytometry. UPS activity was Z-score normalized and scaled relative to the median of a control BY strain engineered to contain the full BY* UBR1 *allele. Each point in the plot shows the median of 10,000 cells for each of 16 independent biological replicates per strain.* p*-values at the top of the plot display the Benjamini-Hochberg-corrected* p*-value for the t-test of the indicated strain versus the strain with the BY* UBR1 *allele. Box plot center lines, box boundaries, and whiskers display the median, interquartile range, and 1.5 times the interquartile range, respectively. C. Barchart summarizing the effects of RM* UBR1 *alleles on UPS activity towards the indicated Type 1 and 2 Arg/N-degrons using data generated as in B.* p*-values in the plot display the Benjamini-Hochberg-corrected* p *value for the t-test of the indicated strain versus the control strain. D. Diagram of the individual BY / RM* UBR1 *promoter variants. E. as in C., but for the RM* UBR1 *promoter and individual BY / RM* UBR1 *promoter variants. F. Sequence logo of the Hap5 binding motif created by the causal -469A>T* UBR1 *promoter variant. G. Multi-species alignment of the* UBR1 *promoter at the causal -469A>T variant. Abbreviations:* “S. para.”, Saccharomyces paradoxus; “S. mik.”, Saccharomyces mikatae, “S. bay.”, Saccharomyces bayanus; “S. arb”, Saccharomyces arboricola; “S. pas.”, Saccharomyces pastorianus; “S. jur”, Saccharomyces jurei.

Consistent with our QTL mapping results, The RM *UBR1* allele significantly decreased the degradation rate of Type 1 Arg/N-degrons and increased the degradation rate of Type 2 Arg/N-degrons (corrected *p <* 0.05, Figure 3B/C). Thus, *UBR1* is a causal gene for the chromosome VII QTL and BY / RM variants in *UBR1* differentially affect the degradation of Type 1 and 2 substrates of the Arg/N-end pathway.

We then tested the effect of partial RM *UBR1* alleles on UPS activity towards Type 1 Arg/N-degrons to identify the associated causal variants. The RM promoter did not alter the degradation of either the Asn or Asp TFTs (corrected *p >* 0.05, Figure 3C, Supplementary Figure 3). The RM open-reading frame (ORF) significantly decreased the degradation of the Asn, but not the Asp TFT (Figure 3C, Supplementary Figure 3). The RM *UBR1* terminator did not affect UPS activity towards either reporter (corrected *p >* 0.05, Figure 3C, Supplementary Figure 3). Thus, variants in the RM *UBR1* ORF are the main determinant of the gene’s effects on the Asn N-degron. The effects of the RM *UBR1* alleles on the Asp N-degron may be driven by epistatic interactions between variants in the promoter, ORF, and terminator or may be too small to be detected in this assay. Together with our Type 2 Arg/N-degron fine-mapping (described below), these results establish that UPS activity QTLs can contain multiple causal DNA variants in a single gene that can differentially affect the turnover of distinct UPS substrates.

Measurements of the effect of partial RM *UBR1* alleles on the degradation of Type 2 Arg/N-degrons showed drastic differences from their effects on Type 1 Arg/N-degrons (Figure 3C). Both the RM *UBR1* promoter and ORF significantly increased UPS activity towards the Trp and Phe Type 2 Arg/N-degrons (corrected *p <* 0.05, Figure 3B/C, Supplementary Figure 3). The RM *UBR1* terminator did not affect the degradation of either Type 2 Arg/N-degron (corrected *p >* 0.05, Figure 3B / C, Supplementary Figure 3). Thus, the RM *UBR1* promoter and ORF each contain at least one causal variant that increases UPS activity towards Type 2 Arg/N-degron-containing substrates.

To identify individual causal variants, we tested the effect of the two BY / RM *UBR1* promoter variants (Figure 3D) on UPS activity towards Type 2 Arg/N-degrons. The -469A*>*T variant significantly increased the degradation rate of the Trp and Phe N-degrons (corrected *p <* 0.05, Figure 3E, Supplementary Figure 4). By contrast, the -197T*>*G variant had no effect on either N-degron, establishing -469A*>*T as the causal nucleotide in the *UBR1* promoter (corrected *p >* 0.05, Figure 3E, Supplementary Figure 4). The magnitude of the effect caused by -469A*>*T suggests that this variant accounts for the majority of the *UBR1* QTL’s effects on the degradation of Type 2 Arg/N-end substrates (Figure 3B/C/E, Supplementary Figures 3/4).

To gain further insight into the causal -469A*>*T variant, we examined its molecular properties, evolutionary history and population frequency using genome sequence data from a panel of 1,011 *S. cerevisiae* strains^66^. The BY allele of the causal -469A*>*T variant in the *UBR1* promoter disrupts a predicted binding site for the transcription activator Hap5 (Figure 3F) and decreased the output of a synthetic reporter in a massively parallel study of yeast promoter variants^67^. BY carries the derived “A” allele at -469A*>*T, which occurs in a poly(T) motif that is highly conserved across yeast species (Figure 3G). The population frequency of -469A*>*T is 1% and the variant is found in only in the Mosaic Region 1 clade that contains the BY strain (Supplementary Figure 5). These results suggest the BY allele decreases UPS activity by decreasing *UBR1* expression, which we subsequently validated at the RNA and protein levels (Figure 5). Moreover, the derived status and low population frequency of the BY allele at position -469 suggests that it may negatively impact organismal fitness, a notion consistent with the generally deleterious consequences of reduced *UBR1* activity or expression^28, 29, 68^.

### Causal Variants in Functionally Diverse Ubiquitin System Genes Influence UPS Activity

Some of the QTLs with the largest effects were specific to distinct N-end rule pathways or substrates and centered on known ubiquitin system genes (Figure 2B). We used allelic engineering to test whether these genes contained causal DNA variants for UPS activity.

A QTL on chromosome X was specific to the Type 1 asparagine (Asn) N-degron of the Arg/N-end pathway (Figure 2B). The QTL’s peak occurred within *NTA1*, which encodes an amidase that converts N-terminal asparagine and glutamine residues to aspartate and glutamate, respectively (Figure 4A). This processing is necessary to convert Asn and Gln N-ends into functional N-degrons^69^. *NTA1* contains multiple BY / RM promoter variants and two missense variants located on the protein’s exterior surface (Figure 4B/D). Consistent with the chromosome X QTL effect, the RM *NTA1* allele significantly increased the degradation rate of the Asn TFT (corrected *p <* 0.05, Figure 4C, Supplementary Figure 6). The RM *NTA1* promoter did not alter the degradation rate of the Asn TFT (corrected *p >* 0.05, Figure 4C). Instead, the two BY / RM *NTA1* missense variants, D111E and E129G, both influenced degradation of the Asn TFT, but in opposite directions. D111E decreased the Asn TFT’s turnover, while, E129G increased the reporter’s turnover (corrected *p <* 0.05, Figure 4C, Supplementary Figure 6). E129G had an effect that was in the same direction as that of the chromosome X QTL and was approximately threefold greater than the magnitude of the effect of D111E (Figure 4C, Supplementary Figure 6). Thus, at *NTA1*, a large-effect variant masks the effect of a variant with a smaller effect.

**Figure 4:**
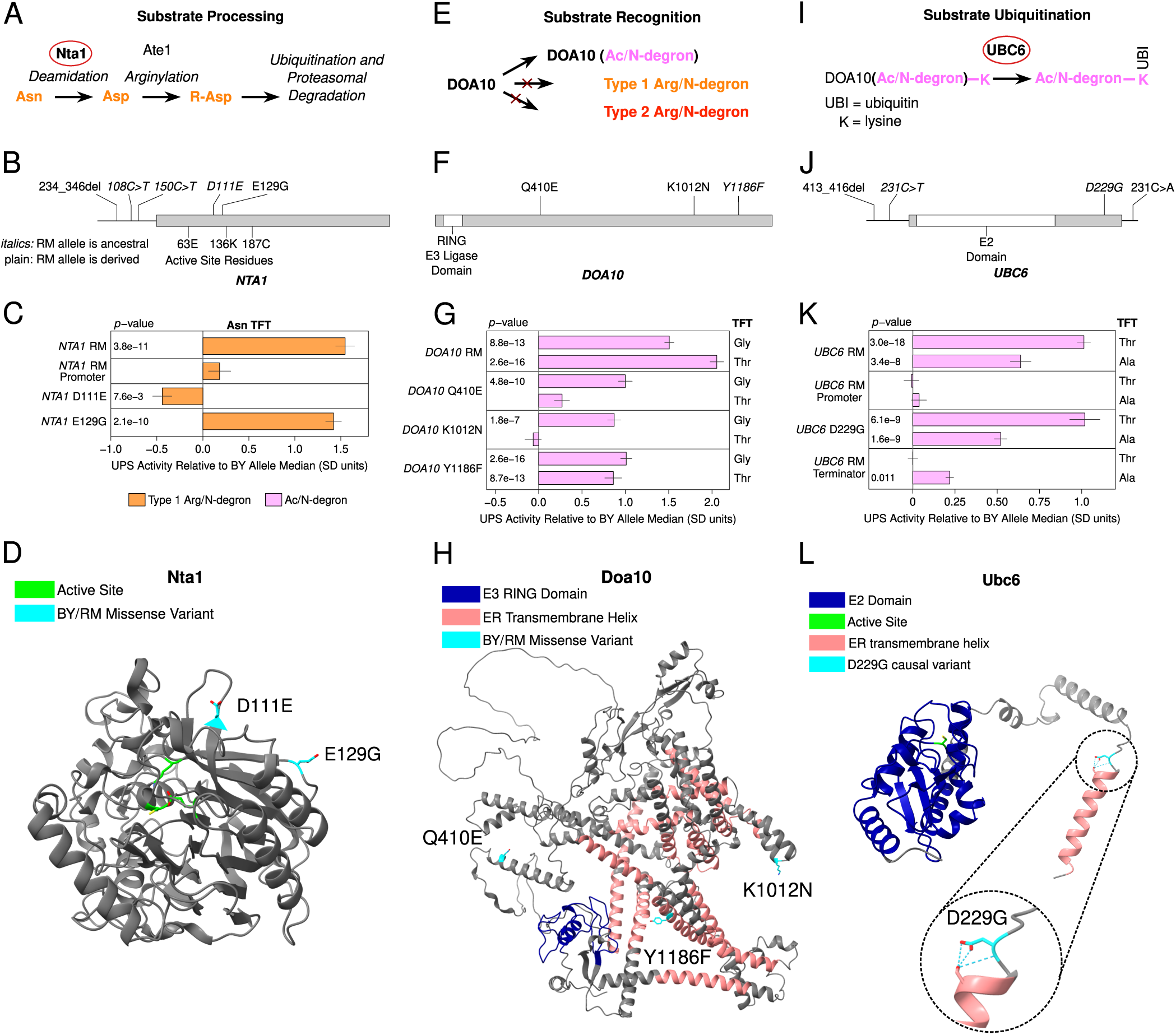
*Identification of causal DNA variants for UPS activity in functionally diverse ubiquitin system genes. A., E., and I. Schematics showing the role of* NTA1 *(A.),* DOA10 *(E.), and* UBC6 *(I.) in UPS substrate processing, recognition, and ubiquitination, respectively. B., F., and J. Location of regulatory and missense variants, as well as active sites and functional domains in* NTA1 *(B.),* DOA10 *(F.), and* UBC6 *(J.). C., G., and K. Fine-mapping results for* NTA1 *(C.),* DOA10 *(G.), and* UBC6 *(K.). Benjamini-Hochberg corrected* p*-values are shown for the t-test of the indicated strain versus a control BY strain engineered to contain the BY allele of each gene. AlphaFold predicted protein structures for Nta1 (D.), Doa10 (H.), and Ubc6 (L.) are shown with causal DNA variants, functional domains, active sites, and transmembrane helices highlighted. The inset in L. shows a predicted hydrogen bonding network at residue 229 in the BY Ubc6 protein.*

A QTL on chromosome IX detected for 6 of 8 Ac/N-end degrons contained *DOA10*, the E3 ligase of the N-end rule that recognizes Ac/N-degrons (Figure 4E). The effect size of this QTL varied between Ac/N-degrons. We, therefore tested the glycine (Gly) and threonine (Thr) reporters to determine whether BY / RM *DOA10* variants exert substrate-specific effects on UPS activity. The RM *DOA10* allele contains three missense variants, Q410E, K1012N, and Y1186F, and does not contain promoter or terminator variants (Figure 4F/H). The RM *DOA10* allele significantly increased the degradation of both the Gly and Thr TFTs (corrected *p <* 0.05, Figure 4H, Supplementary Figure 7). When tested in isolation, all three BY / RM *DOA10* missense variants accelerated the degradation of the Gly TFT (corrected *p <* 0.05, Figure 4H, Supplementary Figure 7). In contrast, only the Y1186F variant significantly increased the degradation of the Thr TFT (Figure 4H, Supplementary Figure 7). The multiple causal variants and substrate-specific effects of individual *DOA10* variants further highlights the complex effects of variation in E3 ubiquitin ligases on UPS activity.

A QTL on chromosome V detected for 7 of 8 Ac/N-degrons contained *UBC6*, the E2 ubiquitin-conjugating enzyme of the Ac/N-end pathway^1, 10^. Ubc6 pairs with Doa10 to ubiquitinate substrates of the Ac/N-end pathway (Figure 4I)^70, 71^. The RM *UBC6* allele contains a 3 base pair deletion in the promoter, 1 missense variant, and 1 terminator variant (Figure 4F). Using the threonine (Thr) and alanine (Ala) TFTs, we established that *UBC6* is a causal gene for this QTL (corrected *p <* 0.05, Figure 4K, Supplementary Figure 8). The D229G missense variant altered UPS activity towards the alanine (Ala) and threonine (Thr) Ac/N-degrons (corrected *p <* 0.05, Figure 4K, Supplementary Figure 8). The *UBC6* RM terminator also significantly increased the degradation rate of the Ala, but not Thr TFT, further establishing the substrate-specific effects of genetic variation on UPS activity (Figure 4K, Supplementary Figure 8). Our results with *UBC6* show that genetic variation influences UPS activity through effects on substrate ubiquitination by E2 ubiquitin ligases, as well as substrate recognition by the E3 ligases Ubr1 and Doa10 described above.

Knowledge of the causal nucleotides in *NTA1*, *DOA10*, and *UBC6* allowed us to examine their molecular properties, evolutionary histories, and population frequencies. A notable feature of causal missense variants was their distal location relative to the active site of the corresponding protein (Figure 4D/H/L). The *NTA1* causal variants occurs on the protein’s exterior surface (Figure 4D), while the causal variants in *DOA10* and *UBC6* occur in or near transmembrane helices that anchor these proteins to the endoplasmic reticulum membrane (Figure 4H/L). Thus, in addition to effects on ubiquitin system gene expression (e.g., *UBR1* above), the molecular basis for the continuous distribution of variant effects on UPS activity may also involve subtle alterations to the stability, localization, or physical interactions of ubiquitin system proteins^72^.

The BY alleles of the causal *DOA10* Y1186F and *UBC6* D229G variants are derived, at low population frequencies (5.1% and 2.2%, respectively), and occur in only 3 non-BY clades (Supplementary Figure 5), similar to the results we obtained for the -469A*>*T *UBR1* variant. Given the generally deleterious effects of reduced UPS activity^12, 13^, these variants may be subject to purifying selection. In contrast, the RM allele of the causal *NTA1* E129G variant is derived, common (51.5% population frequency), and found in most clades (Supplementary Figure 5). The derived RM allele of the causal *NTA1* E129G variant may have been able to rise to comparatively high population frequency because deleting *NTA1* does not decrease competitive fitness^69^.

We examined additional QTLs to nominate candidate causal genes. The most frequently observed UPS QTL was detected for 8 of 8 Ac/N-end and 6 of 12 Arg/N-end TFTs and was located on chromosome XII in the immediate vicinity of a Ty1 insertion in the *HAP1* transcription factor in the BY strain^73^(Figure 2B, Supplementary Table 2). The Ty1 insertion in *HAP1* exerts strongly pleiotropic effects on gene expression, altering the expression of 3,755 genes^74^. Similarly, a QTL on chromosome XIV affected 10 of our 20 N-degron TFTs and was located in the *MKT1* gene (Figure 2B, Supplementary Table 2). *MKT1* encodes a multi-functional RNA binding protein involved in the post-transcriptional regulation of gene expression and is the causal gene for other QTLs previously mapped in the BY/RM cross^75–77^. *HAP1* and *MKT1* are the likely causal genes for the chromosome XII and XIV QTLs, showing that genetic variation may also shape UPS activity through indirect effects on genes with no known connection to the UPS.

Taken together, our analysis of causal genes and nucleotides illustrates the breadth and diversity of genetic influences on UPS activity. Each fine-mapped causal gene harbored multiple causal variants that may differentially affect distinct UPS substrates. Regulatory and missense variants in ubiquitin system genes shape the full sequence of molecular events, including substrate processing, recognition, and ubiquitination, that lead to a protein’s degradation by the UPS.

### Protein-Specific Effects of *UBR1* -469A*>*T on Gene Expression

Previous efforts to understand how genetic variation influences gene expression have revealed considerable discrepancies between genetic effects on mRNA versus protein abundance^40–48^. Many gene expression QTLs alter protein abundance without detectable effects on mRNA levels. We reasoned that protein-specific gene expression QTLs could arise through effects on UPS protein degradation. To test this idea and explore how variant effects on UPS activity influence other aspects of cellular physiology, we measured global gene expression at the protein and RNA levels in the BY strain and a BY strain engineered to contain the causal -469A*>*T RM allele in the *UBR1* promoter (“BY *UBR1* -469A*>*T”). As expected, the derived BY allele decreased *UBR1* protein and RNA levels (Figure 5A/B).

**Figure 5:**
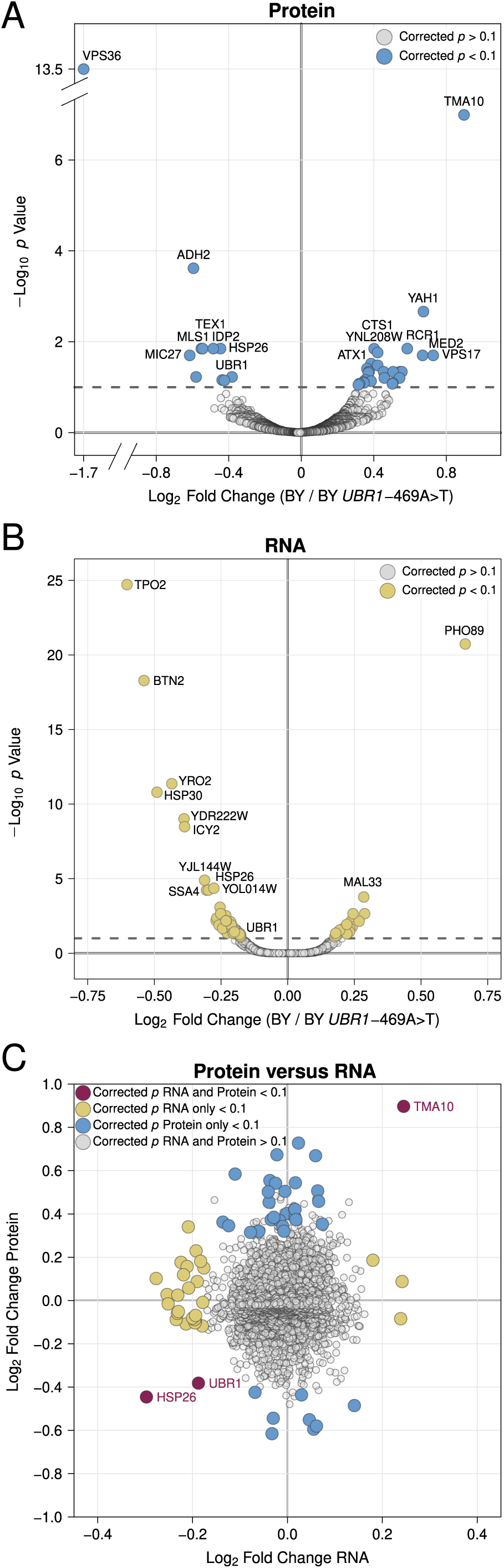
*Proteomic and RNA-seq analysis of the effect* UBR1 *-469A>T promoter variant on gene expression. A. Protein fold-change versus statistical significance for BY versus BY* UBR1 *-469A>T for all detected proteins. Differentially abundant proteins are shown in blue. B. RNA fold-change versus statistical significance for BY versus BY* UBR1 *-469A>T for all detected transcripts. Differentially expressed transcripts are shown in yellow. C. Scatterplot comparing changes in protein and RNA abundance caused by* UBR1 *-469A>T.*

Out of 3,774 proteins quantified by mass spectrometry, 39 proteins were differentially abundant at a 10% FDR (Figure 5A, Supplementary Table 8). Consistent with the reduced UPS activity conferred by the BY *UBR1* allele, 71% (28 of 39) of differentially abundant proteins were increased in BY (Figure 5A, Supplementary Table 4). No Gene Ontology or Reactome pathway terms were enriched in our set of differentially abundant proteins. This result is consistent with recent observations that suggest that altering *UBR1* expression exerts broad effects on protein degradation or related processes controlling protein abundance and that protein sequences, rather than function or subcellular localization, are the primary determinants of degradation rates^38, 39^.

To determine whether differences in protein abundance were reflected at the MRNA level, we used RNA-seq to quantify the levels of 5,675 transcripts. A total of 78 transcripts were differentially expressed between BY and BY *UBR1* -469A*>*T at a 10% FDR (Figure 5B, Supplementary Table 5). Only three genes, *UBR1*, *HSP26*, and *TMA10*, showed significant and concordant changes at the RNA and protein levels (Figure 5C). In contrast to our proteomics results, the BY allele tended to decrease mRNA abundance, causing lower expression at 67 of the 78 (86%) differentially expressed genes (Figure 5B, Supplementary Table 5). Multiple proteostasis-related pathways were enriched among the differentially abundant transcripts (Supplementary Figure 9), driven by the decreased transcript abundance for genes such as *UBR1* and the chaperones *HSP26*, *HSP30*, *HSP31*, and *HSP82* in BY. Our results add to the emerging view of the complex, protein-specific influences of genetic variation on gene expression^40–48^. Specifically, a non-coding variant that decreases expression of a single E3 ubiquitin ligase increases the levels of dozens of proteins without detectable effects on transcript abundance, implicating genetic effects on UPS activity as a prominent source of post-translational variation in gene expression.

## Discussion

Protein degradation by the UPS is an essential biological process that influences virtually all aspects of eukaryotic cellular physiology^1, 3, 13, 17^. Understanding the sources of variation in UPS activity thus has considerable implications for our understanding of numerous cellular and organismal traits, including human health and disease^13, 14, 26, 78^. Our statistically powerful, systematic genetic mapping of the N-end rule has revealed that individual genetic differences create heritable variation in UPS protein degradation. Genetic effects on UPS activity are numerous and comprise a continuous distribution of many loci with small effects and few loci of large effect (Figure 2), similar to other complex traits^36, 37^. Previous efforts to understand how individual genetic differences cause variation in UPS activity have focused on individual disease-causing mutations in UPS genes^28, 78–81^. Our results show that these large-effect mutations in UPS genes sit atop one extreme of a continuous distribution of variant effects that is dominated by many loci of small effect. Aberrant UPS activity is a hallmark of many common diseases with a poorly-understood, complex genetic basis^14, 26, 27^. Our results raise the possibility that the effects of many common, small-effect alleles may contribute to the risk of these diseases through their effects on UPS activity.

Using genome engineering, we experimentally identified causal regulatory and missense variants in four functionally distinct ubiquitin system genes. A major function of the ubiquitin system is conferring specificity to UPS protein degradation^9, 82, 83^. Non-ubiquitinated proteins are blocked by the proteasome’s 19S regulatory particle from degradation by the 20S catalytic core^84^. The selective binding of ubiquitinated substrates by the 19S regulatory particle ensures that only those proteins targeted for degradation enter the proteasome. The activity of the ubiquitin system towards distinct substrates is highly variable, even for proteins degraded by the same UPS pathway^6, 38, 55^. Consistent with these observations, the effects of causal ubiquitin system gene variants were highly substrate-specific (Figures 3/4). Our results raise the question of whether UPS protein degradation is also shaped by variation in proteasome genes and whether any such effects would be less substrate-specific than those in the ubiquitin system. Given the multiple, large-effect QTLs arising from ubiquitin system genes, detecting genetic influences on proteasome activity may benefit from assays that can measure proteasome activity independently of the ubiquitin system.

The remarkable complexity in causal variants we uncovered underscores the challenge of predicting variant effects on UPS protein degradation. Similar to recent results^47, 85^, each of the four QTL regions we fine-mapped contained multiple causal variants in a single gene (Figures 3/4). In the case of *NTA1*, we observed that the effect of the D111E variant was likely masked during QTL mapping by the larger effect of the E129G variant (Figure 4C), highlighting the need to test individual variants in isolation. Causal variants may also exert substrate-specific effects on UPS protein degradation. We observed multiple instances where the magnitude of a causal variant’s effect varied between substrates. In the case of *UBR1*, the RM *UBR1* ORF exerted significant, discrepant effects on the degradation of Arg/N-degrons (Figure 3C). Recent efforts have established that a protein’s sequence critically determines how its degradation is altered by changes in UPS activity^38^. Thus, a complete understanding of a given variant’s influence on UPS protein degradation will require testing its effect on the turnover of multiple substrates with diverse sequence compositions.

Our results suggest that genetic effects on UPS activity are an important source of post-translational variation in gene expression. A promoter variant that reduces UPS activity by decreasing *UBR1* expression alters the abundance of dozens of proteins without detectable effects on levels of the corresponding mRNA transcripts (Figure 5). Ubr1 and Doa10 target distinct sets of cellular proteins^38, 39, 55^. Their genes each contain multiple causal variants that differentially affected individual N-degrons and thus, potentially, endogenous cellular proteins. Similar effects arising from variation in the approximately 100 E3 ubquitin ligases encoded in the *S. cerevisiae* genome^10^ may help explain the numerous protein-specific gene expression QTLs^40, 42–44^. Such effects could be even more prevalent in the human genome, which encodes an estimated 600 E3 ubiquitin ligases^86^.

We have developed a generalizable framework for mapping genetic influences on protein degradation. Our results lay important groundwork for future efforts to understand how heritable differences in UPS activity contribute to variation in complex cellular and organismal traits, including the many diseases marked by aberrant UPS activity.

## Methods

### Tandem Fluorescent Timer Ubiquitin-Proteasome System Activity Reporters

We used tandem fluorescent timers (TFTs) to measure ubiquitin-proteasome system (UPS) activity. TFTs are fusions of two fluorescent proteins (FPs) with distinct spectral profiles and maturation kinetics^58, 59^. In the most common implementation, a TFT consists of a faster maturing green fluorescent protein (GFP) and a slower maturing red fluorescent protein (RFP). Because the FPs in the TFT mature at different rates, the RFP / GFP ratio changes over time. If the degradation rate of a TFT exceeds the maturation rate of the RFP, the -log_2_ RFP / GFP ratio is directly proportional to the construct’s degradation rate^58, 59^. When fused to N-degrons, the TFT’s RFP / GFP ratio measures UPS N-end rule activity^58, 60^. The RFP / GFP ratio is also independent of the TFT’s expression level,^39, 58, 59^ preventing confounding from genetic effects on reporter expression in genetically diverse cell populations.

We used fluorescent proteins from previously characterized TFTs in our experiments^58–60, 87^. superfolder GFP^88^ (sfGFP) was used as the faster maturing FP in all TFTs. sfGFP matures in approximately 5 minutes and has excitation and emission maximums of 485 nm and 510 nm, respectively^88^. The slower maturing FP in each TFT was either mCherry or mRuby. mCherry matures in approximately 40 minutes and has excitation and emission maximums of 587 nm and 610 nm, respectively^89^. The mCherry-sfGFP TFT can detect degradation rate differences in substrates with lifetimes of approximately 80 minutes^58, 87^. mRuby matures in approximately 170 minutes and has excitation and emission maximums of 558 nm and 605 nm, respectively^90^. The mRuby-sfGFP TFT can detect degradation rate differences in substrates with lifetimes of approximately 340 minutes, although it is less sensitive than the mCherry-sfGFP TFT for substrates with half-lives less than 80 minutes^58, 87^. All TFT fluorescent proteins are monomeric. We separated green and red FPs in each TFT with an unstructured 35 amino acid linker sequence to minimize fluorescence resonance energy transfer^58^.

### Construction of Arg/N-end and Ac/N-end Pathway TFTs

To generate TFT constructs with defined N-terminal amino acids, we used the ubiquitin-fusion technique^1, 6, 57^, which involves placing a ubiquitin moiety immediately upstream of a sequence encoding fthe desired N-degron. During translation, ubiquitin-hydrolases cleave the ubiquitin moiety, exposing the N-degron (Figure 1A). We synthesized DNA (Integrated DNA Technologies [IDT], Coralville, Iowa, USA) encoding the *Saccharomyces cerevisiae* ubiquitin sequence and a peptide linker sequence derived from *Escherichia coli β*-galactosidase previously used to identify components of the Arg/N-end and Ac/N-end pathways^6^. The peptide linker sequence is unstructured and contains internal lysine residues required for ubiquitination and degradation by the UPS^6, 71^. Peptide linkers encoding the 20 possible N-terminal amino acids were made by PCR amplifying the linker sequence using oligonucleotides encoding each unique N-terminal amino acid (Supplementary Table 6).

We then devised a general strategy to assemble TFT-containing plasmids with defined N-terminal amino acids (Supplementary Figure 10). We first obtained sequences encoding each reporter element by PCR or DNA synthesis. We codon-optimized the sfGFP, mCherry, mRuby, and the TFT linker sequences for expression in *S. cerevisiae* using the Java Codon Adaptation Tool (JCaT)^91^ and synthesized DNA fragments encoding each sequence (IDT). We used the *TDH3* promoter to drive expression of each TFT reporter. The *TDH3* promoter was PCR-amplified from Addgene plasmid #67639 (a gift from John Wyrick). We used the *ADH1* terminator in all TFT constructs, which we PCR amplified from Addgene plasmid #67639. We used the KanMX cassette^92^, which confers resistance to G418, as the selection module for all TFT constructs and obtained this sequence by PCR amplification from Addgene plasmid #41030 (a gift from Michael Boddy). Thus, each construct has the general structure of *TDH3* promoter, N-degron, linker sequence, TFT, *ADH1* terminator, and the KanMX resistance cassette (Supplementary Figure 10). Based on the half-lives of N-degrons^1, 6, 71^, we used the mCherry-fGFP TFT for all Arg/N-end constructs and the mRuby-sfGFP TFT for all Ac/N-end constructs.

We used Addgene plasmid #35121 (a gift from John McCusker) to construct all TFT plasmids. Digesting this plasmid with BamHI and EcoRV restriction enzymes produces a 2451 bp fragment that we used as a vector backbone for TFT plasmid assembly. We obtained a DNA fragment containing 734 bp of sequence upstream of the *LYP1* start codon, a SwaI restriction site, and 380 bp of sequence downstream of the *LYP1* stop codon by DNA synthesis (IDT). We performed isothermal assembly cloning using the New England Biolabs (NEB; Ipswich, MA, USA) HiFi Assembly Cloning Kit (NEB) to insert the *LYP1* homology sequence into the BamHI/EcoRV digest of Addgene plasmid #35121 to create the final backbone plasmid BFA0190 (Supplementary Figure 10). We then combined SwaI digested BFA0190 and the components of each TFT reporter and used the NEB HiFi Assembly Kit (NEB) to produce each TFT plasmid. The 5’ and 3’ *LYP1* sequences in each TFT contain naturally-occurring SacI and BglII restriction sites, respectively. We digested each TFT plasmid with SacI and BglII (NEB) to obtain a linear DNA transformation fragment (Supplementary Figure 10). The flanking *LYP1* homology and kanMX module in each TFT construct allows selection for reporter integration at the *LYP1* locus using G418^93^ and the toxic amino acid analogue thialysine (S-(2-aminoethyl)-L-cysteine hydrochloride)^94–96^. The sequence identity of all assembled plasmids was verified by Sanger sequencing. The full list of plasmids used in this study is found in Supplementary Table 7.

### Yeast Strains and Handling

We used two strains of the yeast *Saccharomyces cerevisiae* to characterize our TFT reporters and perform genetic mapping of UPS activity. The haploid BY strain (genotype: *MATa his3*Δ *ho*Δ) is closely related to the *S. cerevisiae* S288C laboratory strain. The second mapping strain, RM, was originally isolated from a California vineyard and is haploid with genotype *MATα can1*Δ*::STE2pr-SpHIS5 his3*Δ*::NatMX AMN1-BY ho*Δ*::HphMX URA3-FY*. BY and RM differ at 1 nucleotide per 200 base pairs on average, such that approximately 45,000 single nucleotide variants (SNVs) between the strains can serve as markers in a genetic mapping experiment^37, 43, 44, 63^.

We built additional strains for characterizing our UPS activity reporters by deleting individual UPS genes from the BY strain. Each deletion strain was constructed by replacing the targeted gene with the NatMX cassette^93^, which confers resistance to the antibiotic nourseothricin. We PCR amplified the NatMX cassette from Addgene plasmid #35121 using primers with homology to the 5’ upstream and 3’ downstream sequences of the targeted gene. The oligonucleotides for each gene deletion cassette amplification are listed in Supplementary Table 6. We created a BY strain lacking the *UBR1* gene, which encodes the Arg/N-end pathway E3 ligase Ubr1. We refer to this strain hereafter as “BY ubr1Δ”. We created a BY strain (“BY doa10Δ”) lacking the *DOA10* gene that encodes the Ac/N-end pathway E3 ligase Doa10. Finally, we created a BY strain (“BY rpn4Δ”) lacking the *RPN4* that encodes the proteasome transcription factor Rpn4p. Table 1 lists these strains and their full genotypes. Supplementary Table 8 contains the complete list of strains used in this study.

**Table 1:** *Strain genotypes.*

| Short Name | Genotype | Antibiotic Resistance | Auxotrophies |
| --- | --- | --- | --- |
| BY | *MATa his3Δ hoΔ* |  | histidine |
| RM | *MATα can1Δ::STE2pr-SpHIS5*<br>*his3Δ::NatMX hoΔ::HphMX* | clonNAT, hygromycin | histidine |
| BY *rpn4Δ* | *MATa his3Δ hoΔ rpn4Δ::NatMX* | clonNAT | histidine |
| BY *ubr1Δ* | *MATa his3Δ hoΔ ubr1Δ::NatMX* | clonNAT | histidine |
| BY *doa10Δ* | *MATa his3Δ hoΔ doa10Δ::NatMX* | clonNAT | histidine |

Table 2 describes the media formulations used for all experiments. Synthetic complete amino acid powders (SC -lys and SC -his -lys -ura) were obtained from Sunrise Science (Knoxville, TN, USA). Where indicated, we added the following reagents at the indicated concentrations to yeast media: G418, 200 mg/mL (Fisher Scientific, Pittsburgh, PA, USA); ClonNAT (nourseothricin sulfate, Fisher Scientific), 50 mg/L; thialysine (S-(2-aminoethyl)-L-cysteine hydrochloride; MilliporeSigma, St. Louis, MO, USA), 50 mg/L; canavanine (L-canavanine sulfate, MilliporeSigma), 50 mg/L.

**Table 2:** *Media formulations.*

| Media Name | Abbreviation | Formulation |
| --- | --- | --- |
| Yeast-Peptone-Dextrose | YPD | 10 g/L yeast extract |
|  |  | 20 g/L peptone |
|  |  | 20 g/L dextrose |
| Synthetic Complete | SC | 6.7 g/L yeast nitrogen base |
|  |  | 1.96 g/L amino acid mix -lys |
|  |  | 20 g/L dextrose |
| Haploid Selection | SGA | 6.7 g/L yeast nitrogen base |
|  |  | 1.74 g/L amino acid mix -his -lys -ura |
|  |  | 20 g/L dextrose |
| Sporulation | SPO | 1 g/L yeast extract |
|  |  | 10 g/L potassium acetate |
|  |  | 0.5 g/L dextrose |

### Yeast Transformation

We used a standard yeast transformation protocol to construct reporter control strains and build strains with UPS activity reporters^97^. In brief, we inoculated yeast strains growing on solid YPD medium into 5 mL of YPD liquid medium for overnight growth at 30 °C. The following morning, we diluted 1 mL of saturated culture into 50 mL of fresh YPD and grew the cells for 4 hours. The cells were then successively washed in sterile ultrapure water and transformation solution 1 (10 mM Tris HCl [pH 8.0], 1 mM EDTA [pH 8.0], and 0.1 M lithium acetate). At each step, we pelleted the cells by centrifugation at 3,000 rpm for 2 minutes in a benchtop centrifuge and discarded the supernatant. The cells were suspended in 100 *µ*L of transformation solution 1 along with 50 *µ*g of salmon sperm carrier DNA and 300 ng of transforming DNA. The cells were incubated at 30 °C for 30 minutes and 700 *µ*L of transformation solution 2 (10 mM Tris HCl [pH 8.0], 1 mM EDTA [pH 8.0], and 0.1 M lithium acetate in 40% polyethylene glycol [PEG]) was added to each tube, followed by a 30 minute heat shock at 42 °C. We then washed the transformed cells in sterile, ultrapure water. We added 1 mL of liquid YPD medium to each tube and incubated the tubes for 90 minutes with rolling at 30 °C to allow for expression of the antibiotic resistance cassettes. After washing with sterile, ultrapure water, we plated 200 *µ*L of cells on solid SC -lys medium with G418 and thialysine, and, for strains with the NatMX cassette, clonNAT. For each strain, we streaked multiple independent colonies (biological replicates) from the transformation plate for further analysis as indicated in the text. We verified reporter integration at the targeted genomic locus by colony PCR^98^. The primers used for these experiments are listed in Supplementary Table 6.

### Yeast Mating and Segregant Populations

We created populations of genetically variable, recombinant cells (“segregants”) for genetic mapping using a modified synthetic genetic array (SGA) approach^95, 96^. We first mated BY strains with a given UPS activity reporter to RM by mixing freshly streaked cells of each strain on solid YPD medium. For each UPS activity reporter, we mated two independently-derived clones (biological replicates) to the RM strain. Cells were grown overnight at 30 °C and we selected for diploid cells (successful BY-RM matings) by streaking mated cells onto solid YPD medium with G418 (which selects for the KanMX cassette in the TFT in the BY strain) and clonNAT (which selects for the NatMX cassette in the RM strain). We inoculated 5 mL of YPD with freshly streaked diploid cells for overnight growth at 30 °C. The next day, we pelleted the cultures, washed them with sterile, ultrapure water, and resuspended the cells in 5 mL of SPO liquid medium (Table 2). We sporulated the cells by incubating them at room temperature with rolling for 9 days. After confirming sporulation by brightfield microscopy, we pelleted 2 mL of culture, washed cells with 1 mL of sterile, ultrapure water, and resuspended cells in 300 *µ*L of 1 M sorbitol containing 3 U of Zymolyase lytic enzyme (United States Biological, Salem, MA, USA) to degrade ascal walls. Digestions were carried out at 30 °C with rolling for 2 hours. We then washed the spores with 1 mL of 1 M sorbitol, vortexed for 1 minute at the highest intensity setting, resuspended the cells in sterile ultrapure water, and confirmed the release of cells from ascii by brightfield microscopy. We plated 300 *µ*l of cells onto solid SGA medium containing G418 and canavanine. This media formulation selects for haploid cells with (1) a UPS activity reporter via G418, (2) the *MATa* mating type via the *Schizosaccharomyces pombe HIS5* gene under the control of the *STE2* promoter (which is only active in *MATa* cells), and (3) replacement of the *CAN1* gene with *S. pombe HIS5* via the toxic arginine analog canavanine^95, 96^. Haploid segregant populations were grown for 2 days at 30 °C and harvested by adding 10 mL of sterile, ultrapure water and scraping the cells from each plate. We pelleted each cell suspension by centrifugation at 3000 rpm for 10 minutes and resuspended the cells in 1 mL of SGA medium. We added 450 *µ*L of 40% (v/v) sterile glycerol solution to 750 *µ*L of segregant culture and stored samples in screw cap cryovials at *−*80 °C. We stored 2 independent sporulations of each reporter (derived from our initial matings) as independent biological replicates.

### Flow Cytometry

We measured UPS activity by flow cytometry as follows. Yeast strains were manually inoculated into 400 *µ*L of liquid SC -lys medium with G418 and grown overnight in 2 mL 96 well plates at 30.0 °C with 1000 rpm mixing using a MixMate (Eppendorf, Hamburg, Germany). The following morning, we inoculated a fresh 400 *µ*L of G418-containing SC -lys media with 4 *µ*L of each saturated culture. Cells were grown for an additional 3 hours prior to analysis by flow cytometry. All flow cytometry experiments were performed on an LSR II flow cytometer (BD, Franklin Lakes, NJ, USA) equipped with a 20 mW 488 nm laser with 488/10 and 525/50 filters for measuring forward/side scatter and sfGFP, respectively, as well as a 40 mW 561 nm laser and a 610/20 filter for measuring mCherry and mRuby. Table 3 lists the parameters and settings that were used for all flow cytometry and fluorescence-activated cell sorting (FACS) experiments. We recorded 10,000 cells each from 8 independent biological replicates per strain for our analyses of BY, RM, and reporter control strains.

**Table 3:** *Flow cytometry and FACS settings.*

| Parameter | Laser Line (nm) | Laser Setting (V) | Filter |
| --- | --- | --- | --- |
| forward scatter (FSC) | 488 | 500 | 488/10 |
| side scatter (SSC) | 488 | 275 | 488/10 |
| sfGFP | 488 | 500 | 525/50 |
| mCherry | 561 | 615 | 610/20 |
| mRuby | 561 | 615 | 610/20 |

We analyzed flow cytometry data using R (R Foundation for Statistical Computing, Vienna Austria) and the flowCore R package^99^. We first filtered each flow cytometry dataset to include only those cells within 10% *±* the forward scatter (a proxy for cell size) median. We empirically determined that this gating approach captured the central peak of cells in the FSC histogram. It also removed cellular debris, aggregates of multiple cells, and restricted our analyses to cells of the same approximate size. We observed that the TFT’s output changed with the passage of time during flow cytometry experiments. We used the residuals of a loess regression of the TFT’s output on time to correct for this effect, similar to a previously-described approach^44^.

To characterize our TFT reporters, we used the following analysis steps. We extracted the median -log_2_ RFP / GFP ratio from each of 10,000 cells per strain per reporter. These values were Z-score normalized relative to the sample lowest degradation rate (typically the E3 ligase deletion strain). Following this transformation, the strain with lowest degradation rate has a degradation rate of approximately 0 and the now-scaled RFP/GFP ratio is directly proportional to the construct’s degradation rate. To compare degradation rates between strains and individual UPS activity reporters, we then converted scaled RFP/GFP ratios to Z scores, which we report as “Normalized UPS Activity”. Statistical significance was assessed using a one-way ANOVA with Tukey’s HSD post-hoc test.

For fine-mapping causal genes and variants for UPS activity QTLs, we used the following approach. We extracted the median -log_2_ RFP / GFP ratio from each of 10,000 cells per strain per reporter. These values were Z-score normalized relative to the median of the control strain (a BY strain engineered to contain the BY allele of a candidate causal gene). Statistical significance was assessed using a t-test of each experimental strain versus the control strain with Benjamini-Hochberg correction for multiple testing^100^.

### Fluorescence-Activated Cell Sorting

We selected populations of segregants for QTL mapping using a previously-described approach for isolating phenotypically extreme cell populations by FACS^43, 44^. Segregant populations were thawed approximately 16 hours prior to cell sorting and grown overnight in 5 mL of SGA medium containing G418 and canavanine. The following morning, 1 mL of cells from each segregant population was diluted into a fresh 4 mL of SGA medium containing G418 and canavanine. Segregant cultures were then grown for an additional 4 hours prior to sorting. All FACS experiments were carried out using a FACSAria II cell sorter (BD). We used plots of side scatter (SSC) height by SSC width and forward scatter (FSC) height by FSC width to remove doublets from each sample. We then filtered cells on the basis of FSC area, restricting our sorts to *±* 7.5% of the central FSC peak, which we empirically determined excluded cellular debris and aggregates while encompassing the primary haploid cell population. Finally, we defined a fluorescence-positive population by comparing each segregant population to negative control BY and RM strains without TFTs. We collected pools of 20,000 cells each from three gates drawn on each segregant population:

1. The 2% lower tail of the UPS activity distribution
2. The 2% upper tail of the UPS activity distribution
3. Fluorescence-positive cells without selection on UPS activity (“null pools”), which were used to determine the false positive rate of the QTL mapping method (see below)

We collected cell pools from two independent biological replicates (spore preparations) for each reporter. Each pool of 20,000 cells was collected into sterile 1.5 mL polypropylene tubes containing 1 mL of SGA medium and grown overnight at 30 °C with rolling. The next day, we mixed 750 *µ*L of cells with 450 *µ*L of 40% (v/v) glycerol and stored this mixture in 2 mL 96 well plates at *−*80 °C.

### Genomic DNA Isolation and Library Preparation

We extracted genomic DNA from sorted segregant pools for whole-genome sequencing. Deep-well plates containing glycerol stocks of sorted segregant pools were thawed and 800 *µ*L of each sample was pelleted by centrifugation at 3700 rpm for 10 minutes. We discarded the supernantant and resuspended cell pellets in 800 *µ*L of a 1 M sorbitol solution containing 0.1 M EDTA, 14.3 mM *β*-mercaptoethanol, and 500 U of Zymolyase lytic enzyme to digest cell walls prior to DNA extraction. The digestion reaction was carried out by resuspending cell pellets with mixing at 1000 rpm for 2 minutes followed by incubation for 2 hours at 37 °C. When the digestion reaction finished, we discarded the supernatant, resuspended cells in 50 *µ*L of phosphate buffered saline, and used the Quick-DNA 96 Plus kit (Zymo Research, Irvine, CA, USA) to extract genomic DNA. We followed the manufacturer’s protocol to extract genomic DNA with the following modifications. We incubated cells in a 20 mg/mL proteinase K solution overnight with incubation at 55 °C. After completing the DNA extraction protocol, we eluted DNA using 40 *µ*L of DNA elution buffer (10 mM Tris-HCl [pH 8.5], 0.1 mM EDTA). The DNA concentration for each sample was determined using the Qubit dsDNA BR assay kit (Thermo Fisher Scientific, Waltham, MA, USA) in a 96 well format using a Synergy H1 plate reader (BioTek Instruments, Winooski, VT, USA).

We used a previously-described approach to prepare libraries for short-read whole-genome sequencing on the Illumina Next-Seq platform^43, 44^. We used the Nextera DNA library kit (Illumina, San Diego, CA, USA) according to the manufacturer’s instructions with the following modifications. For the tagmentation reaction, 5 ng of genomic DNA from each sample was diluted in a master mix containing 4 *µ*L of Tagment DNA buffer, 1 *µ*L of sterile molecular biology grade water, and 5 *µ*L of Tagment DNA enzyme diluted 1:20 in Tagment DNA buffer. The tagmentation reaction was run on a SimpliAmp thermal cycler (Thermo Fisher Scientific) using the following parameters: 55 °C temperature, 20 *µ*L reaction volume, 10 minute incubation. To prepare libraries for sequencing, we added 10 *µ*L of the tagmentation reaction to a master mix containing 1 *µ*L of an Illumina i5 and i7 index primer pair mixture, 0.375 *µ*L of ExTaq polymerase (Takara Bio, Mountain View, CA, USA), 5 *µ*L of ExTaq buffer, 4 *µ*L of a dNTP mixture, and 29.625 *µ*L of sterile molecular biology grade water. We generated all 96 possible index oligo combinations using 8 i5 and 12 i7 index primers. The library amplification reaction was run on a SimpliAmp thermal cycler with the following parameters: initial denaturation at 95 °C for 30 seconds, then 17 cycles of 95 °C for 10 seconds (denaturation), 62 °C for 30 seconds (annealing), and 72 °C for 3 minutes (extension). We quantified the DNA concentration of each reaction using the Qubit dsDNA BR assay kit (Thermo Fisher Scientific) and pooled 10 *µ*L of each reaction. This pooled mixture was run on a 2% agarose gel and we extracted and purified DNA in the 400 bp to 600 bp region using the Monarch Gel Extraction Kit (NEB) according to the manufacturer’s instructions.

### Whole-Genome Sequencing

We submitted pooled, purified DNA libraries to the University of Minnesota Genomics Center (UMGC) for Illumina sequencing. Prior to sequencing, UMGC staff performed three quality control (QC) assays. Library concentration was determined using the PicoGreen dsDNA quantification reagent (Thermo Fisher Scientific) with libraries at a concentration of 1 ng/*µ*L passing QC. Library size was determined using the Tapestation electrophoresis system (Agilent Technologies, Santa Clara, CA, USA) with libraries in the range of 200 to 700 bp passing QC. Library functionality was determined using the KAPA DNA Library Quantification kit (Roche, Penzberg, Germany), with libraries with a concentration greater than 2 nM passing. All submitted libraries passed each QC assay. We submitted 7 libraries for sequencing at different times. Libraries were sequenced on a NextSeq 550 instrument (Illumina). Depending on the number of samples, we used the following output settings. For libraries with 70 or more samples (2 libraries), 75 bp paired end sequencing was performed in high-output mode to generate approximately 360 *×* 10^6^ reads. For libraries with 50 or fewer samples (5 libraries), 75 bp paired end sequencing was performed in mid-output mode to generate approximately 120 *×* 10^6^ reads. Average read coverage of the genome ranged from 9 to 35 with a median coverage of 28 across all libraries. Sequence data de-multiplexing was performed by UMGC. Data are currently being deposited into the NIH Sequence Read Archive.

### Raw Whole-Genome Sequencing Data Processing

We calculated allele frequencies from our whole-genome sequencing data using the following pipeline. We initially filtered reads to include only those reads with mapping quality scores greater than 30. We aligned the filtered reads to the *S. cerevisiae* reference genome (version sacCer3) using BWA^101^ (command: “mem -t 24”). We then used samtools^102^ to remove unaligned reads, non-uniquely aligned reads, and PCR duplicates (command: “samtools rmdup -S”). Finally, we produced vcf files containing coverage and allelic read counts at each of 18,871 high-confidence, reliable SNPs^49, 63^ (command: “samtools mpileup -vu -t INFO/AD -l”). Because the BY strain is closely related to the S288C genome reference *S. cerevisiae* strain, we considered BY alleles reference and RM alleles alternative alleles.

### QTL Mapping

We identified QTLs from sequence data following established procedures for bulk segregant analysis^43, 44, 63^. Allele counts in the vcf files generated above were provide to the MULTIPOOL algorithm^103^. MULTIPOOL computes logarithm of the odds (LOD) scores by comparing two models: (1) a model in which the high and low UPS activity pools come from one from common population and thus share the same frequency of the BY and RM allele, and (2) a model in which these pools come from two populations with two different allele frequencies, indicating the presence of a QTL. We identified QTLs as genomic regions exceeding an empirically-derived significance threshold (see below). We used MULTIPOOL with the following settings: bp per centiMorgan = 2,200, bin size = 100 bp, effective pool size = 1,000. As in previous QTL mapping in the BY/RM cross by bulk segregant analysis^43, 44^, we excluded variants with allele frequencies higher than 0.9 or lower than 0.1^43, 44^. We also used MULTIPOOL to estimate confidence intervals for each significant QTL, which we defined as a 2-LOD drop from the QTL peak position. To visualize QTLs and gauge their effects, we also computed the allele frequency differences (ΔAF) at each site between our high and low UPS activity pools. Because allele frequencies are affected by random counting noise, we used loess regression to smooth the allele frequency for each sample before computing ΔAF. We used the smoothed values to plot the ΔAF distribution along the genome and as a measure of QTL effect size.

### Null Sorts and Empirical False Discovery Rate Estimation

We used “null” segregant pools (fluorescence-positive cells with no selection on UPS activity) to empirically estimate the false discovery rate (FDR) of our QTL mapping method. Because these cells are obtained as two pools from the same null population in the same sample, any ΔAF differences between them are the result of technical noise or random variation. We permuted these null comparisons across segregant pools with the same UPS activity reporter for a total of 112 null comparisons. We define the “null QTL rate” at a given LOD threshold as the number of QTLs that exceeded the threshold in these comparisons divided by the number of null comparisons. To determine the FDR for a given LOD score, we then determined the number of QTLs for our experimental comparisons (high UPS activity versus low UPS activity). We define the “experimental QTL rate” as the number of experimental QTLs divided by the number of experimental comparisons. The FDR is thus computed as follows:

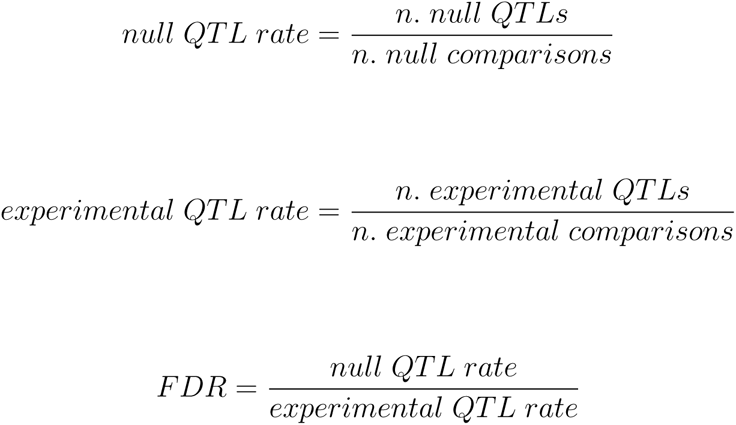

We evaluated the FDR over a LOD range of 2.5 to 10 in 0.5 LOD increments. We found that a LOD value of 4.5 led to a null QTL rate of 0.0625 and an FDR of 0.507%. We used this value as our significance threshold for QTL mapping and further filtered our QTL list by excluding QTLs that were not detected in each of two independent biological replicates. Replicating QTLs were defined as those whose peaks were within 100 kb of each other on the same chromosome with the same direction (positive or negative) of allele frequency difference between high and low UPS activity pools.

### QTL Fine-Mapping by Allelic Engineering

We used “CRISPR-Swap”^64^, a two-step method for scarless allelic editing, to fine-map QTLs to the level of their causal genes and nucleotides. In the first step of CRISPR-Swap, a gene of interest (GOI) is deleted and replaced with a selectable marker. In the second step, cells are co-transformed with (1) a plasmid that expresses CRISPR-cas9 and a guide RNA targeting the selectable marker and (2) a repair template encoding the desired allele of the GOI.

We used CRISPR-Swap to generate BY strains harboring either RM alleles or chimeric BY/RM alleles of several genes, as described below. To do so, we first replaced the gene of interest in BY with the NatMX selectable marker by transforming a PCR product encoding the NatMX cassette with 40 bp overhangs at the 5’ and 3’ ends of the targeted gene. To generate *NatMX::GOI*Δ transformation fragments, we PCR amplified NatMX from Addgene plasmid #35121 with the primers listed in Supplementary Table 6 using Phusion Hot Start Flex DNA polymerase (NEB). The NatMX cassette was transformed into the BY strain using the methods described above and transformants were plated onto YPD medium containing clonNAT. We verified the deletion of each gene of interest from single-colony purified transformants by colony PCR (primer sequences listed in Supplementary Table 6).

We then modified the original CRISPR-Swap plasmid (PFA0055, Addgene plasmid #131774) to replace its *LEU2* selectable marker with the *HIS3* selectable marker, creating plasmid PFA0227 (Supplementary Table 7). To build PFA0277, we first digested PFA0055 with restriction enzymes BsmBI-v2 and HpaI to remove the *LEU2* selectable marker. We synthesized the *S. cerevisiae HIS3* selectable marker from plasmid pRS313^104^ with 20 base pairs of overlap to BsmBI-v2/HpaI-digested PFA0055 on both ends. We used this synthetic *HIS3* fragment and BsmBI-v2/HpaI-digested PFA0055 to create plasmid PFA0227 by isothermal assembly cloning using the HiFi Assembly Cloning Kit (NEB) according to the manufacturer’s instructions. In addition to the *HIS3* selectable marker, PFA0227 contains the cas9 gene driven by the constitutively active *TDH3* promoter and a guide RNA, gCASS5a, that directs cleavage of a site immediately upstream of the *TEF* promoter used to drive expression of the MX series of selectable markers^64, 93^. We verified the sequence of PFA0227 by Sanger sequencing.

We used genomic DNA from BY and RM strains as a template to PCR amplify repair templates for CRISPR-Swap. Genomic DNA was extracted from BY and RM strains using the “10 minute prep” protocol^105^. We amplified full-length repair templates from RM and BY containing each GOI’s promoter, open-reading frame (ORF), and terminator using Phusion Hot Start Flex DNA polymerase (NEB). We also created chimeric repair templates containing combinations of BY and RM alleles using PCR splicing by overlap extension^106^. Table 4 lists the repair templates used for CRISPR swap. The sequence of all repair templates was verified by Sanger sequencing.

**Table 4:** UBR1 *CRISPR-Swap repair templates.*

| Gene | Allele Name | Promoter | ORF | Terminator |
| --- | --- | --- | --- | --- |
| *UBR1* | *UBR1* BY | BY | BY | BY |
| *UBR1* | *UBR1* RM | RM | RM | RM |
| *UBR1* | *UBR1* RM promoter | RM | BY | BY |
| *UBR1* | *UBR1* RM ORF | BY | RM | BY |
| *UBR1* | *UBR1* RM terminator | BY | BY | RM |
| *UBR1* | *UBR1* -469A>T | -469, RM; all other, BY | BY | BY |
| *UBR1* | *UBR1* -197T>G | -197, RM; all other, BY | BY | BY |
| *DOA10* | *DOA10* BY | BY | BY | BY |
| *DOA10* | *DOA10* RM | RM | RM | RM |
| *DOA10* | *DOA10* Q410E | BY | 1228, RM; all other, BY | BY |
| *DOA10* | *DOA10* K1012N | BY | 3036, RM; all other, BY | BY |
| *DOA10* | *DOA10* Y1186F | BY | 3557, RM; all other, BY | BY |
| *NTA1* | *NTA1* BY | BY | BY | BY |
| *NTA1* | *NTA1* RM | RM | RM | RM |
| *NTA1* | *NTA1* RM promoter | RM | BY | BY |
| *NTA1* | *NTA1* D111E | RM | 331, RM; all other, BY | BY |
| *NTA1* | *NTA1* E129G | RM | 386, RM; all other, BY | BY |
| *UBC6* | *UBC6* BY | BY | BY | BY |
| *UBC6* | *UBC6* RM | RM | RM | RM |
| *UBC6* | *UBC6* RM promoter | RM | BY | BY |
| *UBC6* | *UBC6* D229G | BY | 1686, RM; all other, BY | BY |
| *UBC6* | *UBC6* RM terminator | BY | BY | RM |

To create allele swap strains, we co-transformed BY strains with 200 ng of plasmid PFA0227 and 1.5 *µ*g of GOI repair template. Transformants were selected and single colony purified on synthetic complete medium lacking histidine and then patched onto solid YPD medium. We tested each strain for the desired exchange of the NatMX selectable marker with a *UBR1* allele by patching strains onto solid YPD medium containing clonNAT. We then verified allelic exchange in strains lacking clonNAT resistance by colony PCR (primers listed in Supplementary Table 6). We kept 16 independently-derived biological replicates of each allele swap strain. To test the effects of each allele swap, we transformed UPS activity reporters into our allele swap strains and characterized reporter activity by flow cytometry using the methods described above.

We tested whether a QTL on chromosome V results from variation in *UBC6* using CRISPR-Swap. Deleting *UBC6* caused a large growth defect relative to the wild-type BY strain. Providing cells with multiple *UBC6* alleles, including the BY allele, did not correct the growth rate defect. We did not observe growth defects in any other fine-mapping strains.

### RNA Isolation

We isolated total RNA from 5 independent biological replicates each of the wild-type BY strain and a BY strain edited to contain the -469A*>*T RM variant in the *UBR1* promoter (hereafter “*UBR1* -469A*>*T BY”). All 10 samples were grown and harvested at the same time. BY and *UBR1* -469A*>*T BY strains were grown overnight in 5 mL of SC medium. The following day, the cultures were diluted to an OD of 0.05 in 100 mL of fresh SC medium and grown for approximately 7 hours. When the optical density (OD) of each culture was approximately 0.40, the cells were pelleted by centrifugation at 3, 000 rpm for 10 minutes. Pellets were then washed by resuspending them in 1 mL of sterile ultrapure water, followed by centrifugation at 3, 000 rpm for 3 minutes to again pellet the cells. Following this step, cell pellets were resuspended in 1 mL of ultrapure water and split into 4 aliquots, each containing 250 *µ*L. After re-centrifuging and discarding the supernatant, the pellets were snap frozen by immersion in liquid nitrogen, followed by storage at *−*80 °C. Pellets were subsequently used for RNA isolation and mass spectrometric proteomic analysis, as described below.

Total RNA was extracted from frozen cell pellets using the ZR Fungal/Bacterial miniprep kit (Zymo), according to the manufacturer’s instructions. Briefly, total RNA was isolated from cell pellets in two batches, each containing equal numbers of BY and *UBR1* -469A*>*T BY samples. After thawing, pellets were resuspended in lysis buffer and transferred to screwcap lysis tubes containing glass beads. Tubes were secured in a Mini-BeadBeater (BioSpec Products, Bartlesville, OK, USA) and cells were processed in 5 cycles of 2 minutes of agitation followed by 2 minutes at *−*80 °C. The cell lysate/bead mixture was centrifuged for 1 minute at 16, 000 x g and 400 *µ*L of 95% ethanol was added to the cleared supernatant followed by mixing. Samples were then spun through a binding column and on-column DNA digest was performed with DNase I (Zymo) according to the manufacturer’s instructions. Total RNA was eluted from columns using 50 *µ*L of RNase-free ultrapure water. The concentration of each sample was quantified using RiboGreen; all samples had a concentration greater than 300 ng/*µ*L. The integrity of each sample was assessed at UMGC using the Tapestation (Agilent) and an RNA ScreenTape. RNA integrity numbers ranged from 9.7 to 10.0 (where 10.0 is the maximum possible score), with a median value of 9.9. All RNA samples were stored at *−*80 °C.

### RNA-seq

We isolated mRNA from each total RNA sample using the 550 ng of total RNA input and the NEBNext Poly(A) mRNA Magnetic Isolation Module (NEB). All samples were processed in a single batch and the isolated mRNA from each sample was used to prepare RNA sequencing libraries using the NEBNext Ultra II Directional RNA Library Prep kit (NEB) according to the manufacturer’s instructions. Libraries were amplified using NEBNext Ultra II Q5 polymerase and unique combinations of primers from the NEBNext Multiplex Oligos for Illumina (NEB). The following amplification protocol was used: initial denaturation at 98 °C for 30 seconds, followed by 10 cycles of 98 °C (10 seconds; denaturation), 65 °C (75 seconds; annealing and extension), and a 65 °C final extension for 5 minutes. PCR reactions were pooled using equal amounts of DNA and submitted to UMGC for three quality control assays, which measured the library concentration by PicoGreen, library functionality by KAPA qPCR, and library size using the Tapestation electrophoresis system (Agilent). The resulting library contained a small amount of adapter dimer (approximately 9%), which was subsequently removed via a bead-based cleanup. The final, cleaned library passed all three QC assays and was sequenced on a Next-Seq 2000 instrument (Illumina) in paired-end mode with 150 bp reads. The sequencing run generated 1,367,252,076 reads with an average of 136,725,207 (range: 112,285,619 to 152,571,763) reads per sample.

### RNA-seq Data Processing and Analysis

We performed quality control and preprocessing of RNA-seq data using fastp^107^. Our initial processing removed reads with a length less than 36 bp and any reads where the mean quality dropped below a mean quality score of 15 in a 4 bp window. We also used fastp to trim adapter sequences from the ends of all reads. We then used Kallisto^108^ to pseudoalign processed reads to the *S. cerevisiae* transcriptome, which was obtained from Ensembl (version 96)^109^.

To identify differentially expressed transcripts, we used the estimated counts obtained from Kallisto as a measure of gene expression and filtered the estimated counts using the following procedures. First, we computed a transcript Transcript Integrity Number (TIN) for each gene using the RSeqQC^110^ and removed any genes with a TIN less than 1 for any sample. We also removed any genes that Kallisto estimated to have an effective length less of less than 1 and those genes whose estimated counts were less than 10 in any sample. The resuling dataset comprised 5,676 expressed genes. Raw RNA-seq reads and processed counts are in the process of being deposited into the Gene Expression Omnibus repository. We used DESeq2^111^ to perform statistical analysis of the resulting dataset. We used the RNA harvest batch and OD at time of sample harvest as covariates in our analysis. To further control for confounding sample-to-sample variation, we used surrogate variable analysis^112, 113^, which identified two significant surrogate variables that were subsequently added to our statistical model. We corrected for multiple testing using the Benjamini-Hochberg method^100^ and considered significant differences as those with a corrected *p*-value less than 0.1.

To link differences in transcript abundance to biological pathways, we performed gene ontology enrichment analysis using PANTHER^114^. The “statistical overrepresentation test” was used to search for gene ontology (GO) biological processes and Reactome pathways enriched in our set of 78 transcripts differentially expressed between BY and “*UBR1* -469A*>*T” BY. We used the 5,676 genes quantified in our RNA-seq statistical analysis as the reference set and used the Benjamini-Hochberg method^100^ to correct for multiple testing. GO terms and Reactome pathways with a corrected *p*-value less than 0.05 were considered significant in our analysis.

### Protein Isolation and Proteomic Analysis by Mass Spectrometry

To quantify gene expression at the protein level, we submitted five cell pellets each from the same BY and -469A*>*T BY cultures used for RNA-seq analysis to the University of Minnesota Center for Mass Spectrometry and Proteomics (CMSP) for proteomic analysis by mass spectrometry. Cell pellets were resuspended for protein extraction in a protein extraction buffer containing 7 M urea, 2 M thiourea, 0.4 M triethylammonium bicarbonate pH 8.5, 20% acetonitrile, and 4 mM tris(2-carboxyethyl)phosphine. Cell lysis and protein extraction was then performed using the Barocycler NEP2320 (Pressure Biosciences, Medford, MA, USA).

Samples were prepared and analyzed by mass spectrometry as follows. CMSP first labeled individual samples using the tandem mass tag (TMT) 10plex labeling kit (Thermo). After tagging, samples were pooled for analysis by mass spectrometry on an Orbitrap Tribrid Eclipse instrument (Thermo). Database searching was performed using the Proteome Discoverer software and the statistical analysis of protein abundance was performed in Scaffold (Proteome Software, Portland, OR, USA). We considered proteins to be differentially abundant between strains if they had a and a Benjamini-Hochberg corrected *p*-value less than 0.1. We performed ontological enrichment analysis of differentially abundant proteins using PANTHER as described above, except that the set of 3,774 detected proteins was used as the reference set.

### Evolutionary Analysis of Variants

We inferred the allelic status of individual variants by comparing them to two outgroups: a likely-ancestral Taiwanese *S. cerevisiae* isolate and the sister species *Saccharomyces paradoxus*. We classified variants as ancestral if they were found in at least one outgroup. All alleles analyzed in this study could be unambiguously classified using this approach. We extracted the population frequency of all analyzed variants using genome sequence data from a panel of 1,011 *S. cerevisiae* isolates^66^.

### Data and Statisical Analysis

All data were analyzed using R (version 3.6.1; R Project for Statistical Computing). For all boxplots, the center line shows the median, the box excludes the upper and lower quartiles, the whiskers extend to 1.5 times the interquartile range. Protein structure predictions were obtained from the AlphaFold Protein Structure Database^115^ and visualized using ChimeraX^116^. DNA binding motifs were determined using the Yeast Transcription Factor Specificity Compendium database^117^. Final figures and illustrations were made using Inkscape (version 0.92; Inkscape Project).

Computational scripts used to process data, for statistical analysis, and to generate figures are available at http://www.github.com/mac230/N-end_Rule_QTL_paper.

## Supporting information

All N-degron QTL Mapping Traces

## Acknowledgments

We thank Leonid Kruglyak for the BY and RM yeast strains and Michael Knop for technical assistance in implementing the TFT reporter system. We thank the University of Minnesota’s Flow Cytometry Resource, Genomics Center, and Center for Mass Spectrometry and Proteomics for their contributions to the project. We thank Margaret Kliebhan for the BY / RM variant file used for QTL mapping. We thank the members of the Albert laboratory and the BioKansas Scientific Writing Program for critical feedback on the manuscript.

## Author Contributions

Conceptualization: MAC, FWA

Methodology: MAC, GM, FWA

Formal Analysis: MAC

Investigation: MAC, GM

Funding Acquisition: MAC, FWA

Resources: FWA

Supervision: MAC, FWA

Visualization: MAC

Writing - Original Draft: MAC

Writing - Review and Editing: MAC, FWA

## Supplementary Text

### Supplementary Note 1 - Genetic Influences on the Proline N-Degron

We observed that, consistent with previous results^71, 118^, the proline N-degron TFT was only partially stabilized in BY *DOA10*Δ (Figure 1D, Supplementary Figure 1, Supplementary Table 1). Ubiquitin is inefficiently cleaved in the ubiquitin-fusion technique when followed by a proline N-degron^1, 57^, leading to the production of two species, proline N-degron constructs and constructs with uncleaved N-terminal ubiquitin moieties. N-terminal ubiquitin functions as a degron and is recognized and degraded by the ubiquitin-fusion degradation (UFD) UPS pathway^119^. The proline N-end TFT thus measures the activities of the Ac/N-end pathway towards the proline N-degron and the UFD pathway towards the N-terminal ubiquitin fusion degron.

Despite this partial limitation, we were able to map genetic influences on the proline N-degron. We detected 5 Ac/N-end pathway-specific QTLs using our proline reporter, including those resulting from variation in *DOA10* and *UBC6*. There were no additional QTLs affecting the majority of Ac/N-end reporters that were not detected with the proline reporter (Figure 2B, Supplementary Table 2). We therefore conclude that the QTLs identified with the proline reporter correspond primarily to true genetic influences on the proline N-degron.

Three QTLs, on one chromosome XII and two on chromosome XVI were detected only with the proline reporter (Figure 2B, Supplementary Table 2). The intervals for these QTLs did not contain genes previously linked to either the proline N-degron, the Ac/N-end pathway, or the UFD pathway.

### Supplementary Note 2 - Global Analysis of Arg/N-end and Ac/N-end QTLs

We evaluated the extent to which individual QTLs were shared across pathways and reporters. To do so, we divided each chromosome into adjacent 100 kb bins. We considered a QTL to be shared between reporters if the peak position for two or more QTLs from different reporters were within the same bin. We observed that QTLs were often unique to an individual N-degron (Figure 2B, Supplementary Figure 2), highlighting the complexity of genetic influences on N-end rule pathways. There was relatively little sharing of QTLs within the Arg/N-end pathway, for which only one QTL affected the majority of reporters (Figure 2B, Supplementary Figure 2). By contrast, the Ac/N-end reporters tended to share more of their QTLs (Figure 2B, Supplementary Figure 2). In particular, a QTL on chromosome XII affected all Ac/N-end reporters and Ac/N-end-specific QTLs on chromosomes I, V, and VII affected 7 of 8 Ac/N-end reporters (Figure 2B). These results suggest that genetic influences on the degradation of Ac/N-end rule substrates may affect this pathway more broadly than those of the Arg/N-end pathway. This notion is consistent with the molecular mechanisms that generate Ac/N-degrons, which share several molecular events, including co-translational excision of methionine residues and acetylation of the N-degron^1, 71^. By contrast, the mechanisms that generate Arg/N-degrons are less general^1^, consistent with the largely N-degron-specific QTL architectures we observe for this pathway. These results thus provide additional evidence that UPS pathways are shaped by distinct, complex genetic architectures.

## Supplementary Figures

**Supplementary Figure 1:**
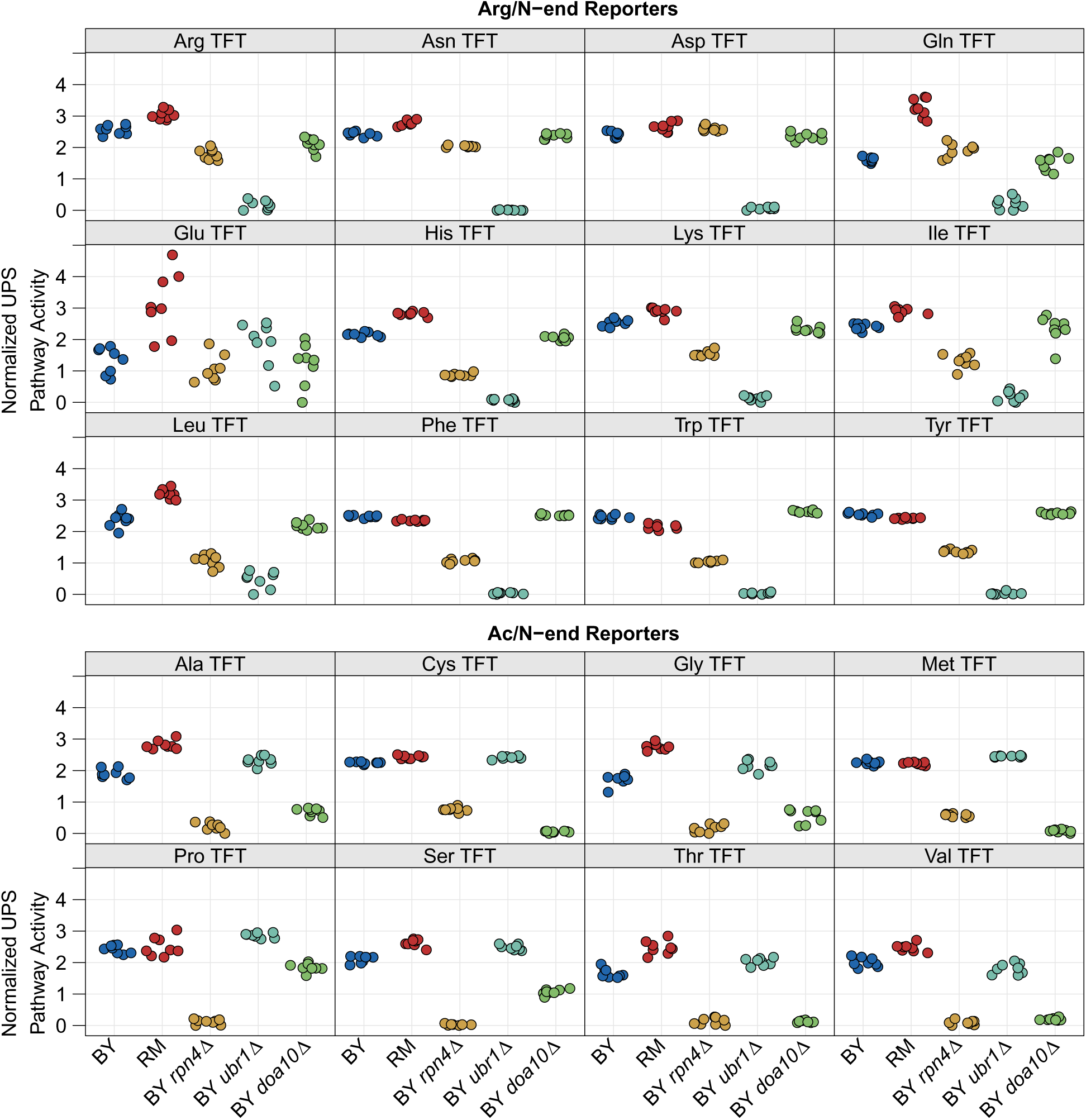
*Comparison of UPS activity between strains across N-degron reporters. The -log_2_ RFP / GFP ratio value was extracted from 10,000 cells from each of 8 independent biological replicates per strain per reporter and converted to Z-scores. High values correspond to high UPS activity and low values correspond to low UPS activity. Tukey HSD* p *values for each between strain comparison for each reporter are listed in Supplementary Table 1.*

**Supplementary Figure 2:**
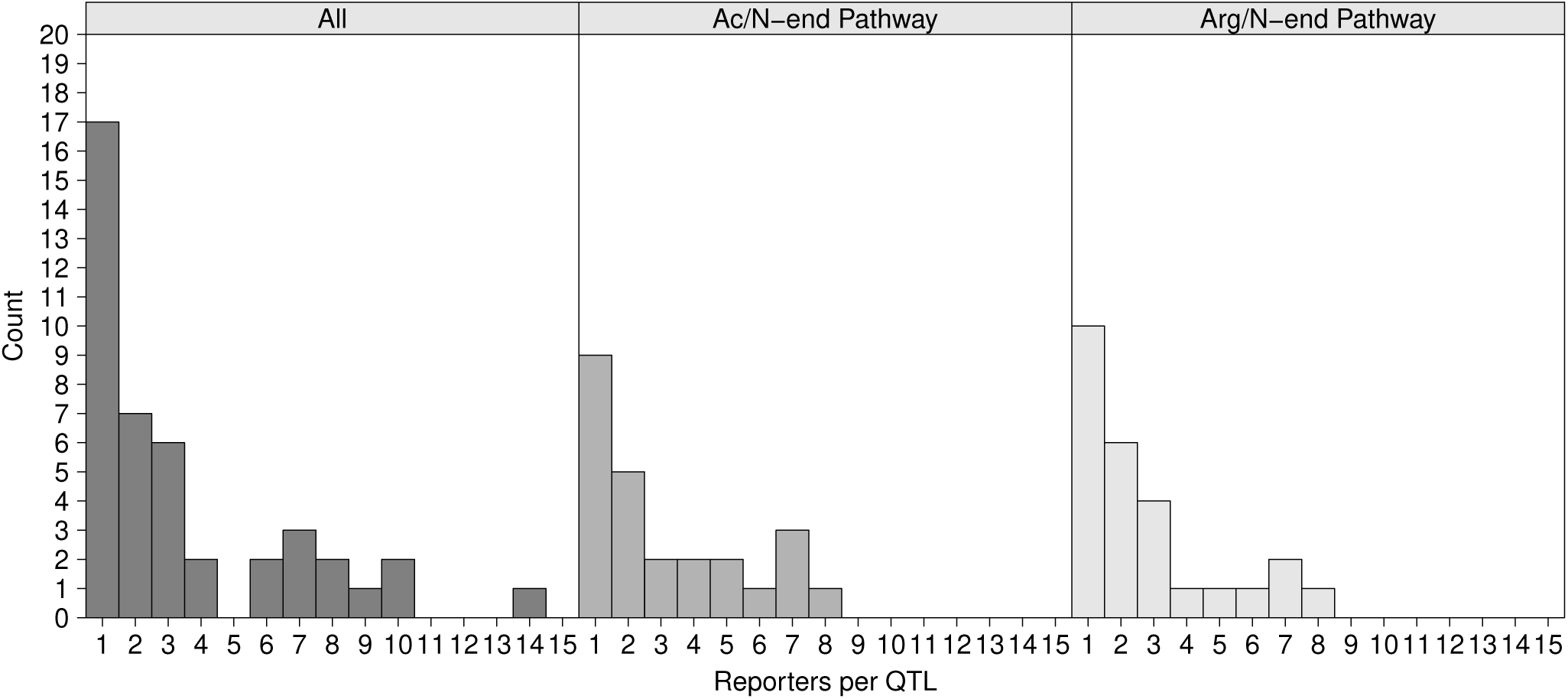
*Pathway and degron specificity of UPS activity QTLs. The histograms show the number of N-degrons affected by individual QTLs for all N-degrons (left), Ac/N-degrons (middle), and Arg/N-degrons (right).*

**Supplementary Figure 3:**
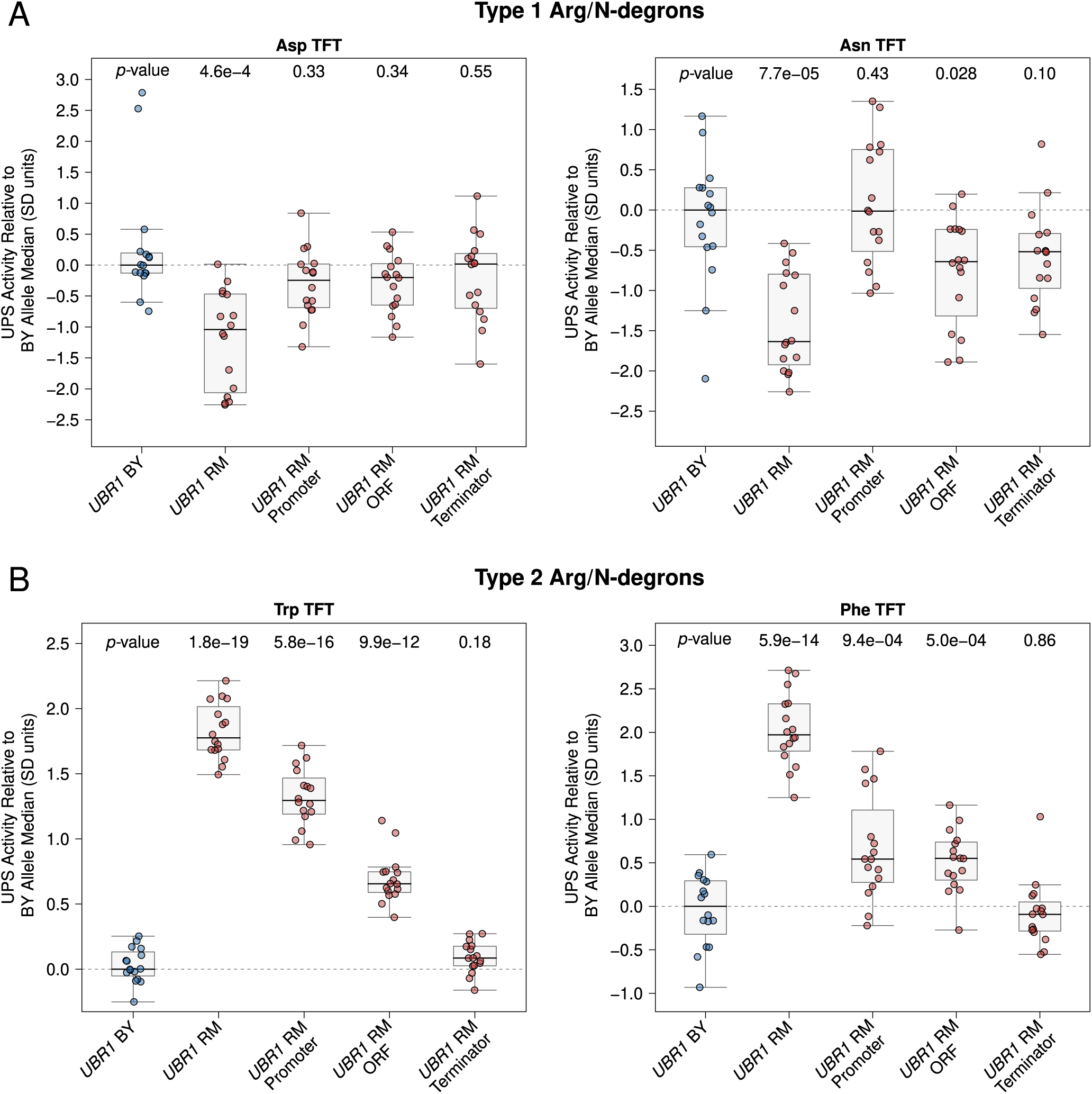
UBR1 *full gene fine-mapping results. The BY strain was engineered to contain full or partial RM* UBR1 *alleles as indicated and UPS activity towards the indicated Type 1 and Type 2 Arg/N-degron TFTs was measured by flow cytometry. UPS activity was Z-score normalized and scaled relative to the median of a control BY strain engineered to contain the full BY* UBR1 *allele. Each point in the plot shows the median of 10,000 cells for each of 16 independent biological replicates per strain per reporter.* p*-values at the top of the plot display the Benjamini-Hochberg-corrected* p *value for the t-test of the indicated strain versus the strain with the BY* UBR1 *allele. Box plot center lines, box boundaries, and whiskers display the median, interquartile range, and 1.5 times the interquartile range, respectively. A. UPS activity towards the indicated Type 1 Arg/N-degrons. B. UPS activity towards the indicated Type 2 Arg/N-degrons.*

**Supplementary Figure 4:**
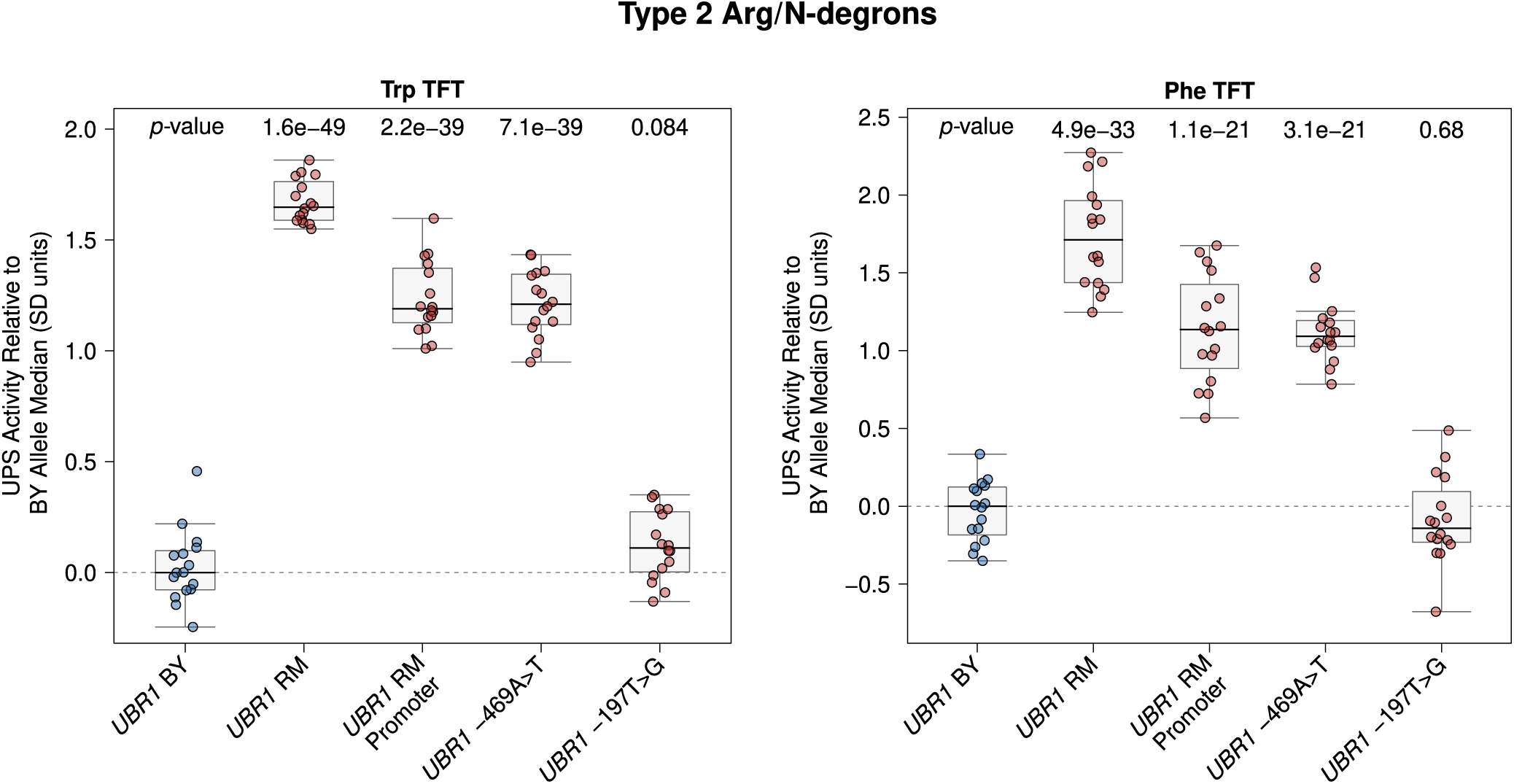
*Fine-mapping the causal nucleotide in the* UBR1 *promoter. The BY strain was engineered to carry the RM* UBR1 *promoter or one of the two single nucleotide BY / RM* UBR1 *promoter variants as indicated and UPS activity towards the Trp and Phe Type 2 Arg/N-degrons was measured by flow cytometry. UPS activity was Z-score normalized and scaled relative to the median of a control BY strain engineered to contain the full BY* UBR1 *allele. Each point in the plot shows the median of 10,000 cells for each of 16 independent biological replicates per strain per reporter.* p*-values at the top of the plot display the Benjamini-Hochberg-corrected* p *value for the t-test of the indicated strain versus the strain with the BY* UBR1 *allele. Box plot center lines, box boundaries, and whiskers display the median, interquartile range, and 1.5 times the interquartile range, respectively.*

**Supplementary Figure 5:**
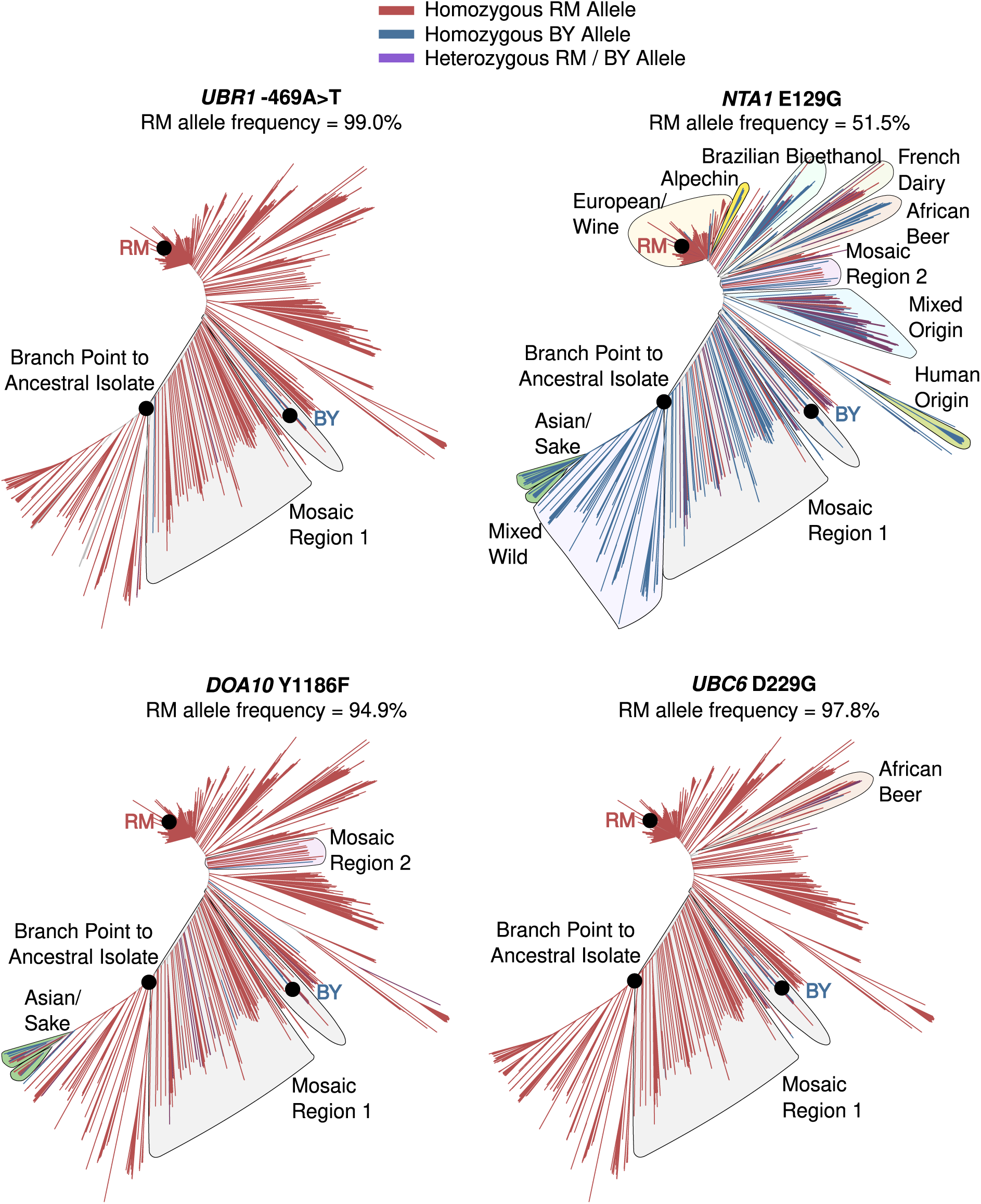
*Population frequencies and distribution of causal variants. Tree diagrams show genetic distance among a global panel of* S. cerevisiae *isolates with branches colored according to which allele a strain carries. Indicated clades with the BY allele for a causal DNA variant are outlined.*

**Supplementary Figure 6:**
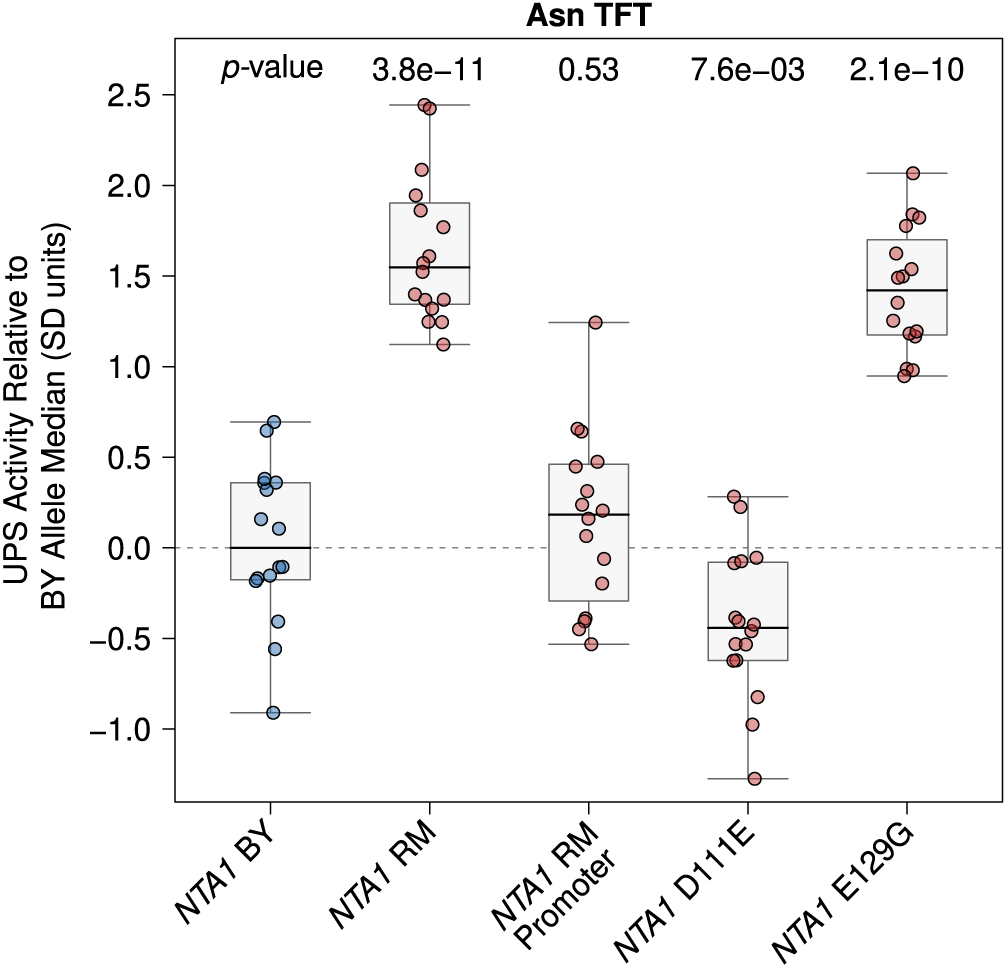
*Full* NTA1 *fine-mapping results. The BY strain was engineered to contain full or partial RM* NTA1 *alleles as indicated and UPS activity towards the Asn Arg/N-degron was measured by flow cytometry. UPS activity was Z-score normalized and scaled relative to the median of a control BY strain engineered to contain the full BY* NTA1 *allele. Each point in the plot shows the median of 10,000 cells for each of 16 independent biological replicates per strain per reporter.* p*-values at the top of the plot display the Benjamini-Hochberg-corrected* p *value for the t-test of the indicated strain versus the strain with the BY* UBR1 *allele. Box plot center lines, box boundaries, and whiskers display the median, interquartile range, and 1.5 times the interquartile range, respectively.*

**Supplementary Figure 7:**
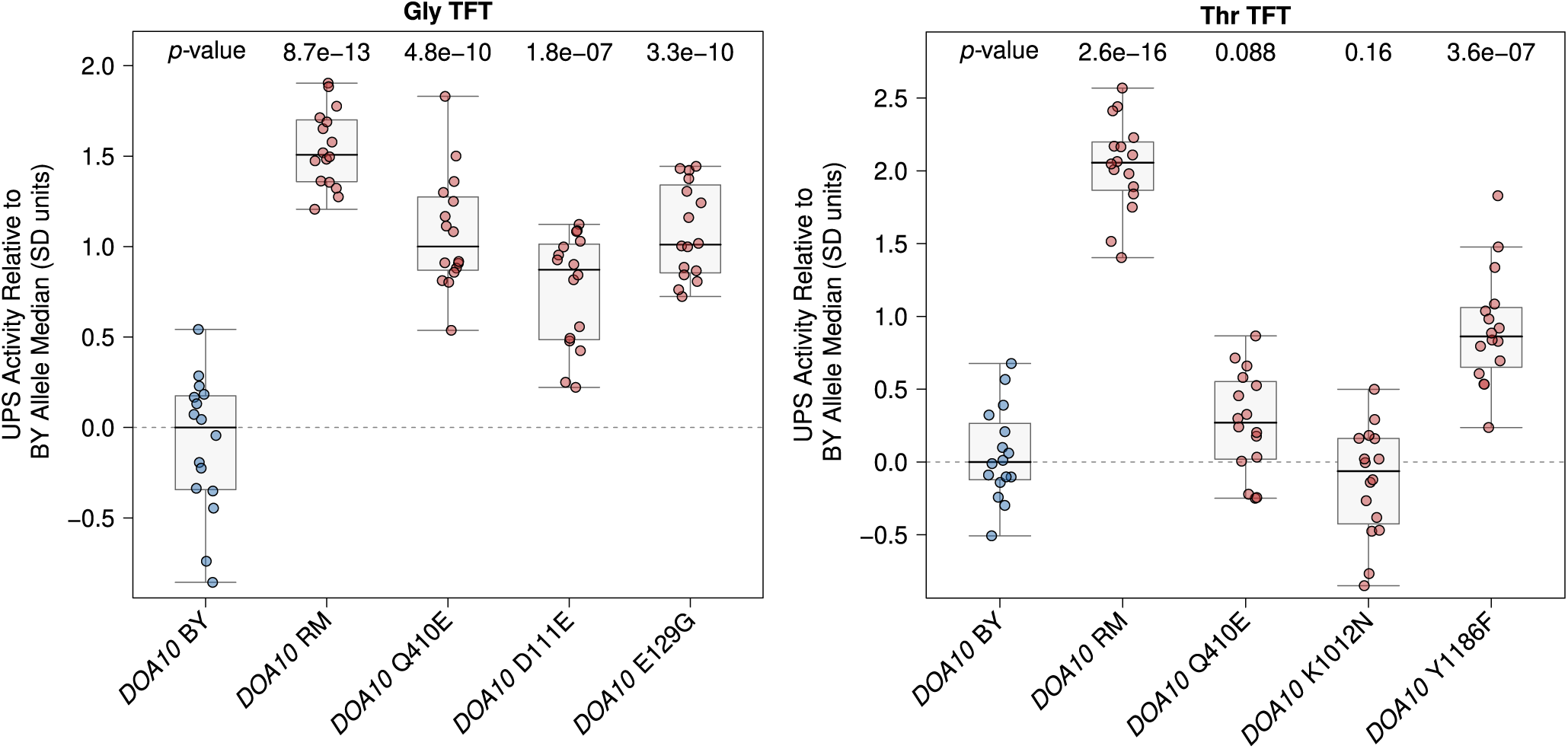
*Full* DOA10 *fine-mapping results. The BY strain was engineered to contain full or partial RM* DOA10 *alleles as indicated and UPS activity towards the Gly and Thr Ac/N-degrons was measured by flow cytometry. UPS activity was Z-score normalized and scaled relative to the median of a control BY strain engineered to contain the full BY* DOA10 *allele. Each point in the plot shows the median of 10,000 cells for each of 16 independent biological replicates per strain per reporter.* p*-values at the top of the plot display the Benjamini-Hochberg-corrected* p *value for the t-test of the indicated strain versus the strain with the BY* UBR1 *allele. Box plot center lines, box boundaries, and whiskers display the median, interquartile range, and 1.5 times the interquartile range, respectively.*

**Supplementary Figure 8:**
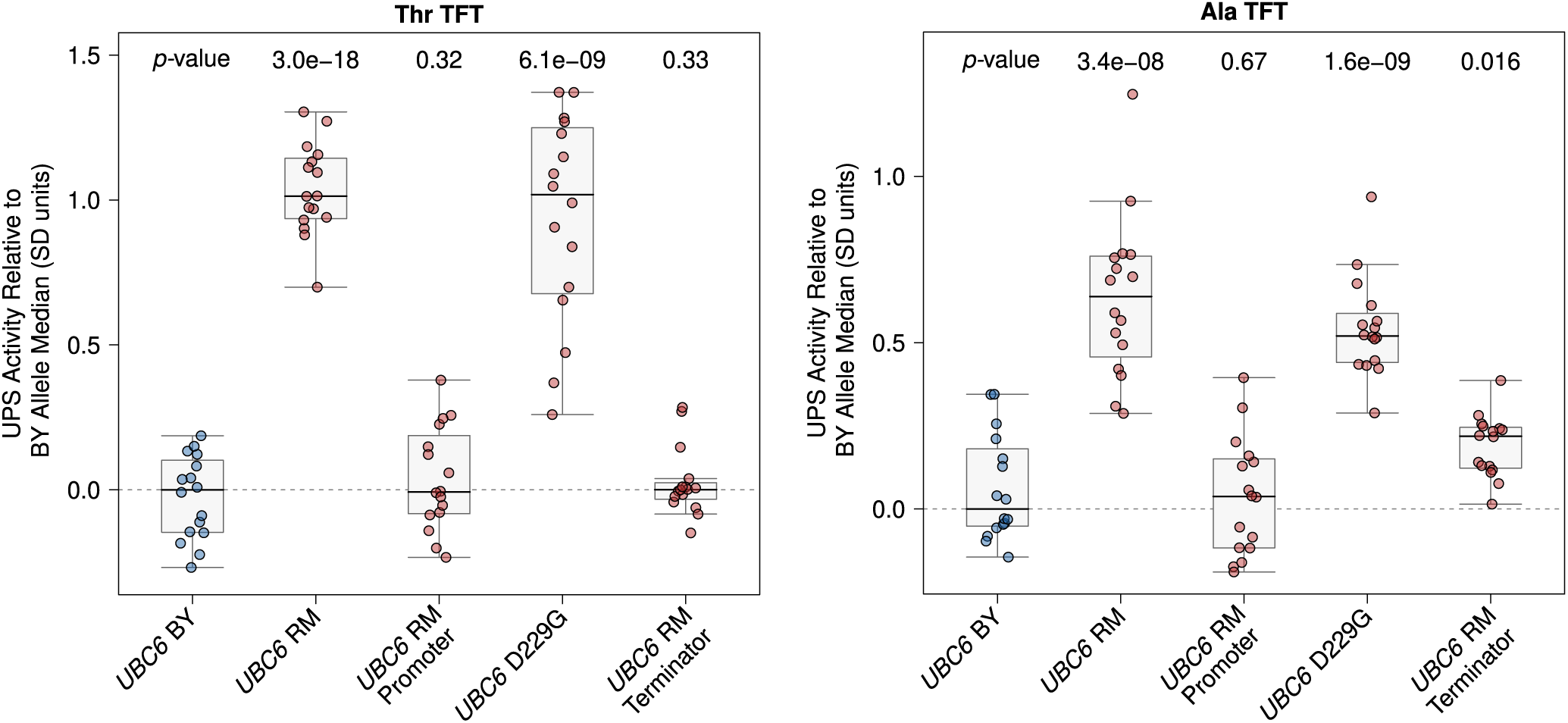
*Full* UBC6 *fine-mapping results. The BY strain was engineered to contain full or partial RM* UBC6 *alleles as indicated and UPS activity towards the Thr and Ala Ac/N-degrons was measured by flow cytometry. UPS activity was Z-score normalized and scaled relative to the median of a control BY strain engineered to contain the full BY* UBC6 *allele. Each point in the plot shows the median of 10,000 cells for each of 16 independent biological replicates per strain per reporter.* p*-values at the top of the plot display the Benjamini-Hochberg-corrected* p *value for the t-test of the indicated strain versus the strain with the BY* UBR1 *allele. Box plot center lines, box boundaries, and whiskers display the median, interquartile range, and 1.5 times the interquartile range, respectively.*

**Supplementary Figure 9:**
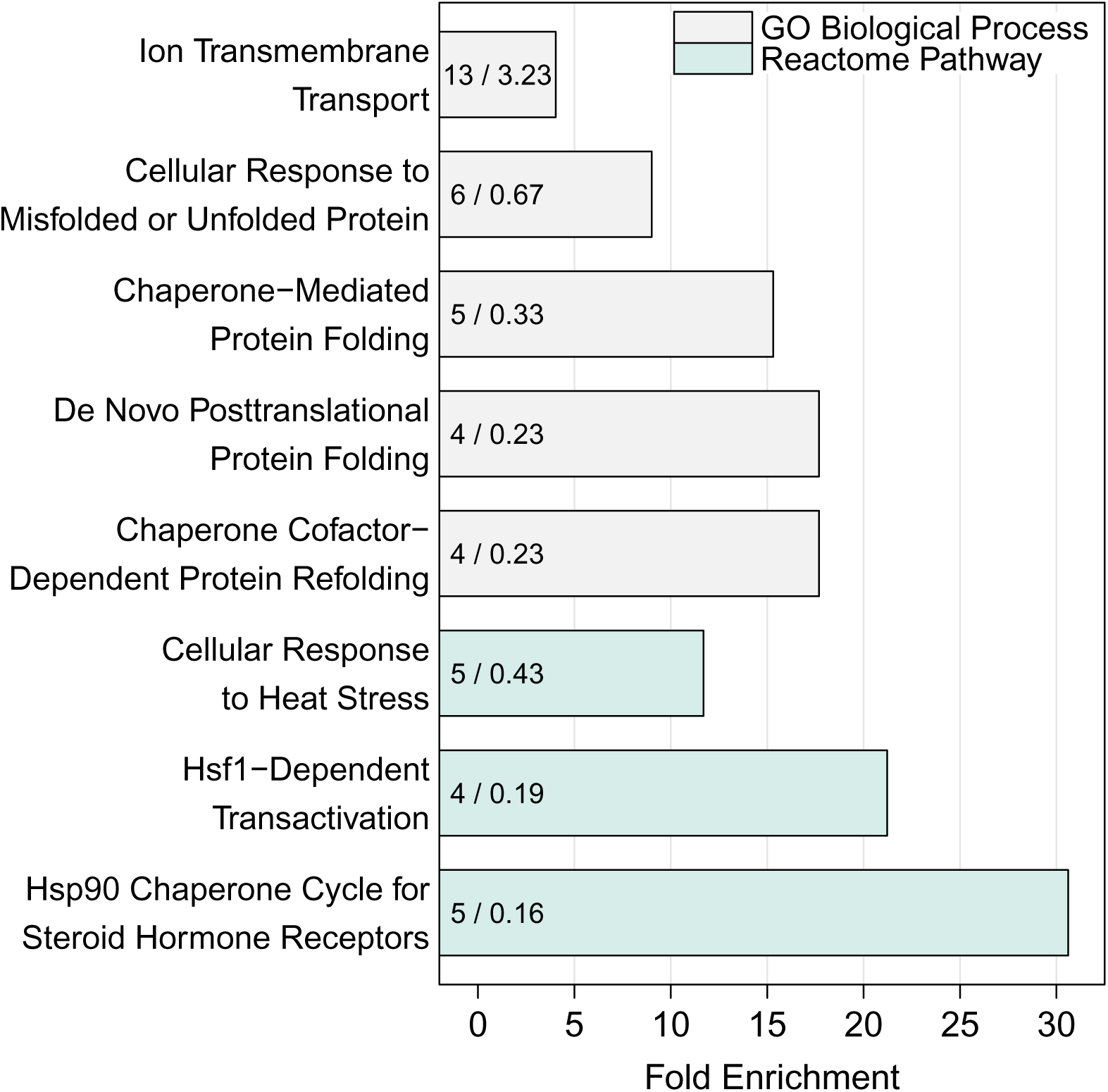
*Barchart of significantly over-represented Gene Ontology Biological Process and Reactome Pathway terms identified using the list of differentially expressed mRNA transcripts between the wild-type BY strain and BY* UBR1 *-469A>T. The numbers in the bars denote the observed (left) and expected (right) number of genes for a given process or pathway.*

**Supplementary Figure 10:**
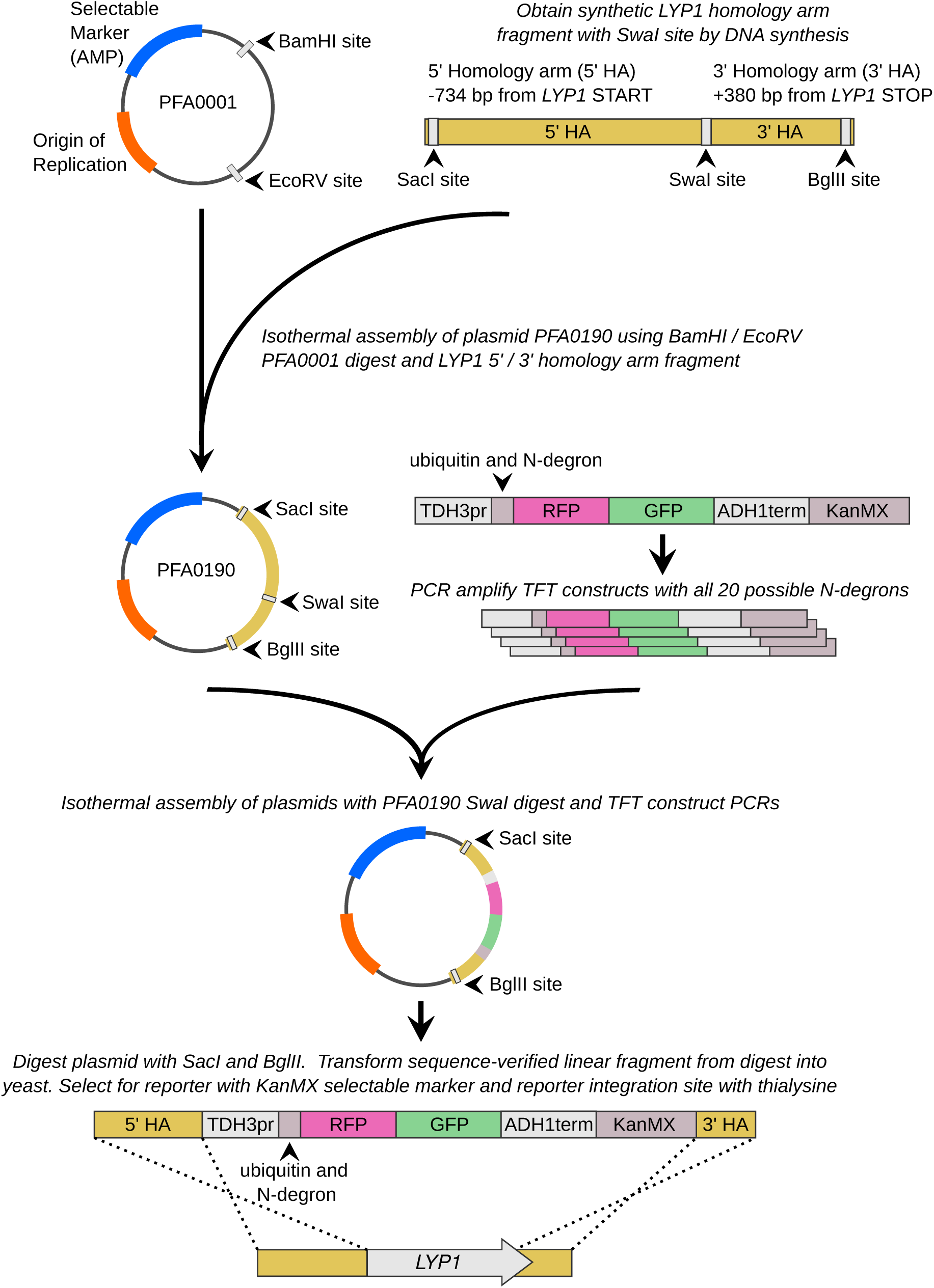
*Overview of the constructs and strain construction steps used to generate yeast strains harboring TFT UPS activity reporters.*

## References

1. A. Varshavsky. The N-end rule pathway and regulation by proteolysis. Protein Sci., 20(8):1298–1345, Aug 2011.

2. G. A. Collins and A. L. Goldberg. The Logic of the 26S Proteasome. Cell, 169(5):792–806, May 2017.

3. J. Hanna and D. Finley. A proteasome for all occasions. FEBS Lett, 581(15):2854–2861, Jun 2007.

4. O. Coux, K. Tanaka, and A. L. Goldberg. Structure and functions of the 20S and 26S proteasomes. Annu Rev Biochem, 65:801–847, 1996.

5. A. Hershko and A. Ciechanover. The ubiquitin system. Annu Rev Biochem, 67:425–479, 1998.

6. A. Bachmair, D. Finley, and A. Varshavsky. In vivo half-life of a protein is a function of its aminoterminal residue. Science, 234(4773):179–186, Oct 1986.

7. A. Ciechanover, A. Orian, and A. L. Schwartz. Ubiquitin-mediated proteolysis: biological regulation via destruction. Bioessays, 22(5):442–451, May 2000.

8. A. Varshavsky. Naming a targeting signal. Cell, 64(1):13–15, Jan 1991.

9. J. S. Bett. Proteostasis regulation by the ubiquitin system. Essays Biochem, 60(2):143–151, 10 2016.

10. D. Finley, H. D. Ulrich, T. Sommer, and P. Kaiser. The ubiquitin-proteasome system of Saccharomyces cerevisiae. Genetics, 192(2):319–360, Oct 2012.

11. A. F. Kisselev, T. N. Akopian, K. M. Woo, and A. L. Goldberg. The sizes of peptides generated from protein by mammalian 26 and 20 S proteasomes. Implications for understanding the degradative mechanism and antigen presentation. J Biol Chem, 274(6):3363–3371, Feb 1999.

12. C. Pohl and I. Dikic. Cellular quality control by the ubiquitin-proteasome system and autophagy. Science, 366(6467):818–822, 11 2019.

13. A. L. Schwartz and A. Ciechanover. The ubiquitin-proteasome pathway and pathogenesis of human diseases. Annu Rev Med, 50:57–74, 1999.

14. M. Schmidt and D. Finley. Regulation of proteasome activity in health and disease. Biochim Biophys Acta, 1843(1):13–25, Jan 2014.

15. E. M. Sontag, W. I. M. Vonk, and J. Frydman. Sorting out the trash: the spatial nature of eukaryotic protein quality control. Curr Opin Cell Biol, 26:139–146, Feb 2014.

16. S. Grimm, A. Höhn, and T. Grune. Oxidative protein damage and the proteasome. Amino Acids, 42(1):23–38, Jan 2012.

17. D. Finley and M. A. Prado. The Proteasome and Its Network: Engineering for Adaptability. Cold Spring Harb Perspect Biol, 12(1), 01 2020.

18. R. S. Marshall and R. D. Vierstra. Proteasome storage granules protect proteasomes from autophagic degradation upon carbon starvation. Elife, 7, 04 2018.

19. D. Laporte, B. Salin, B. Daignan-Fornier, and I. Sagot. Reversible cytoplasmic localization of the proteasome in quiescent yeast cells. J Cell Biol, 181(5):737–745, Jun 2008.

20. M. Bajorek, D. Finley, and M. H. Glickman. Proteasome disassembly and downregulation is correlated with viability during stationary phase. Curr Biol, 13(13):1140–1144, Jul 2003.

21. A. Stolzing and T. Grune. The proteasome and its function in the ageing process. Clin Exp Dermatol, 26(7):566–572, Oct 2001.

22. M. A. Baraibar and B. Friguet. Changes of the proteasomal system during the aging process. Prog Mol Biol Transl Sci, 109:249–275, 2012.

23. R. Shringarpure and K. J. Davies. Protein turnover by the proteasome in aging and disease. Free Radic Biol Med, 32(11):1084–1089, Jun 2002.

24. N. P. Dantuma and L. C. Bott. The ubiquitin-proteasome system in neurodegenerative diseases: pre-cipitating factor, yet part of the solution. Front Mol Neurosci, 7:70, 2014.

25. J. M. Deger, J. E. Gerson, and R. Kayed. The interrelationship of proteasome impairment and oligomeric intermediates in neurodegeneration. Aging Cell, 14(5):715–724, Oct 2015.

26. L. Petrucelli and T. M. Dawson. Mechanism of neurodegenerative disease: role of the ubiquitin pro-teasome system. Ann Med, 36(4):315–320, 2004.

27. C. Zheng, T. Geetha, and J. R. Babu. Failure of ubiquitin proteasome system: risk for neurodegenerative diseases. Neurodegener Dis, 14(4):161–175, 2014.

28. M. Zenker, J. Mayerle, A. Reis, and M. M. Lerch. Genetic basis and pancreatic biology of Johanson-Blizzard syndrome. Endocrinol Metab Clin North Am, 35(2):243–253, Jun 2006.

29. M. Zenker, J. Mayerle, M. M. Lerch, A. Tagariello, K. Zerres, P. R. Durie, M. Beier, G. Hülskamp, C. Guzman, H. Rehder, F. A. Beemer, B. Hamel, P. Vanlieferinghen, R. Gershoni-Baruch, M. W. Vieira, M. Dumic, R. Auslender, V. L. Gil-da Silva-Lopes, S. Steinlicht, M. Rauh, S. A. Shalev, C. Thiel, A. B. Ekici, A. Winterpacht, Y. T. Kwon, A. Varshavsky, and A. Reis. Deficiency of UBR1, a ubiquitin ligase of the N-end rule pathway, causes pancreatic dysfunction, malformations and mental retardation (Johanson-Blizzard syndrome). Nat Genet, 37(12):1345–1350, Dec 2005.

30. A. Brehm, Y. Liu, A. Sheikh, B. Marrero, E. Omoyinmi, Q. Zhou, G. Montealegre, A. Biancotto, A. Reinhardt, A. Almeida de Jesus, M. Pelletier, W. L. Tsai, E. F. Remmers, L. Kardava, S. Hill, H. Kim, H. J. Lachmann, A. Megarbane, J. J. Chae, J. Brady, R. D. Castillo, D. Brown, A. V. Casano, L. Gao, D. Chapelle, Y. Huang, D. Stone, Y. Chen, F. Sotzny, C. C. Lee, D. L. Kastner, A. Torrelo, A. Zlotogorski, S. Moir, M. Gadina, P. McCoy, R. Wesley, K. I. Rother, K. Rother, P. W. Hildebrand, P. Brogan, E. Krüger, I. Aksentijevich, and R. Goldbach-Mansky. Additive loss-of-function proteasome subunit mutations in CANDLE/PRAAS patients promote type I IFN production. J Clin Invest, 125(11):4196–4211, Nov 2015.

31. K. Arima, A. Kinoshita, H. Mishima, N. Kanazawa, T. Kaneko, T. Mizushima, K. Ichinose, H. Nakamura, A. Tsujino, A. Kawakami, M. Matsunaka, S. Kasagi, S. Kawano, S. Kumagai, K. Ohmura, T. Mimori, M. Hirano, S. Ueno, K. Tanaka, M. Tanaka, I. Toyoshima, H. Sugino, A. Yamakawa, K. Tanaka, N. Niikawa, F. Furukawa, S. Murata, K. Eguchi, H. Ida, and K. Yoshiura. Proteasome assembly defect due to a proteasome subunit beta type 8 (PSMB8) mutation causes the autoinflammatory disorder, Nakajo-Nishimura syndrome. Proc Natl Acad Sci U S A, 108(36):14914–14919, Sep 2011.

32. Y. Liu, Y. Ramot, A. Torrelo, A. S. Paller, N. Si, S. Babay, P. W. Kim, A. Sheikh, C. C. Lee, Y. Chen, A. Vera, X. Zhang, R. Goldbach-Mansky, and A. Zlotogorski. Mutations in proteasome subunit B type 8 cause chronic atypical neutrophilic dermatosis with lipodystrophy and elevated temperature with evidence of genetic and phenotypic heterogeneity. Arthritis Rheum, 64(3):895–907, Mar 2012.

33. K. Xia, H. Guo, Z. Hu, G. Xun, L. Zuo, Y. Peng, K. Wang, Y. He, Z. Xiong, L. Sun, Q. Pan, Z. Long, X. Zou, X. Li, W. Li, X. Xu, L. Lu, Y. Liu, Y. Hu, D. Tian, L. Long, J. Ou, Y. Liu, X. Li, L. Zhang, Y. Pan, J. Chen, H. Peng, Q. Liu, X. Luo, W. Su, L. Wu, D. Liang, H. Dai, X. Yan, Y. Feng, B. Tang, J. Li, Z. Miedzybrodzka, J. Xia, Z. Zhang, X. Luo, X. Zhang, D. St Clair, J. Zhao, and F. Zhang. Common genetic variants on 1p13.2 associate with risk of autism. Mol Psychiatry, 19(11):1212–1219, Nov 2014.

34. S. J. Diskin, M. Capasso, R. W. Schnepp, K. A. Cole, E. F. Attiyeh, C. Hou, M. Diamond, E. L. Carpenter, C. Winter, H. Lee, J. Jagannathan, V. Latorre, A. Iolascon, H. Hakonarson, M. Devoto, and J. M. Maris. Common variation at 6q16 within HACE1 and LIN28B influences susceptibility to neuroblastoma. Nat Genet, 44(10):1126–1130, Oct 2012.

35. Y. S. Cho, C. H. Chen, C. Hu, J. Long, R. T. Ong, X. Sim, F. Takeuchi, Y. Wu, M. J. Go, T. Yamauchi, Y. C. Chang, S. H. Kwak, R. C. Ma, K. Yamamoto, L. S. Adair, T. Aung, Q. Cai, L. C. Chang, Y. T. Chen, Y. Gao, F. B. Hu, H. L. Kim, S. Kim, Y. J. Kim, J. J. Lee, N. R. Lee, Y. Li, J. J. Liu, W. Lu, J. Nakamura, E. Nakashima, D. P. Ng, W. T. Tay, F. J. Tsai, T. Y. Wong, M. Yokota, W. Zheng, R. Zhang, C. Wang, W. Y. So, K. Ohnaka, H. Ikegami, K. Hara, Y. M. Cho, N. H. Cho, T. J. Chang, Y. Bao, Å. K. Hedman, A. P. Morris, M. I. McCarthy, R. Takayanagi, K. S. Park, W. Jia, L. M. Chuang, J. C. Chan, S. Maeda, T. Kadowaki, J. Y. Lee, J. Y. Wu, Y. Y. Teo, E. S. Tai, X. O. Shu, K. L. Mohlke, N. Kato, B. G. Han, M. Seielstad, B. F. Voight, L. J. Scott, V. Steinthorsdottir, A. P. Morris, C. Dina, R. P. Welch, E. Zeggini, C. Huth, Y. S. Aulchenko, G. Thorleifsson, L. J. McCullch, T. Ferreira, H. Grallert, N. Amin, G. Wu, C. J. Willer, S. Raychaudhuri, S. A. McCarroll, C. Langenberg, O. M. Hofmann, J. Dupuis, L. Qi, A. V. Segrè, M. van Hoek, P. Navarro, K. Ardlie, B. Balkau, R. Benediktsson, A. J. Bennett, R. Blagieva, E. Boerwinkle, L. L. Bonnycastle, K. B. Bostöm, B. Bravenboer, S. Bumpstead, N. P. Burtt, G. Charpentier, P. S. Chines, M. Cornelis, D. J. Couper, G. Crawford, A. S. Doney, K. S. Elliott, A. L. Elliott, M. R. Erdos, C. S. Fox, C. S. Franklin, M. Ganser, C. Gieger, N. Grarup, T. Green, S. Griffin, C. J. Groves, C. Guiducci, S. Hadjadj, N. assanali, C. Herder, B. Isomaa, A. U. Jackson, P. R. Johnson, T. Jørgensen, W. H. Kao, N. Klopp, A. Kong, P. Kraft, J. Kuusisto, T. Lauritzen, M. Li, A. Lieverse, C. M. Lindgren, V. Lyssenko, M. Marre, T. Meitinger, K. Midthjell, M. A. Morken, N. Narisu, P. Nilsson, K. R. Owen, F. Payne, J. R. Perry, A. K. Petersen, C. Platou, C. Proença, I. Prokopenko, W. Rathmann, N. W. Rayner, N. R. Robertson, G. Rocheleau, M. Roden, M. J. Samp-son, R. Saxena, B. M. Shields, P. Shrader, G. Sigurdsson, T. Sparsø, K. Strassburger, H. M. Stringham, Q. Sun, A. J. Swift, B. Thorand, J. Tichet, T. Tuomi, R. M. van Dam, T. W. van Haeften, T. van Herpt, J. V. van Vliet-Ostaptchouk, G. B. Walters, M. N. Weedon, C. Wijmenga, J. Witteman, R. N. Bergman, S. Cauchi, F. S. Collins, A. L. Gloyn, U. Gyllensten, T. Hansen, W. A. Hide, G. A. Hitman, A. Hofman, D. J. Hunter, K. Hveem, M. Laakso, K. L. Mohlke, A. D. Morris, C. N. Palmer, P. P. Pramstaller, I. Rudan, E. Sijbrands, L. D. Stein, J. Tuomilehto, A. Uitterlinden, M. Walker, N. J. Wareham, R. M. Watanabe, G. R. Abecasis, B. O. Boehm, H. Campbell, M. J. Daly, A. T. Hattersley, F. B. Hu, J. B. Meigs, J. S. Pankow, O. Pedersen, H. E. Wichmann, I. Barroso, J. C. Florez, T. M. Frayling, L. Groop, R. Sladek, U. Thorsteinsdottir, J. F. Wilson, T. Illig, P. Froguel, C. M. van Duijn, K. Stefansson, D. Altshuler, M. Boehnke, M. I. McCarthy, K. Ahmadi, C. Ainali, A. Barrett, V. Bataille, J. T. Bell, A. Buil, P. Deloukas, E. T. Dermitzakis, A. Dimas, R. Durbin, D. Glass, E. Grundberg, C. Hammond, N. Hassanali, Å . K. Hedman, C. Ingle, D. Knowles, M. Krestyaninova, C. Lindgren, C. Lowe, M. I. McCarthy, P. di Meglio, S. Montgomery, F. Nestle, A. C. Nica, J. Nisbet, S. O’Rahilly, L. Parts, S. Potter, M. Sekowska, S. Y. Shin, K. S. Small, N. Soranzo, T. D. Spector, G. Surdulescu, M. Travers, S. Tsoka, L. Tsaprouni, A. Wilk, T. P. Yang, and K. T. Zondervan. Meta-analysis of genome-wide association studies identifies eight new loci for type 2 diabetes in east Asians. Nat Genet, 44(1):67–72, Dec 2011.

36. T. F. Mackay, E. A. Stone, and J. F. Ayroles. The genetics of quantitative traits: challenges and prospects. Nat Rev Genet, 10(8):565–577, Aug 2009.

37. I. M. Ehrenreich, J. P. Gerke, and L. Kruglyak. Genetic dissection of complex traits in yeast: insights from studies of gene expression and other phenotypes in the BYxRM cross. Cold Spring Harb Symp Quant Biol, 74:145–153, 2009.

38. R. Christiano, H. Arlt, S. Kabatnik, N. Mejhert, Z. W. Lai, R. V. Farese, and T. C. Walther. A Systematic Protein Turnover Map for Decoding Protein Degradation. Cell Rep, 33(6):108378, 11 2020.

39. K. E. Kong, B. Fischer, M. Meurer, I. Kats, Z. Li, F. Rühle, J. D. Barry, D. Kirrmaier, V. Chevyreva, B. J. San Luis, M. Costanzo, W. Huber, B. J. Andrews, C. Boone, M. Knop, and A. Khmelinskii. Timer-based proteomic profiling of the ubiquitin-proteasome system reveals a substrate receptor of the GID ubiquitin ligase. Mol Cell, 81(11):2460–2476, 06 2021.

40. A. Battle, Z. Khan, S. H. Wang, A. Mitrano, M. J. Ford, J. K. Pritchard, and Y. Gilad. Genomic variation. Impact of regulatory variation from RNA to protein. Science, 347(6222):664–667, Feb 2015.

41. A. Ghazalpour, B. Bennett, V. A. Petyuk, L. Orozco, R. Hagopian, I. N. Mungrue, C. R. Farber, J. Sinsheimer, H. M. Kang, N. Furlotte, C. C. Park, P. Z. Wen, H. Brewer, K. Weitz, D. G. Camp, C. Pan, R. Yordanova, I. Neuhaus, C. Tilford, N. Siemers, P. Gargalovic, E. Eskin, T. Kirchgessner, D. J. Smith, R. D. Smith, and A. J. Lusis. Comparative analysis of proteome and transcriptome variation in mouse. PLoS Genet, 7(6):e1001393, Jun 2011.

42. B. A. Mirauta, D. D. Seaton, D. Bensaddek, A. Brenes, M. J. Bonder, H. Kilpinen, O. Stegle, A. I. Lamond, C. A. Agu, A. Alderton, P. Danecek, R. Denton, R. Durbin, D. J. Gaffney, A. Goncalves, R. Halai, S. Harper, C. M. Kirton, A. Kolb-Kokocinski, A. Leha, S. A. McCarthy, Y. Memari, M. Patel, E. Birney, F. P. Casale, L. Clarke, P. W. Harrison, H. Kilpinen, I. Streeter, D. Denovi, O. Stegle, A. I. Lamond, R. Meleckyte, N. Moens, F. M. Watt, W. H. Ouwehand, and P. Beales. Population-scale proteome variation in human induced pluripotent stem cells. Elife, 9, 08 2020.

43. F. W. Albert, S. Treusch, A. H. Shockley, J. S. Bloom, and L. Kruglyak. Genetics of single-cell protein abundance variation in large yeast populations. Nature, 506(7489):494–497, Feb 2014.

44. C. Brion, S. M. Lutz, and F. W. Albert. Simultaneous quantification of mRNA and protein in single cells reveals post-transcriptional effects of genetic variation. Elife, 9, Nov 2020.

45. J. M. Chick, S. C. Munger, P. Simecek, E. L. Huttlin, K. Choi, D. M. Gatti, N. Raghupathy, K. L. Svenson, G. A. Churchill, and S. P. Gygi. Defining the consequences of genetic variation on a proteome-wide scale. Nature, 534(7608):500–505, 06 2016.

46. C. Cenik, E. S. Cenik, G. W. Byeon, F. Grubert, S. I. Candille, D. Spacek, B. Alsallakh, H. Tilgner, C. L. Araya, H. Tang, E. Ricci, and M. P. Snyder. Integrative analysis of RNA, translation, and protein levels reveals distinct regulatory variation across humans. Genome Res, 25(11):1610–1621, Nov 2015.

47. N. S. Abell, M. K. DeGorter, M. J. Gloudemans, E. Greenwald, K. S. Smith, Z. He, and S. B. Montgomery. Multiple causal variants underlie genetic associations in humans. Science, 375(6586):1247– 1254, 03 2022.

48. E. J. Foss, D. Radulovic, S. A. Shaffer, D. R. Goodlett, L. Kruglyak, and A. Bedalov. Genetic variation shapes protein networks mainly through non-transcriptional mechanisms. PLoS Biol, 9(9):e1001144, Sep 2011.

49. J. S. Bloom, I. M. Ehrenreich, W. T. Loo, T. L. Lite, and L. Kruglyak. Finding the sources of missing heritability in a yeast cross. Nature, 494(7436):234–237, Feb 2013.

50. Y. Geffen, A. Appleboim, R. G. Gardner, N. Friedman, R. Sadeh, and T. Ravid. Mapping the Landscape of a Eukaryotic Degronome. Mol Cell, 63(6):1055–1065, 09 2016.

51. H. Yu, A. K. Singh Gautam, S. R. Wilmington, D. Wylie, K. Martinez-Fonts, G. Kago, M. Warburton, S. Chavali, T. Inobe, I. J. Finkelstein, M. M. Babu, and A. Matouschek. Conserved Sequence Preferences Contribute to Substrate Recognition by the Proteasome. J Biol Chem, 291(28):14526–14539, Jul 2016.

52. H. C. Yen, Q. Xu, D. M. Chou, Z. Zhao, and S. J. Elledge. Global protein stability profiling in mammalian cells. Science, 322(5903):918–923, Nov 2008.

53. A. Varshavsky. N-degron and C-degron pathways of protein degradation. Proc Natl Acad Sci U S A, 116(2):358–366, 01 2019.

54. V. Solomon, S. H. Lecker, A. L. Goldberg, and A. L. Goldberg. The N-end rule pathway catalyzes a major fraction of the protein degradation in skeletal muscle. J Biol Chem, 273(39):25216–25222, Sep 1998.

55. I. Kats, A. Khmelinskii, M. Kschonsak, F. Huber, R. A. Knieß, A. Bartosik, and M. Knop. Mapping Degradation Signals and Pathways in a Eukaryotic N-terminome. Mol Cell, 70(3):488–501, 05 2018.

56. A. Varshavsky. The N-end rule: functions, mysteries, uses. Proc Natl Acad Sci U S A, 93(22):12142– 12149, Oct 1996.

57. A. Varshavsky. Ubiquitin fusion technique and related methods. Meth. Enzymol., 399:777–799, 2005.

58. A. Khmelinskii, P. J. Keller, A. Bartosik, M. Meurer, J. D. Barry, B. R. Mardin, A. Kaufmann, S. Trautmann, M. Wachsmuth, G. Pereira, W. Huber, E. Schiebel, and M. Knop. Tandem fluorescent protein timers for in vivo analysis of protein dynamics. Nat. Biotechnol., 30(7):708–714, Jun 2012.

59. A. Khmelinskii and M. Knop. Analysis of protein dynamics with tandem fluorescent protein timers. Methods Mol. Biol., 1174:195–210, 2014.

60. A. Khmelinskii, E. Blaszczak, M. Pantazopoulou, B. Fischer, D. J. Omnus, G. Le Dez, A. Brossard, A. Gunnarsson, J. D. Barry, M. Meurer, D. Kirrmaier, C. Boone, W. Huber, G. Rabut, P. O. Ljungdahl, and M. Knop. Protein quality control at the inner nuclear membrane. Nature, 516(7531):410–413, Dec 2014.

61. Y. Xie and A. Varshavsky. RPN4 is a ligand, substrate, and transcriptional regulator of the 26S proteasome: a negative feedback circuit. Proc. Natl. Acad. Sci. U.S.A., 98(6):3056–3061, Mar 2001.

62. R. W. Michelmore, I. Paran, and R. V. Kesseli. Identification of markers linked to disease-resistance genes by bulked segregant analysis: a rapid method to detect markers in specific genomic regions by using segregating populations. Proc Natl Acad Sci U S A, 88(21):9828–9832, Nov 1991.

63. I. M. Ehrenreich, N. Torabi, Y. Jia, J. Kent, S. Martis, J. A. Shapiro, D. Gresham, A. A. Caudy, and L. Kruglyak. Dissection of genetically complex traits with extremely large pools of yeast segregants. Nature, 464(7291):1039–1042, Apr 2010.

64. S. Lutz, C. Brion, M. Kliebhan, and F. W. Albert. DNA variants affecting the expression of numerous genes in trans have diverse mechanisms of action and evolutionary histories. PLoS Genet, 15(11):e1008375, 11 2019.

65. B. Bartel, I. Winning, and A. Varshavsky. The recognition component of the N-end rule pathway. EMBO J., 9(10):3179–3189, Oct 1990.

66. J. Peter, M. De Chiara, A. Friedrich, J. X. Yue, D. Pflieger, A. Bergström, A. Sigwalt, B. Barre, K. Freel, A. Llored, C. Cruaud, K. Labadie, J. M. Aury, B. Istace, K. Lebrigand, P. Barbry, S. Engelen, A. Lemainque, P. Wincker, G. Liti, and J. Schacherer. Genome evolution across 1,011 Saccharomyces cerevisiae isolates. Nature, 556(7701):339–344, 04 2018.

67. K. Renganaath, R. Cheung, L. Day, S. Kosuri, L. Kruglyak, and F. W. Albert. Systematic identification of cis-regulatory variants that cause gene expression differences in a yeast cross. Elife, 9, Nov 2020.

68. E. Chen, Y. T. Kwon, M. S. Lim, I. D. Dubé, and M. R. Hough. Loss of Ubr1 promotes aneuploidy and accelerates B-cell lymphomagenesis in TLX1/HOX11-transgenic mice. Oncogene, 25(42):5752–5763, Sep 2006.

69. R. T. Baker and A. Varshavsky. Yeast N-terminal amidase. A new enzyme and component of the N-end rule pathway. J Biol Chem, 270(20):12065–12074, May 1995.

70. R. Swanson, M. Locher, and M. Hochstrasser. A conserved ubiquitin ligase of the nuclear envelope/endoplasmic reticulum that functions in both ER-associated and Matalpha2 repressor degradation. Genes Dev, 15(20):2660–2674, Oct 2001.

71. C. S. Hwang, A. Shemorry, and A. Varshavsky. N-terminal acetylation of cellular proteins creates specific degradation signals. Science, 327(5968):973–977, Feb 2010.

72. J. H. Oh, J. Y. Hyun, S. J. Chen, and A. Varshavsky. Five enzymes of the Arg/N-degron pathway form a targeting complex: The concept of superchanneling. Proc Natl Acad Sci U S A, 117(20):10778–10788, 05 2020.

73. M. Gaisne, A. M. Bécam, J. Verdière, and C. J. Herbert. A ’natural’ mutation in Saccharomyces cerevisiae strains derived from S288c affects the complex regulatory gene HAP1 (CYP1). Curr Genet, 36(4):195–200, Oct 1999.

74. F. W. Albert, J. S. Bloom, J. Siegel, L. Day, and L. Kruglyak. Genetics of trans-regulatory variation in gene expression. Elife, 7, 07 2018.

75. S. Jain, J. R. Wheeler, R. W. Walters, A. Agrawal, A. Barsic, and R. Parker. ATPase-Modulated Stress Granules Contain a Diverse Proteome and Substructure. Cell, 164(3):487–498, Jan 2016.

76. R. B. Wickner. MKT1, a nonessential Saccharomyces cerevisiae gene with a temperature-dependent effect on replication of M2 double-stranded RNA. J Bacteriol, 169(11):4941–4945, Nov 1987.

77. T. Icho, H. S. Lee, S. S. Sommer, and R. B. Wickner. Molecular characterization of chromosomal genes affecting double-stranded RNA replication in Saccharomyces cerevisiae. Basic Life Sci, 40:165–171, 1986.

78. A. V. Gomes. Genetics of proteasome diseases. Scientifica (Cairo), 2013:637629, 2013.

79. A. K. Agarwal, C. Xing, G. N. DeMartino, D. Mizrachi, M. D. Hernandez, A. B. Sousa, L. Martínez de Villarreal, H. G. dos Santos, and A. Garg. PSMB8 encoding the B5i proteasome subunit is mutated in joint contractures, muscle atrophy, microcytic anemia, and panniculitis-induced lipodystrophy syndrome. Am J Hum Genet, 87(6):866–872, Dec 2010.

80. H. X. Deng, W. Chen, S. T. Hong, K. M. Boycott, G. H. Gorrie, N. Siddique, Y. Yang, F. Fecto, Y. Shi, H. Zhai, H. Jiang, M. Hirano, E. Rampersaud, G. H. Jansen, S. Donkervoort, E. H. Bigio, B. R. Brooks, K. Ajroud, R. L. Sufit, J. L. Haines, E. Mugnaini, M. A. Pericak-Vance, and T. Siddique. Mutations in UBQLN2 cause dominant X-linked juvenile and adult-onset ALS and ALS/dementia. Nature, 477(7363):211–215, Aug 2011.

81. A. Kröll-Hermi, F. Ebstein, C. Stoetzel, V. Geoffroy, E. Schaefer, S. Scheidecker, S. Bär, M. Takamiya, K. Kawakami, B. A. Zieba, F. Studer, V. Pelletier, C. Eyermann, C. Speeg-Schatz, V. Laugel, D. Lipsker, F. Sandron, S. McGinn, A. Boland, J. F. Deleuze, L. Kuhn, J. Chicher, P. Hammann, S. Friant, C. Etard, E. Krüger, J. Muller, U. Strähle, and H. Dollfus. Proteasome subunit PSMC3 variants cause neurosensory syndrome combining deafness and cataract due to proteotoxic stress. EMBO Mol Med, 12(7):e11861, 07 2020.

82. D. Komander and M. Rape. The ubiquitin code. Annu Rev Biochem, 81:203–229, 2012.

83. E. S. Johnson, B. Bartel, W. Seufert, and A. Varshavsky. Ubiquitin as a degradation signal. EMBO J, 11(2):497–505, Feb 1992.

84. T. Inobe and A. Matouschek. Paradigms of protein degradation by the proteasome. Curr Opin Struct Biol, 24:156–164, Feb 2014.

85. S. Lutz, K. Van Dyke, M. A. Feraru, and F. W. Albert. Multiple epistatic DNA variants in a single gene affect gene expression in trans. Genetics, 220(1), 01 2022.

86. W. Li, M. H. Bengtson, A. Ulbrich, A. Matsuda, V. A. Reddy, A. Orth, S. K. Chanda, S. Batalov, and C. A. Joazeiro. Genome-wide and functional annotation of human E3 ubiquitin ligases identifies MULAN, a mitochondrial E3 that regulates the organelle’s dynamics and signaling. PLoS One, 3(1):e1487, Jan 2008.

87. A. Khmelinskii, M. Meurer, C. T. Ho, B. Besenbeck, J. F?ller, M. K. Lemberg, B. Bukau, A. Mogk, and M. Knop. Incomplete proteasomal degradation of green fluorescent proteins in the context of tandem fluorescent protein timers. Mol. Biol. Cell, 27(2):360–370, Jan 2016.

88. J. D. Pedelacq, S. Cabantous, T. Tran, T. C. Terwilliger, and G. S. Waldo. Engineering and characterization of a superfolder green fluorescent protein. Nat. Biotechnol., 24(1):79–88, Jan 2006.

89. N. C. Shaner, R. E. Campbell, P. A. Steinbach, B. N. Giepmans, A. E. Palmer, and R. Y. Tsien. Improved monomeric red, orange and yellow fluorescent proteins derived from Discosoma sp. red fluorescent protein. Nat. Biotechnol., 22(12):1567–1572, Dec 2004.

90. S. Kredel, F. Oswald, K. Nienhaus, K. Deuschle, C. R?cker, M. Wolff, R. Heilker, G. U. Nienhaus, and J. Wiedenmann. mRuby, a bright monomeric red fluorescent protein for labeling of subcellular structures. PLoS ONE, 4(2):e4391, 2009.

91. A. Grote, K. Hiller, M. Scheer, R. M?nch, B. N?rtemann, D. C. Hempel, and D. Jahn. JCat: a novel tool to adapt codon usage of a target gene to its potential expression host. Nucleic Acids Res., 33(Web Server issue):W526–531, Jul 2005.

92. A. Wach, A. Brachat, R. P?hlmann, and P. Philippsen. New heterologous modules for classical or PCR-based gene disruptions in Saccharomyces cerevisiae. Yeast, 10(13):1793–1808, Dec 1994.

93. A. L. Goldstein and J. H. McCusker. Three new dominant drug resistance cassettes for gene disruption in Saccharomyces cerevisiae. Yeast, 15(14):1541–1553, Oct 1999.

94. J. H. Zwolshen and J. K. Bhattacharjee. Genetic and biochemical properties of thialysine-resistant mutants of Saccharomyces cerevisiae. J Gen Microbiol, 122(2):281–287, Feb 1981.

95. A. Baryshnikova, M. Costanzo, S. Dixon, F. J. Vizeacoumar, C. L. Myers, B. Andrews, and C. Boone. Synthetic genetic array (SGA) analysis in Saccharomyces cerevisiae and Schizosaccharomyces pombe. Methods Enzymol, 470:145–179, 2010.

96. E. Kuzmin, M. Costanzo, B. Andrews, and C. Boone. Synthetic Genetic Array Analysis. Cold Spring Harb Protoc, 2016(4):pdb.prot088807, Apr 2016.

97. R. D. Gietz and R. H. Schiestl. High-efficiency yeast transformation using the LiAc/SS carrier DNA/PEG method. Nat Protoc, 2(1):31–34, 2007.

98. A. C. Ward. Rapid analysis of yeast transformants using colony-PCR. Biotechniques, 13(3):350, Sep 1992.

99. F. Hahne, N. LeMeur, R. R. Brinkman, B. Ellis, P. Haaland, D. Sarkar, J. Spidlen, E. Strain, and R. Gentleman. flowCore: a Bioconductor package for high throughput flow cytometry. BMC Bioinformatics, 10:106, Apr 2009.

100. Yoav Benjamini and Yosef Hochberg. Controlling the false discovery rate: A practical and powerful approach to multiple testing. Journal of the Royal Statistical Society Series B (Methodological), 57(1):289–300, 1995.

101. H. Li and R. Durbin. Fast and accurate short read alignment with Burrows-Wheeler transform. Bioinformatics, 25(14):1754–1760, Jul 2009.

102. H. Li, B. Handsaker, A. Wysoker, T. Fennell, J. Ruan, N. Homer, G. Marth, G. Abecasis, and R. Durbin. The Sequence Alignment/Map format and SAMtools. Bioinformatics, 25(16):2078–2079, Aug 2009.

103. M. D. Edwards and D. K. Gifford. High-resolution genetic mapping with pooled sequencing. BMC Bioinformatics, 13 Suppl 6:S8, Apr 2012.

104. R. S. Sikorski and P. Hieter. A system of shuttle vectors and yeast host strains designed for efficient manipulation of DNA in Saccharomyces cerevisiae. Genetics, 122(1):19–27, May 1989.

105. C. S. Hoffman and F. Winston. A ten-minute DNA preparation from yeast efficiently releases autonomous plasmids for transformation of Escherichia coli. Gene, 57(2-3):267–272, 1987.

106. R. M. Horton, H. D. Hunt, S. N. Ho, J. K. Pullen, and L. R. Pease. Engineering hybrid genes without the use of restriction enzymes: gene splicing by overlap extension. Gene, 77(1):61–68, Apr 1989.

107. S. Chen, Y. Zhou, Y. Chen, and J. Gu. fastp: an ultra-fast all-in-one FASTQ preprocessor. Bioinformatics, 34(17):i884–i890, 09 2018.

108. N. L. Bray, H. Pimentel, P. Melsted, and L. Pachter. Near-optimal probabilistic RNA-seq quantification. Nat Biotechnol, 34(5):525–527, 05 2016.

109. K. L. Howe, P. Achuthan, J. Allen, J. Allen, J. Alvarez-Jarreta, M. R. Amode, I. M. Armean, A. G. Azov, R. Bennett, J. Bhai, K. Billis, S. Boddu, M. Charkhchi, C. Cummins, L. Da Rin Fioretto, C. Davidson, K. Dodiya, B. El Houdaigui, R. Fatima, A. Gall, C. Garcia Giron, T. Grego, C. Guijarro-Clarke, L. Haggerty, A. Hemrom, T. Hourlier, O. G. Izuogu, T. Juettemann, V. Kaikala, M. Kay, I. Lavidas, T. Le, D. Lemos, J. Gonzalez Martinez, J. C. Marugán, T. Maurel, A. C. McMahon, S. Mohanan, B. Moore, M. Muffato, D. N. Oheh, D. Paraschas, A. Parker, A. Parton, I. Prosovetskaia, M. P. Sakthivel, A. I. A. Salam, B. M. Schmitt, H. Schuilenburg, D. Sheppard, E. Steed, M. Szpak, M. Szuba, K. Taylor, A. Thormann, G. Threadgold, B. Walts, A. Winterbottom, M. Chakiachvili, A. Chaubal, N. De Silva, B. Flint, A. Frankish, S. E. Hunt, G. R. IIsley, N. Langridge, J. E. Loveland, F. J. Martin, J. M. Mudge, J. Morales, E. Perry, M. Ruffier, J. Tate, D. Thybert, S. J. Trevanion, F. Cunningham, A. D. Yates, D. R. Zerbino, and P. Flicek. Ensembl 2021. Nucleic Acids Res, 49(D1):D884–D891, 01 2021.

110. L. Wang, S. Wang, and W. Li. RSeQC: quality control of RNA-seq experiments. Bioinformatics, 28(16):2184–2185, Aug 2012.

111. M. I. Love, W. Huber, and S. Anders. Moderated estimation of fold change and dispersion for RNA-seq data with DESeq2. Genome Biol, 15(12):550, 2014.

112. J. T. Leek, W. E. Johnson, H. S. Parker, A. E. Jaffe, and J. D. Storey. The sva package for removing batch effects and other unwanted variation in high-throughput experiments. Bioinformatics, 28(6):882– 883, Mar 2012.

113. J. T. Leek and J. D. Storey. Capturing heterogeneity in gene expression studies by surrogate variable analysis. PLoS Genet, 3(9):1724–1735, Sep 2007.

114. H. Mi, D. Ebert, A. Muruganujan, C. Mills, L. P. Albou, T. Mushayamaha, and P. D. Thomas. PANTHER version 16: a revised family classification, tree-based classification tool, enhancer regions and extensive API. Nucleic Acids Res, 49(D1):D394–D403, 01 2021.

115. J. Jumper, R. Evans, A. Pritzel, T. Green, M. Figurnov, O. Ronneberger, K. Tunyasuvunakool, R. Bates, A. Žídek, A. Potapenko, A. Bridgland, C. Meyer, S. A. A. Kohl, A. J. Ballard, A. Cowie, B. Romera-Paredes, S. Nikolov, R. Jain, J. Adler, T. Back, S. Petersen, D. Reiman, E. Clancy, M. Zielinski, M. Steinegger, M. Pacholska, T. Berghammer, S. Bodenstein, D. Silver, O. Vinyals, A. W. Senior, K. Kavukcuoglu, P. Kohli, and D. Hassabis. Highly accurate protein structure prediction with AlphaFold. Nature, 596(7873):583–589, Aug 2021.

116. E. F. Pettersen, T. D. Goddard, C. C. Huang, E. C. Meng, G. S. Couch, T. I. Croll, J. H. Morris, and T. E. Ferrin. UCSF ChimeraX: Structure visualization for researchers, educators, and developers. Protein Sci, 30(1):70–82, 01 2021.

117. C. G. de Boer and T. R. Hughes. YeTFaSCo: a database of evaluated yeast transcription factor sequence specificities. Nucleic Acids Res, 40(Database issue):D169–179, Jan 2012.

118. C. A. Gilchrist, D. A. Gray, and R. T. Baker. A ubiquitin-specific protease that efficiently cleaves the ubiquitin-proline bond. J Biol Chem, 272(51):32280–32285, Dec 1997.

119. E. S. Johnson, P. C. Ma, I. M. Ota, and A. Varshavsky. A proteolytic pathway that recognizes ubiquitin as a degradation signal. J Biol Chem, 270(29):17442–17456, Jul 1995.

