## Supplementary material for "Variation in Ubiquitin System Genes Creates Substrate-Specific Effects on Proteasomal Protein Degradation": All N-degron QTL Mapping Traces

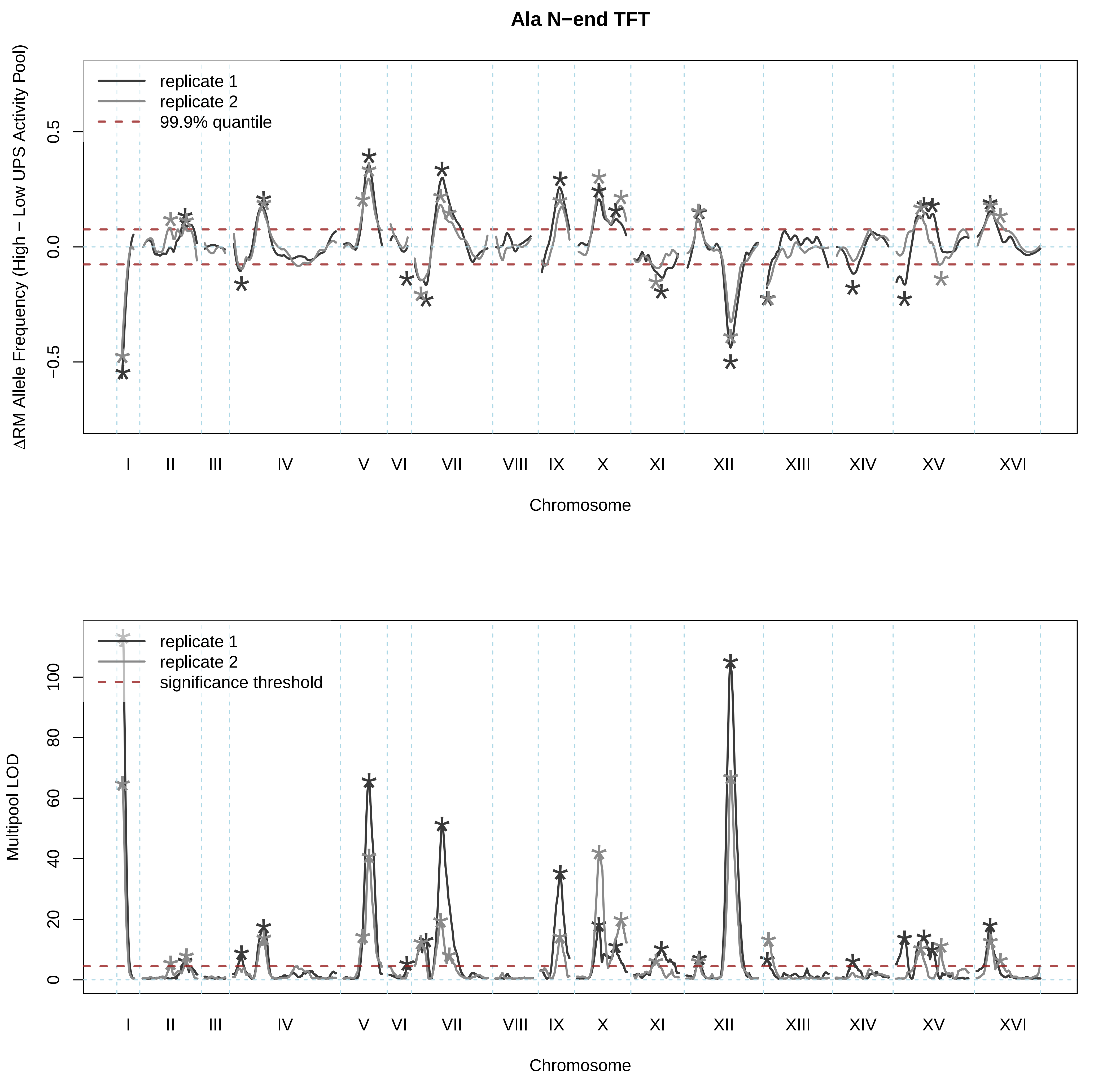

Ala N−end TFT
replicate 1
replicate 2
99.9% quantile
ΔRM Allele Frequency (High − Low UPS Activity Pool)
0.5
0.0
−0.5
I
II
III
IV
V
VI
VII
VIII
IX
X
XI
XII
XIII
XIV
XV
XVI
Chromosome
replicate 1
replicate 2
significance threshold
Multipool LOD
100
80
60
40
20
0
I
II
III
IV
V
VI
VII
VIII
IX
X
XI
XII
XIII
XIV
XV
XVI
Chromosome

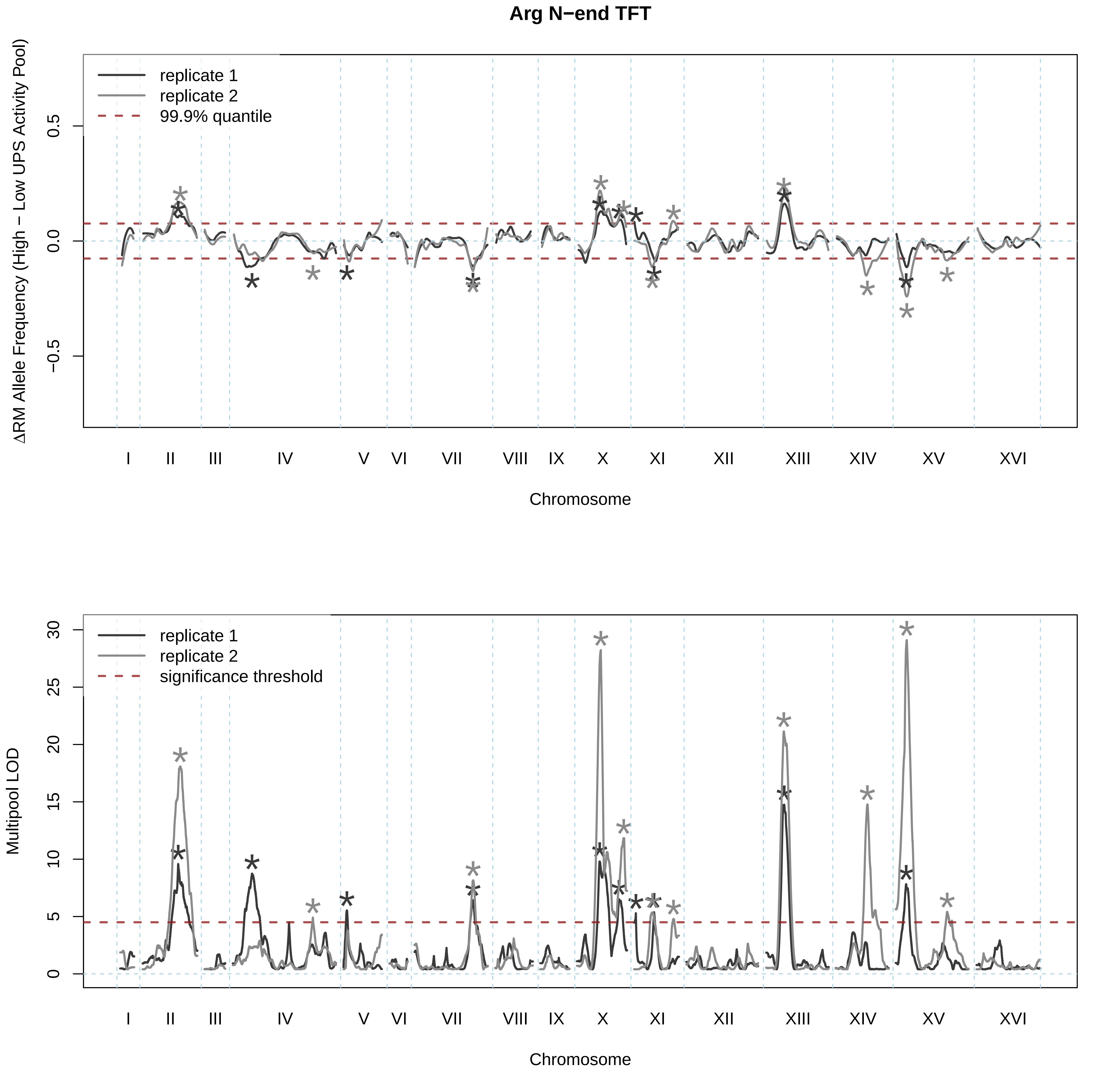
Arg N−end TFT
replicate 1
replicate 2
99.9% quantile
ΔRM Allele Frequency (High − Low UPS Activity Pool)
0.5
0.0
−0.5
I
II
III
IV
V
VI
VII
VIII
IX
X
XI
XII
XIII
XIV
XV
XVI
Chromosome
replicate 1
replicate 2
significance threshold
Multipool LOD
0
5
10
15
20
25
30
Chromosome

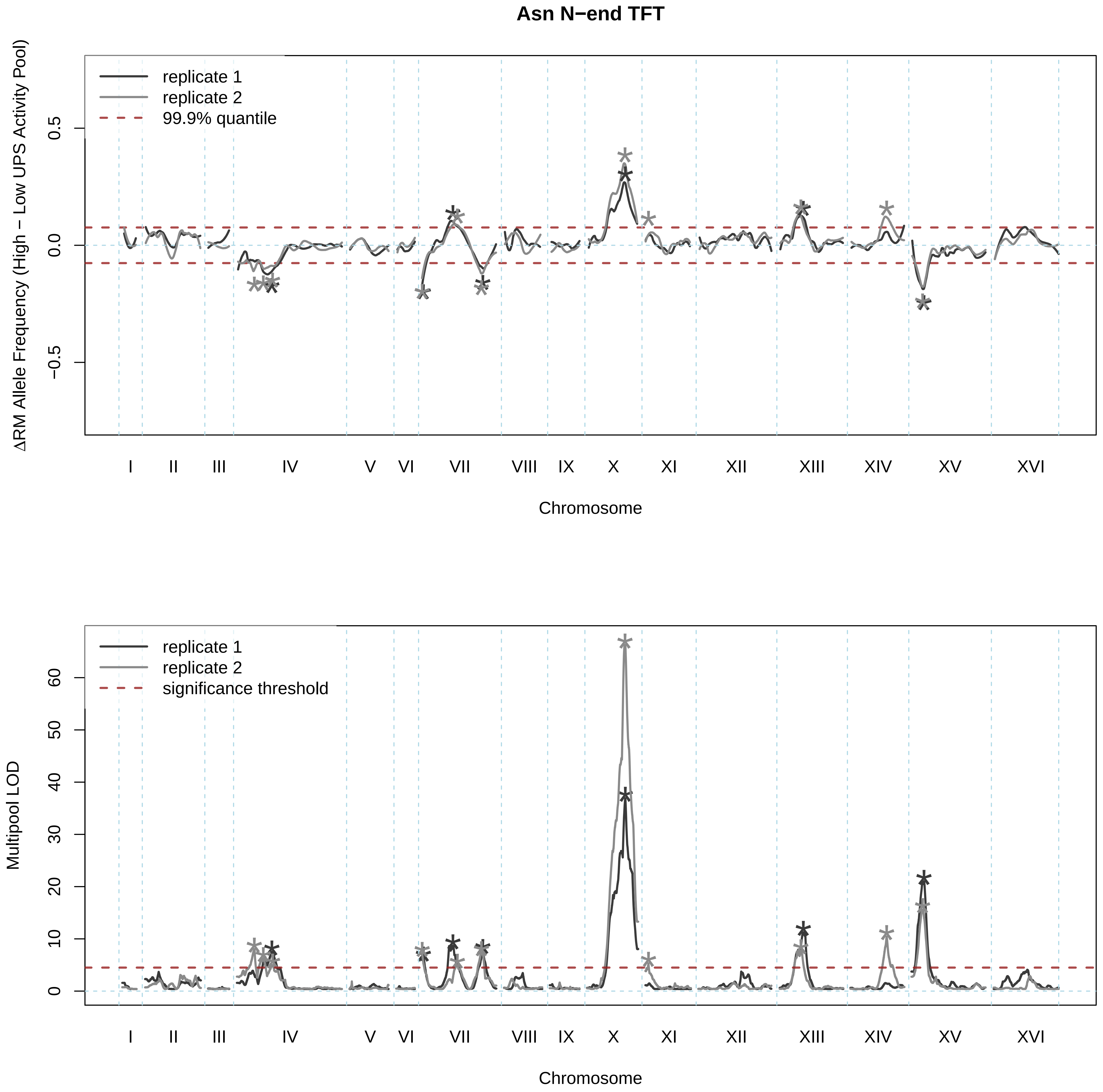
Asn N−end TFT
replicate 1
replicate 2
99.9% quantile
ΔRM Allele Frequency (High − Low UPS Activity Pool)
0.5
0.0
−0.5
I
II
III
IV
V
VI
VII
VIII
IX
X
XI
XII
XIII
XIV
XV
XVI
Chromosome
replicate 1
replicate 2
significance threshold
Multipool LOD
60
50
40
30
20
10
0
I
II
III
IV
V
VI
VII
VIII
IX
X
XI
XII
XIII
XIV
XV
XVI
Chromosome

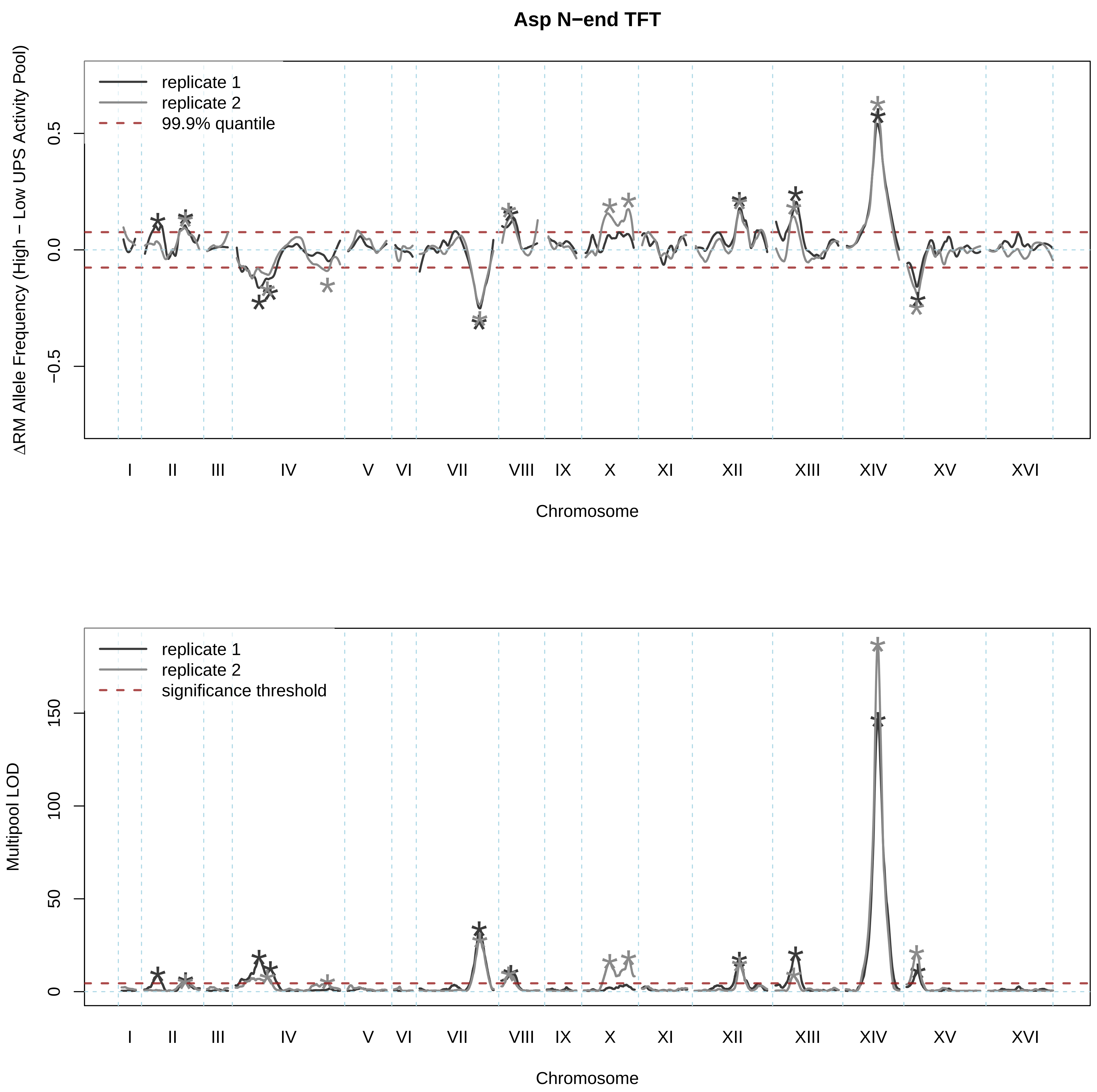

Asp N−end TFT
replicate 1
replicate 2
99.9% quantile
ΔRM Allele Frequency (High − Low UPS Activity Pool)
0.5
0.0
−0.5
I
II
III
IV
V
VI
VII
VIII
IX
X
XI
XII
XIII
XIV
XV
XVI
Chromosome
replicate 1
replicate 2
significance threshold
Multipool LOD
150
100
50
0
I
II
III
IV
V
VI
VII
VIII
IX
X
XI
XII
XIII
XIV
XV
XVI
Chromosome

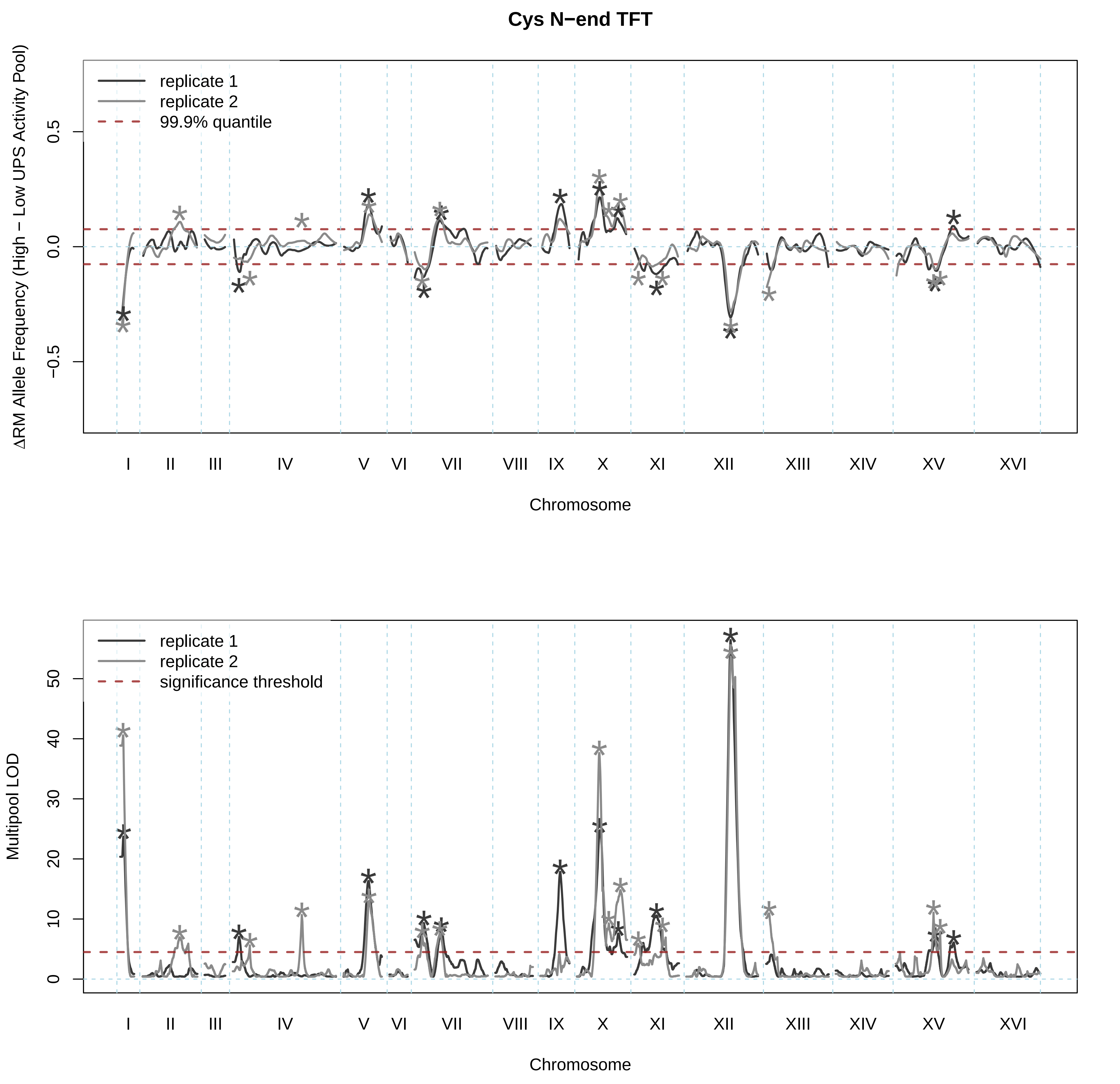

Cys N−end TFT
replicate 1
replicate 2
99.9% quantile
ΔRM Allele Frequency (High − Low UPS Activity Pool)
0.5
0.0
−0.5
I
II
III
IV
V
VI
VII
VIII
IX
X
XI
XII
XIII
XIV
XV
XVI
Chromosome
replicate 1
replicate 2
significance threshold
Multipool LOD
0
10
20
30
40
50
I
II
III
IV
V
VI
VII
VIII
IX
X
XI
XII
XIII
XIV
XV
XVI
Chromosome

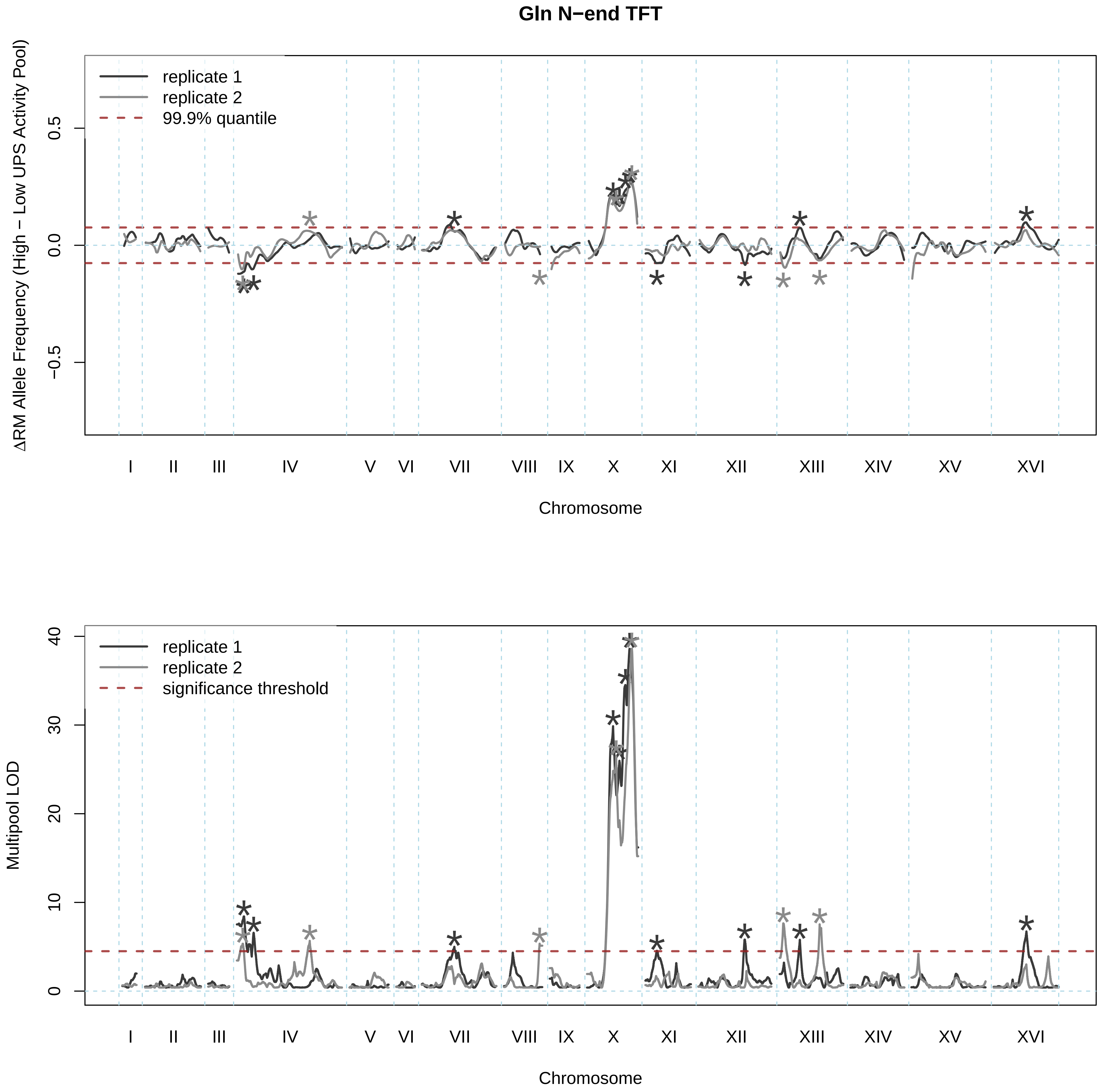

Gln N−end TFT
replicate 1
replicate 2
99.9% quantile
ΔRM Allele Frequency (High − Low UPS Activity Pool)
0.5
0.0
−0.5
I
II
III
IV
V
VI
VII
VIII
IX
X
XI
XII
XIII
XIV
XV
XVI
Chromosome
replicate 1
replicate 2
significance threshold
Multipool LOD
40
30
20
10
0
I
II
III
IV
V
VI
VII
VIII
IX
X
XI
XII
XIII
XIV
XV
XVI
Chromosome

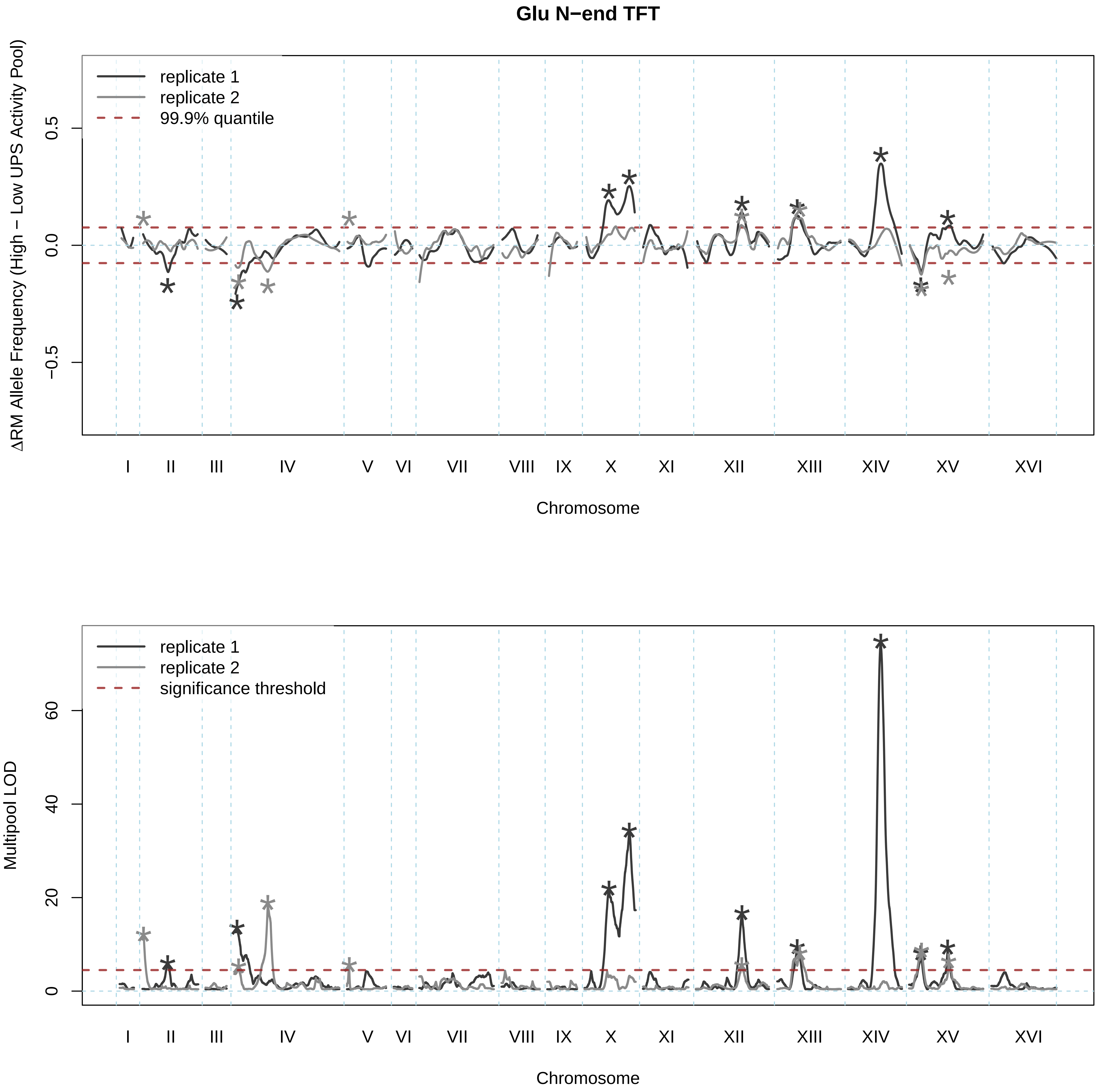
Glu N−end TFT
replicate 1
replicate 2
99.9% quantile
ΔRM Allele Frequency (High − Low UPS Activity Pool)
0.5
0.0
−0.5
I
II
III
IV
V
VI
VII
VIII
IX
X
XI
XII
XIII
XIV
XV
XVI
Chromosome
replicate 1
replicate 2
significance threshold
Multipool LOD
60
40
20
0
I
II
III
IV
V
VI
VII
VIII
IX
X
XI
XII
XIII
XIV
XV
XVI
Chromosome

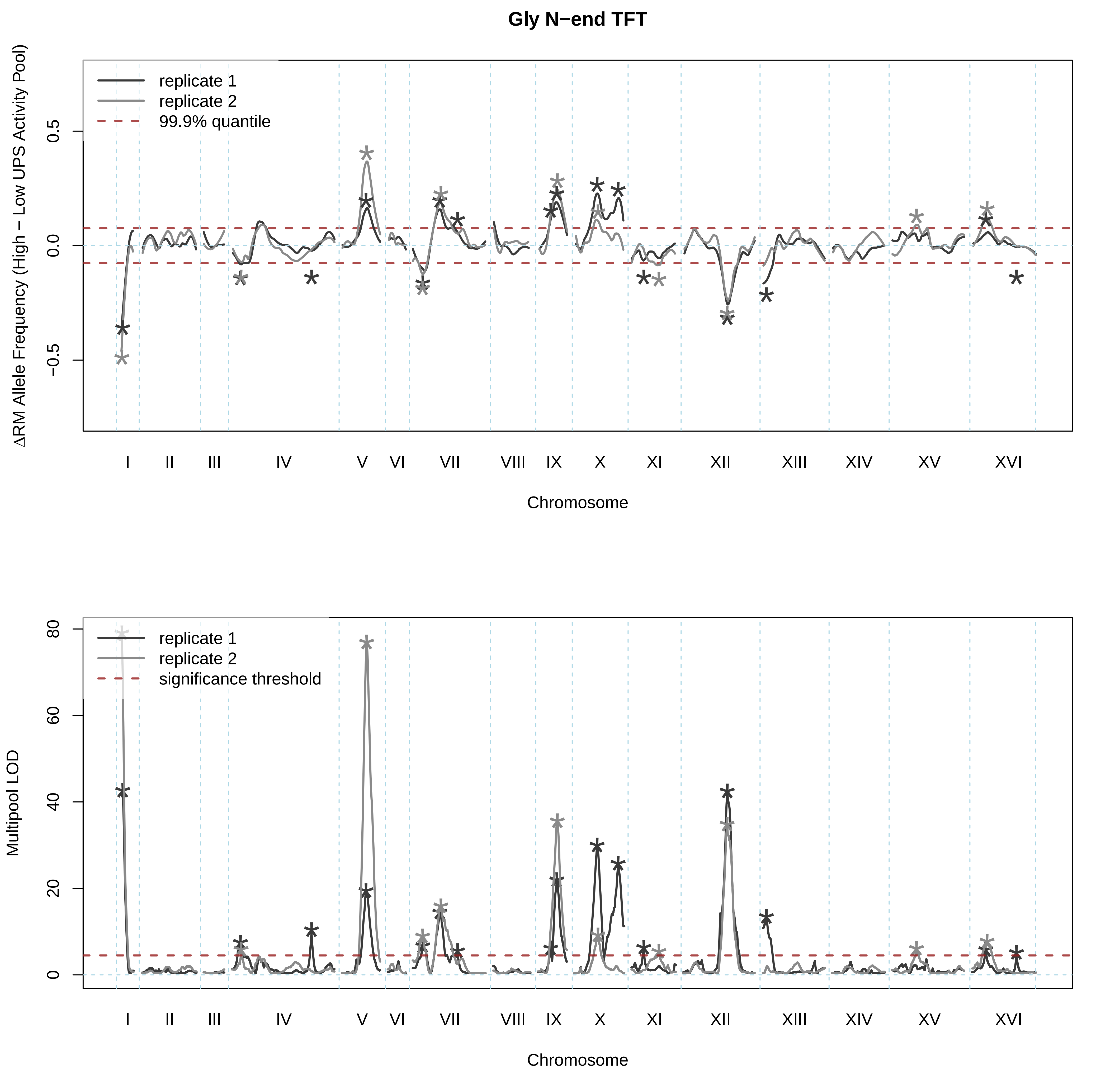
Gly N−end TFT
replicate 1
replicate 2
99.9% quantile
ΔRM Allele Frequency (High − Low UPS Activity Pool)
0.5
0.0
−0.5
I
II
III
IV
V
VI
VII
VIII
IX
X
XI
XII
XIII
XIV
XV
XVI
Chromosome
replicate 1
replicate 2
significance threshold
Multipool LOD
80
60
40
20
0
Chromosome

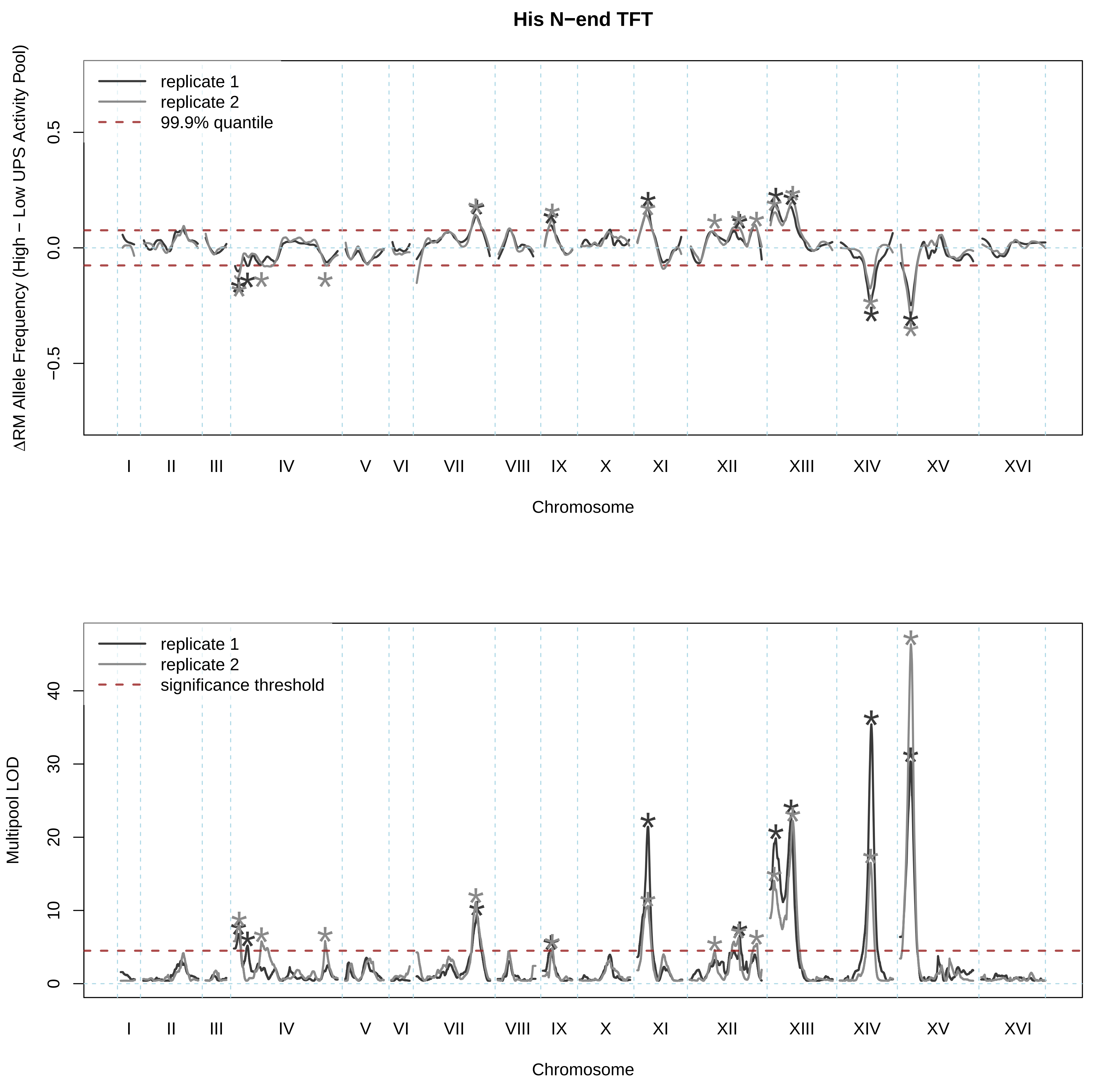
His N−end TFT
ΔRM Allele Frequency (High − Low UPS Activity Pool)
replicate 1
replicate 2
99.9% quantile
0.5
0.0
−0.5
I II III IV V VI VII VIII IX X XI XII XIII XIV XV XVI
Chromosome
Multipool LOD
replicate 1
replicate 2
significance threshold
0
10
20
30
40
I II III IV V VI VII VIII IX X XI XII XIII XIV XV XVI
Chromosome

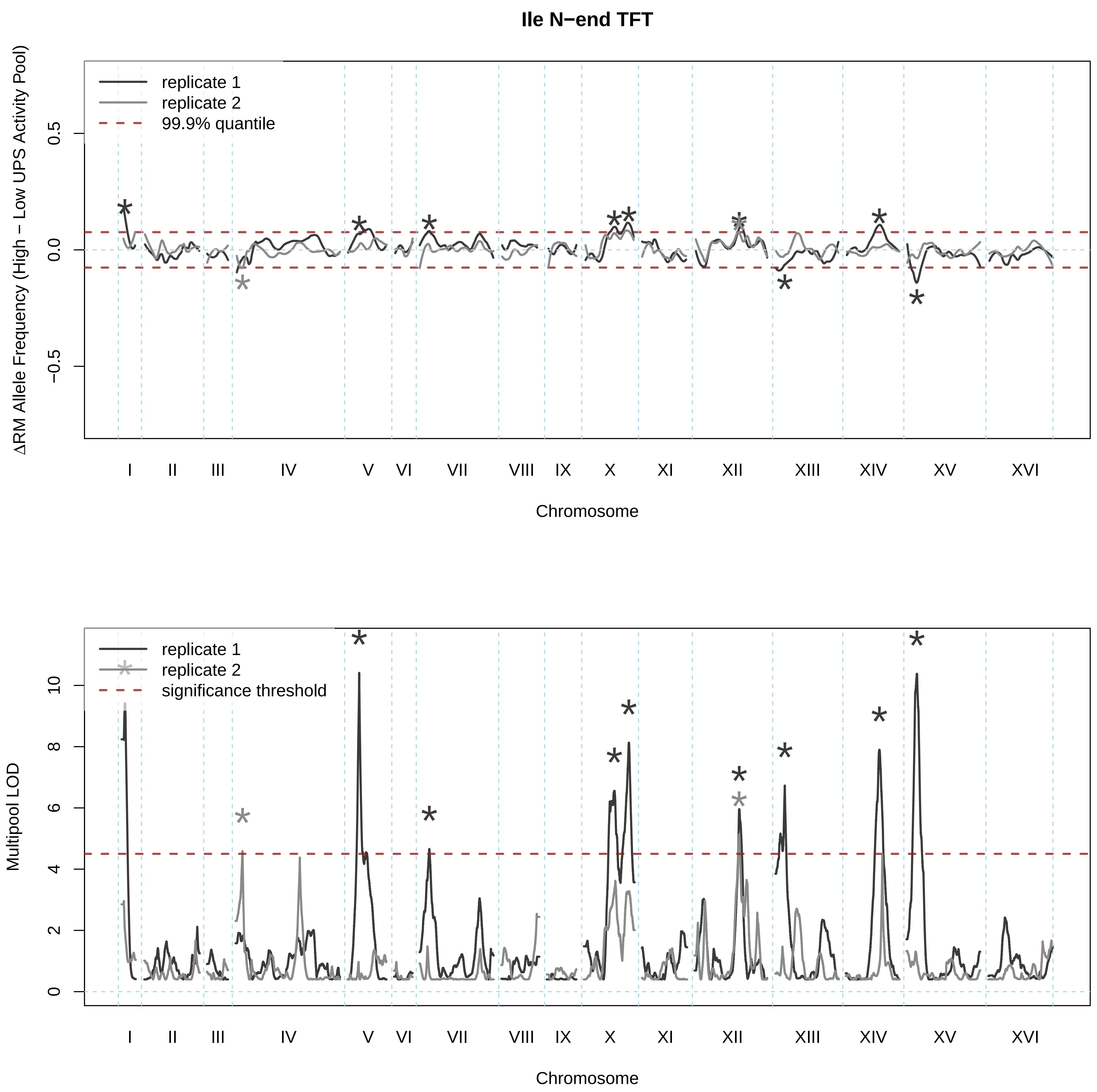

Ile N−end TFT
replicate 1
replicate 2
99.9% quantile
ΔRM Allele Frequency (High − Low UPS Activity Pool)
0.5
0.0
−0.5
I
II
III
IV
V
VI
VII
VIII
IX
X
XI
XII
XIII
XIV
XV
XVI
Chromosome
replicate 1
replicate 2
significance threshold
Multipool LOD
10
8
6
4
2
0
I
II
III
IV
V
VI
VII
VIII
IX
X
XI
XII
XIII
XIV
XV
XVI
Chromosome

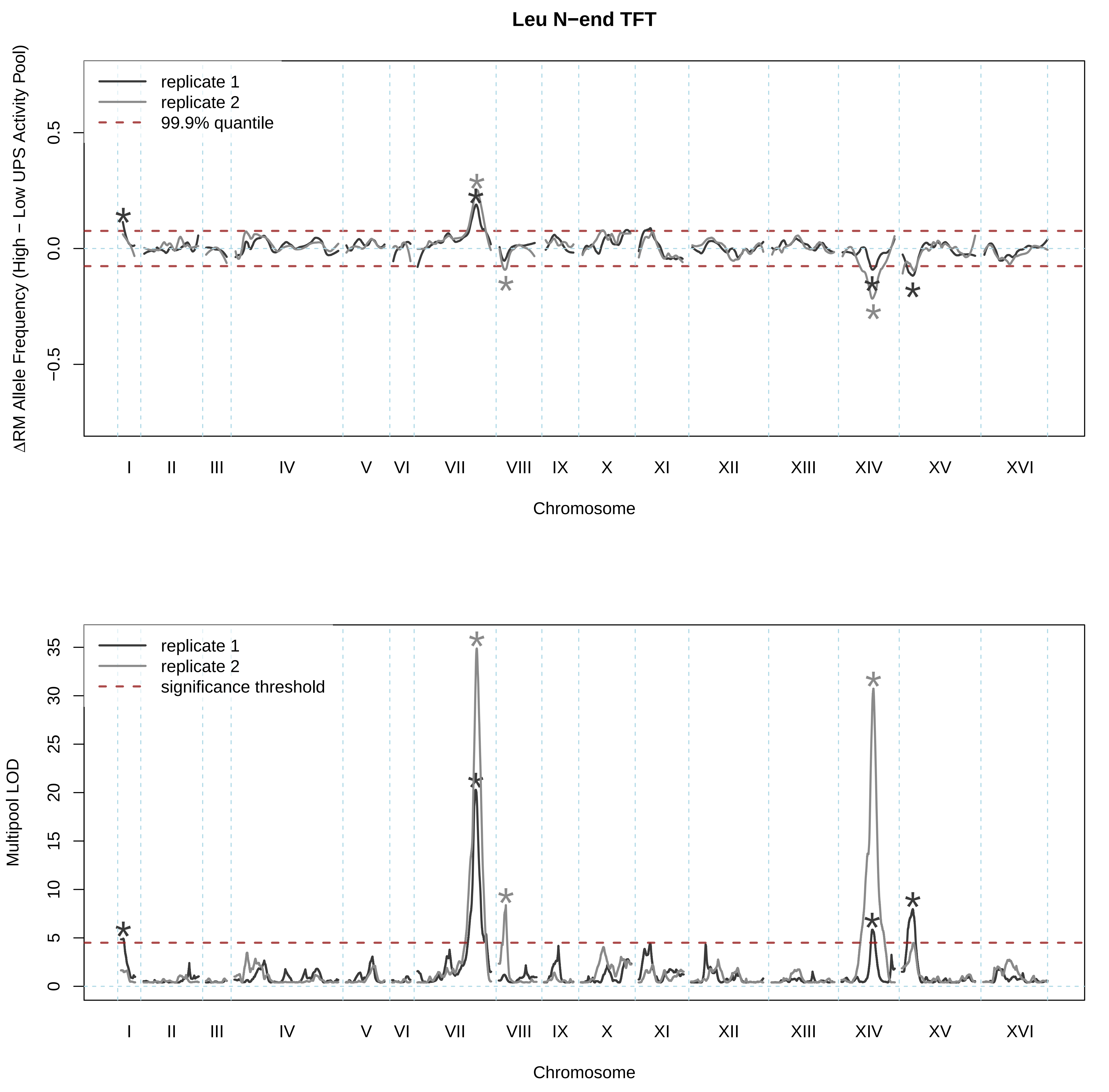
Leu N−end TFT
replicate 1
replicate 2
99.9% quantile
ΔRM Allele Frequency (High − Low UPS Activity Pool)
0.5
0.0
−0.5
I II III IV V VI VII VIII IX X XI XII XIII XIV XV XVI
Chromosome
replicate 1
replicate 2
significance threshold
Multipool LOD
35
30
25
20
15
10
5
0
I II III IV V VI VII VIII IX X XI XII XIII XIV XV XVI
Chromosome

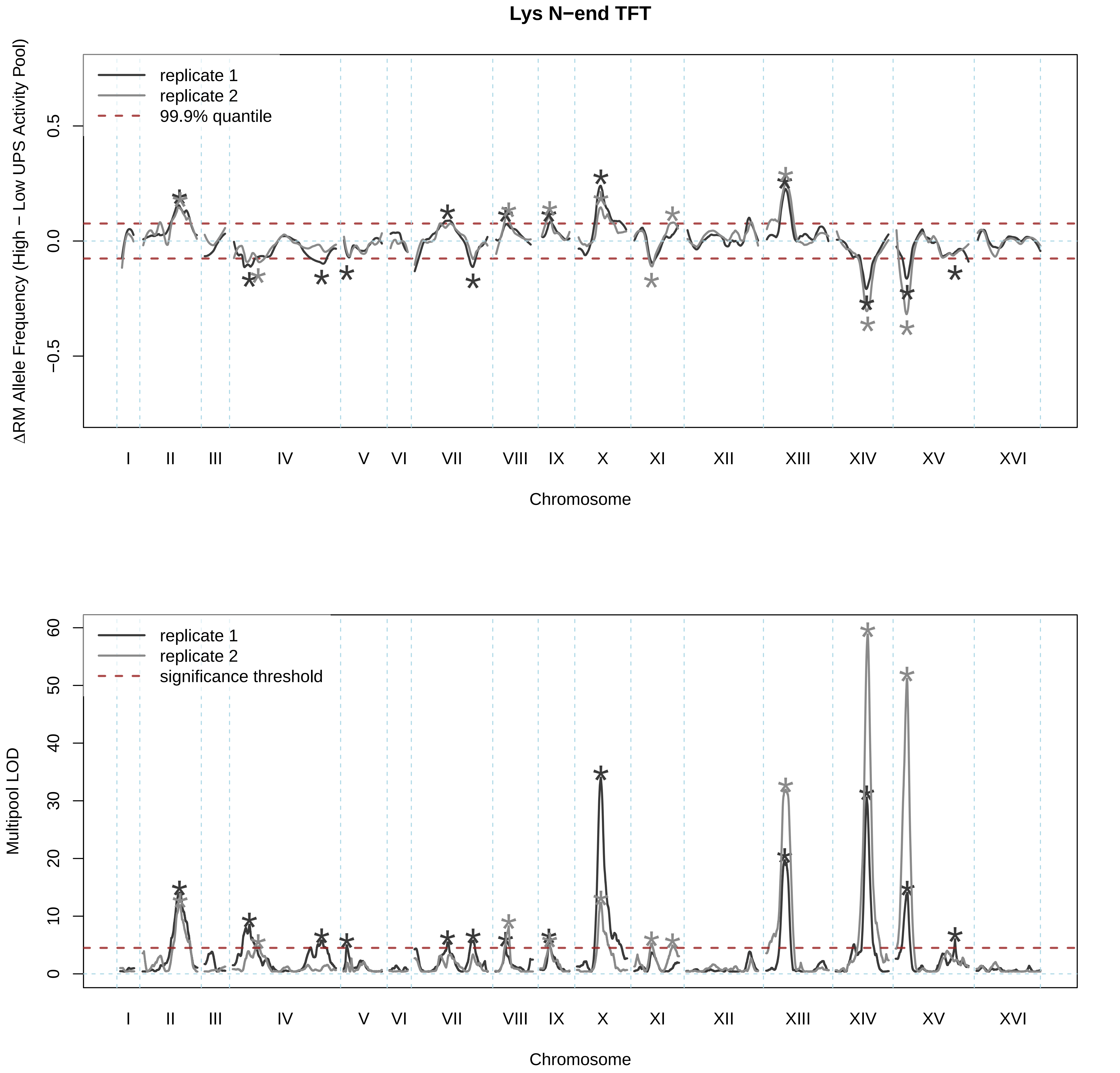

Lys N−end TFT
replicate 1
replicate 2
99.9% quantile
ΔRM Allele Frequency (High − Low UPS Activity Pool)
0.5
0.0
−0.5
I
II
III
IV
V
VI
VII
VIII
IX
X
XI
XII
XIII
XIV
XV
XVI
Chromosome
replicate 1
replicate 2
significance threshold
Multipool LOD
60
50
40
30
20
10
0
I
II
III
IV
V
VI
VII
VIII
IX
X
XI
XII
XIII
XIV
XV
XVI
Chromosome

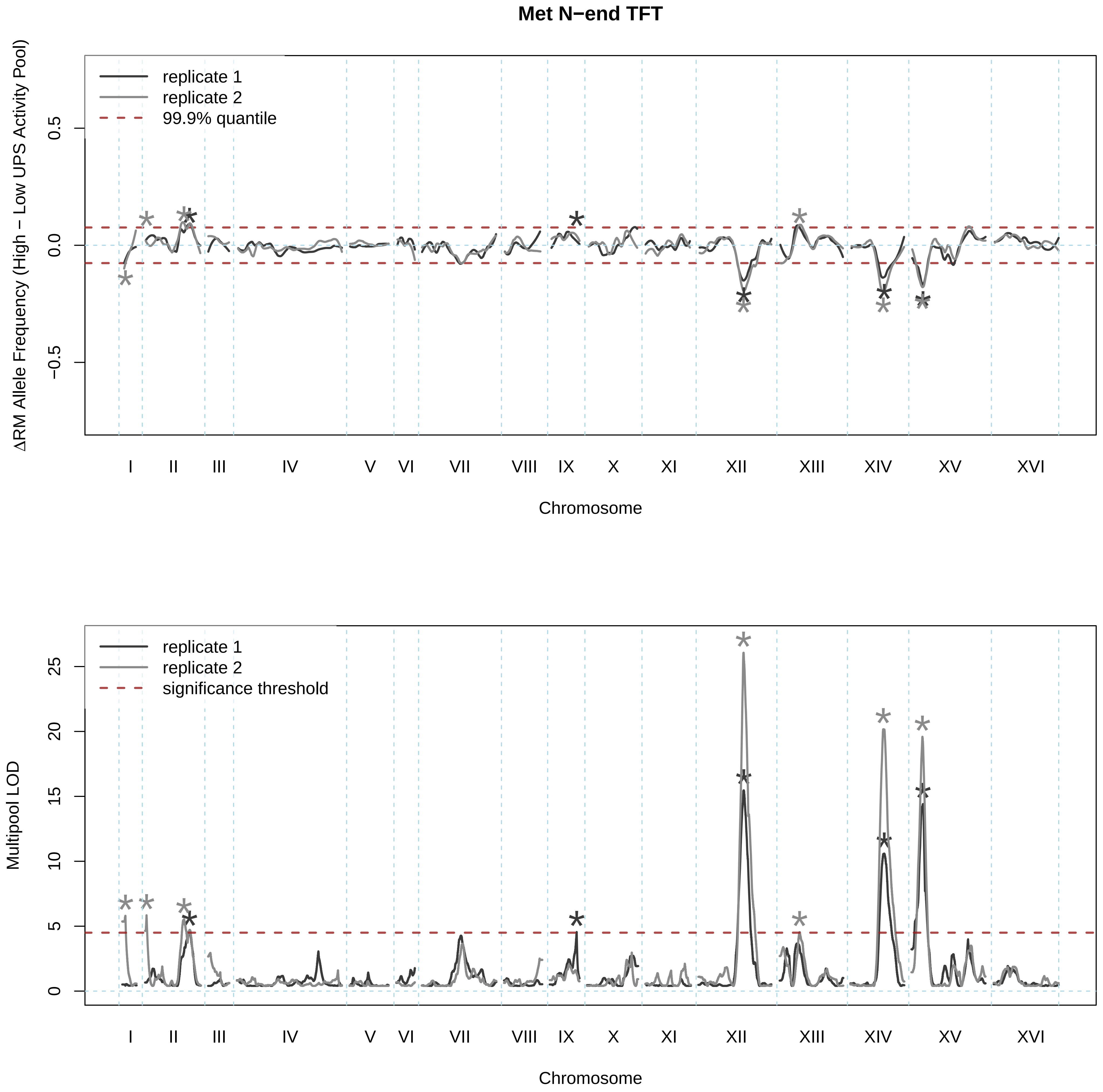
Met N−end TFT
ΔRM Allele Frequency (High − Low UPS Activity Pool)
replicate 1
replicate 2
99.9% quantile
0.5
0.0
−0.5
I
II
III
IV
V
VI
VII
VIII
IX
X
XI
XII
XIII
XIV
XV
XVI
Chromosome
Multipool LOD
replicate 1
replicate 2
significance threshold
25
20
15
10
5
0
I
II
III
IV
V
VI
VII
VIII
IX
X
XI
XII
XIII
XIV
XV
XVI
Chromosome

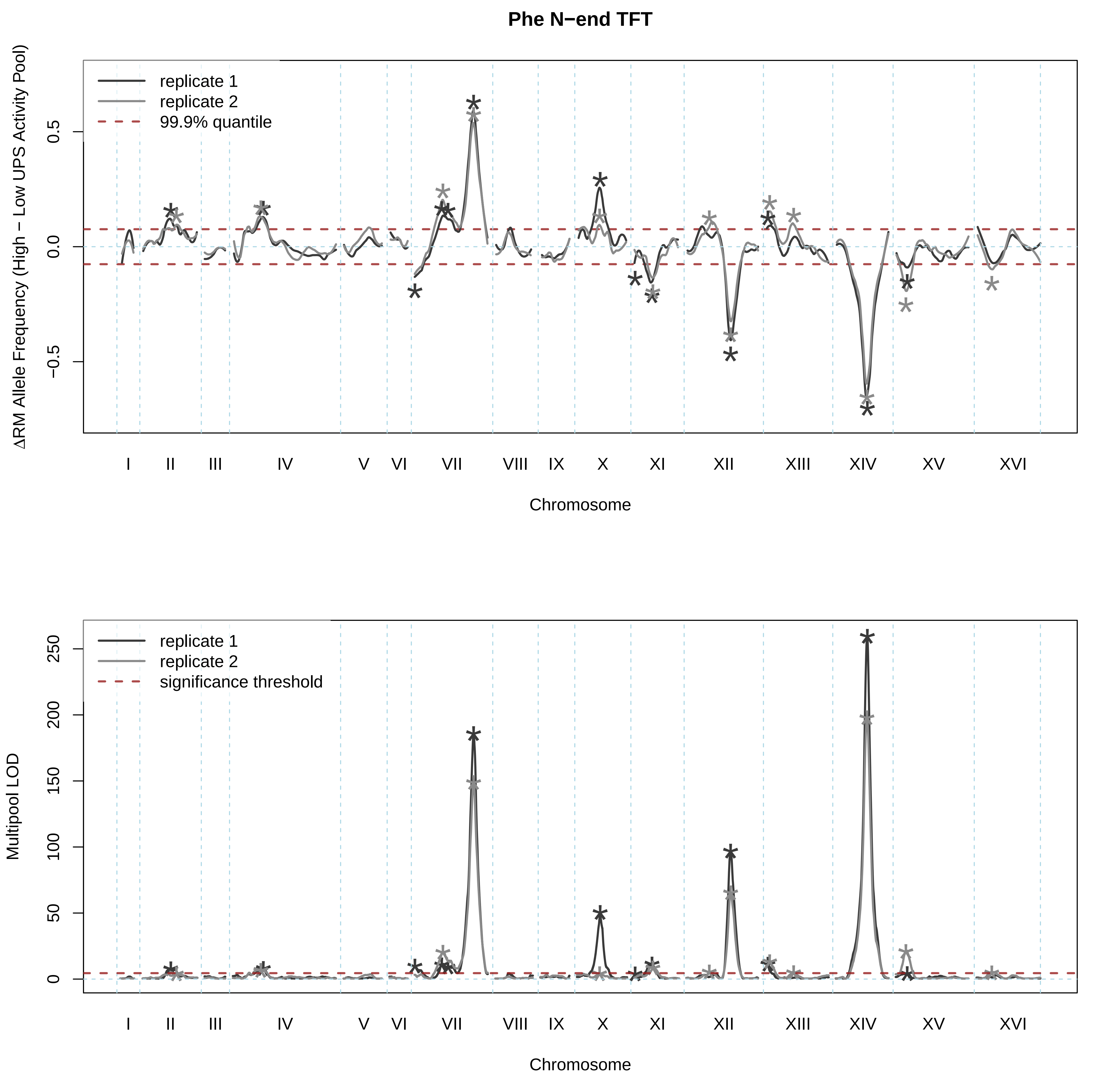

Phe N−end TFT
replicate 1
replicate 2
99.9% quantile
ΔRM Allele Frequency (High − Low UPS Activity Pool)
0.5
0.0
−0.5
I
II
III
IV
V
VI
VII
VIII
IX
X
XI
XII
XIII
XIV
XV
XVI
Chromosome
replicate 1
replicate 2
significance threshold
Multipool LOD
250
200
150
100
50
0
I
II
III
IV
V
VI
VII
VIII
IX
X
XI
XII
XIII
XIV
XV
XVI
Chromosome

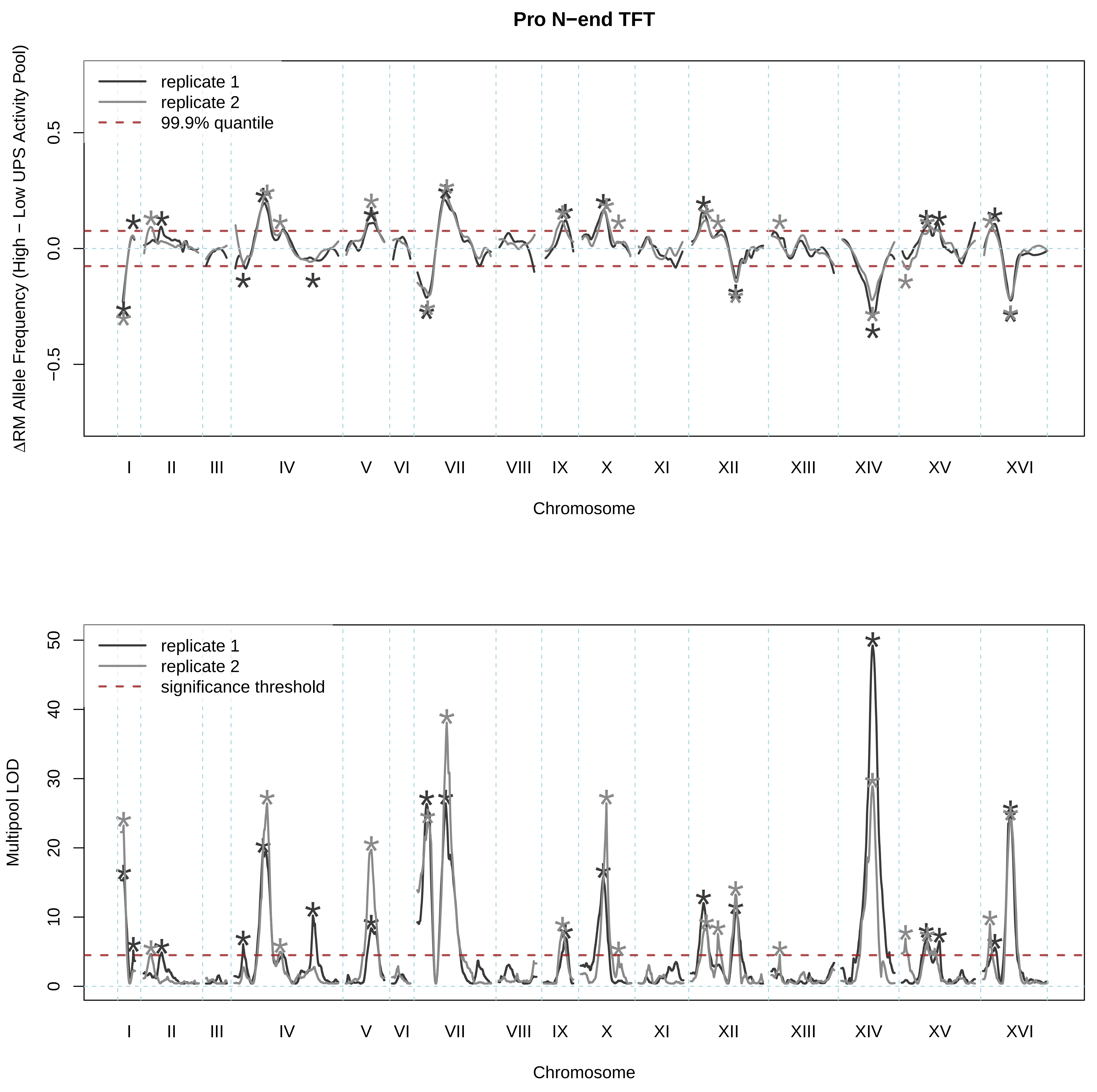

Pro N−end TFT
replicate 1
replicate 2
99.9% quantile
ΔRM Allele Frequency (High − Low UPS Activity Pool)
0.5
0.0
−0.5
I
II
III
IV
V
VI
VII
VIII
IX
X
XI
XII
XIII
XIV
XV
XVI
Chromosome
replicate 1
replicate 2
significance threshold
Multipool LOD
50
40
30
20
10
0
Chromosome

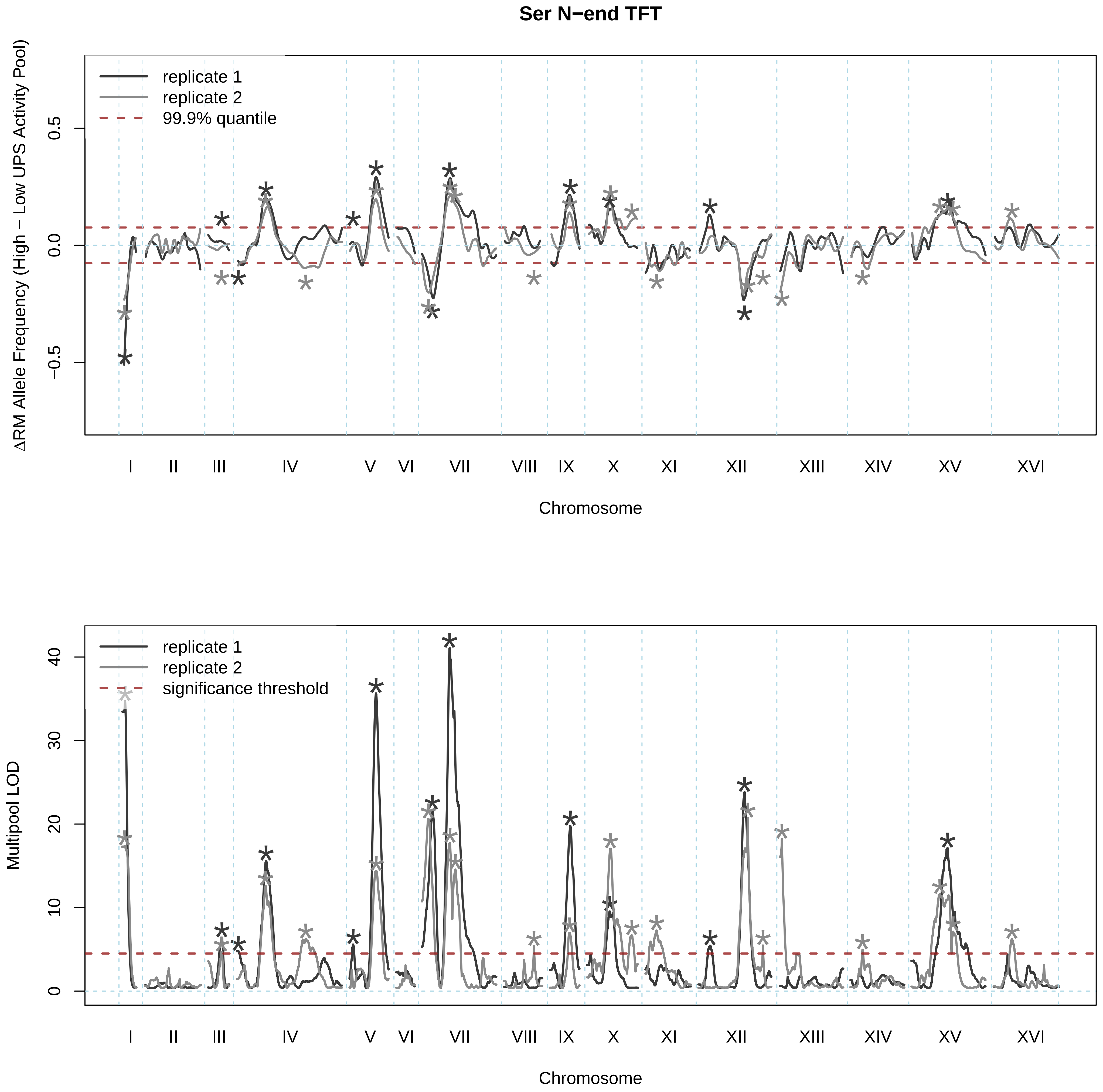
Ser N−end TFT
ΔRM Allele Frequency (High − Low UPS Activity Pool)
replicate 1
replicate 2
99.9% quantile
0.5
0.0
−0.5
I
II
III
IV
V
VI
VII
VIII
IX
X
XI
XII
XIII
XIV
XV
XVI
Chromosome
Multipool LOD
replicate 1
replicate 2
significance threshold
40
30
20
10
0
I
II
III
IV
V
VI
VII
VIII
IX
X
XI
XII
XIII
XIV
XV
XVI
Chromosome

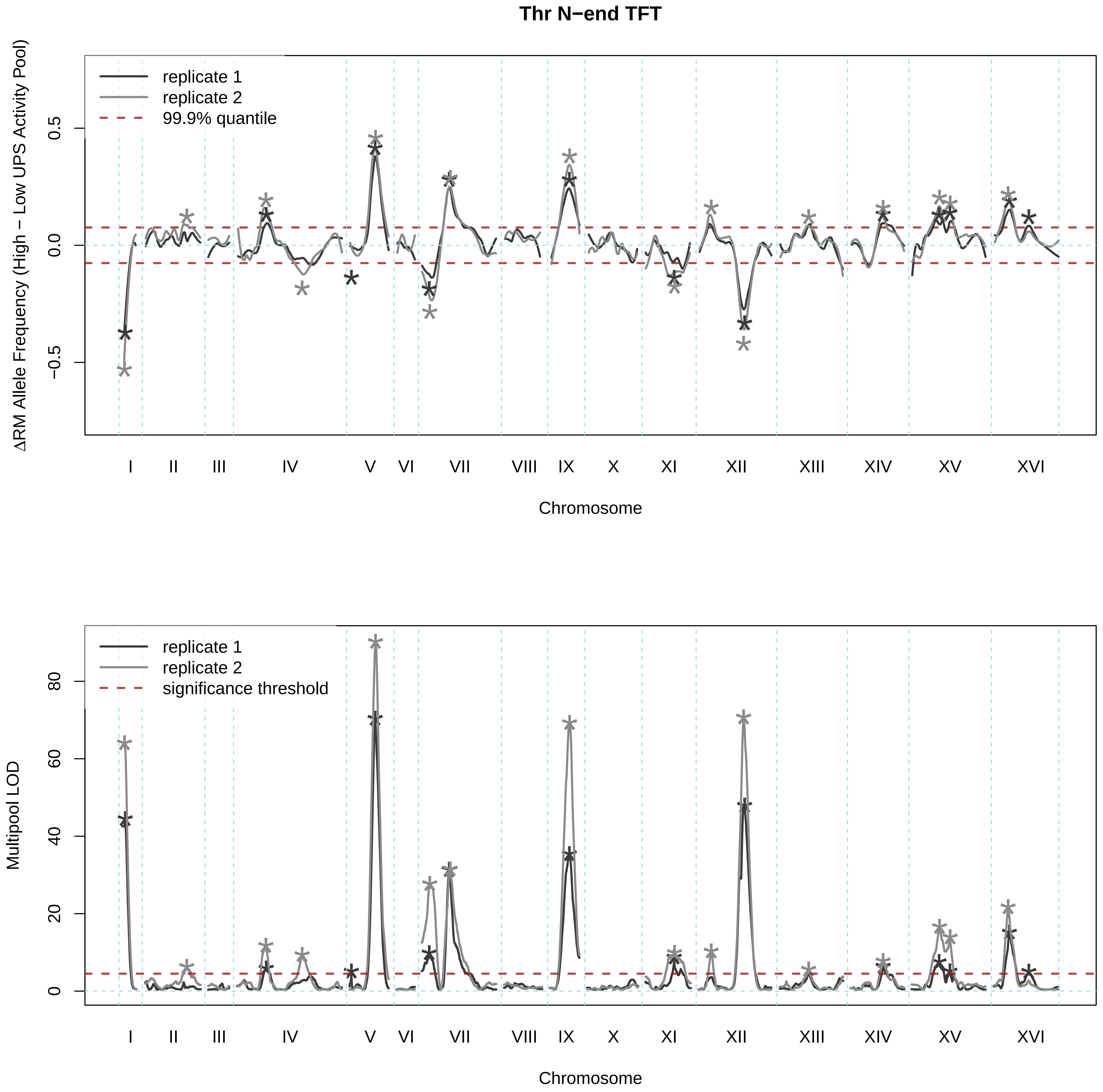
Thr N−end TFT
replicate 1
replicate 2
99.9% quantile
ΔRM Allele Frequency (High − Low UPS Activity Pool)
0.5
0.0
−0.5
I
II
III
IV
V
VI
VII
VIII
IX
X
XI
XII
XIII
XIV
XV
XVI
Chromosome
replicate 1
replicate 2
significance threshold
Multipool LOD
80
60
40
20
0
I
II
III
IV
V
VI
VII
VIII
IX
X
XI
XII
XIII
XIV
XV
XVI
Chromosome

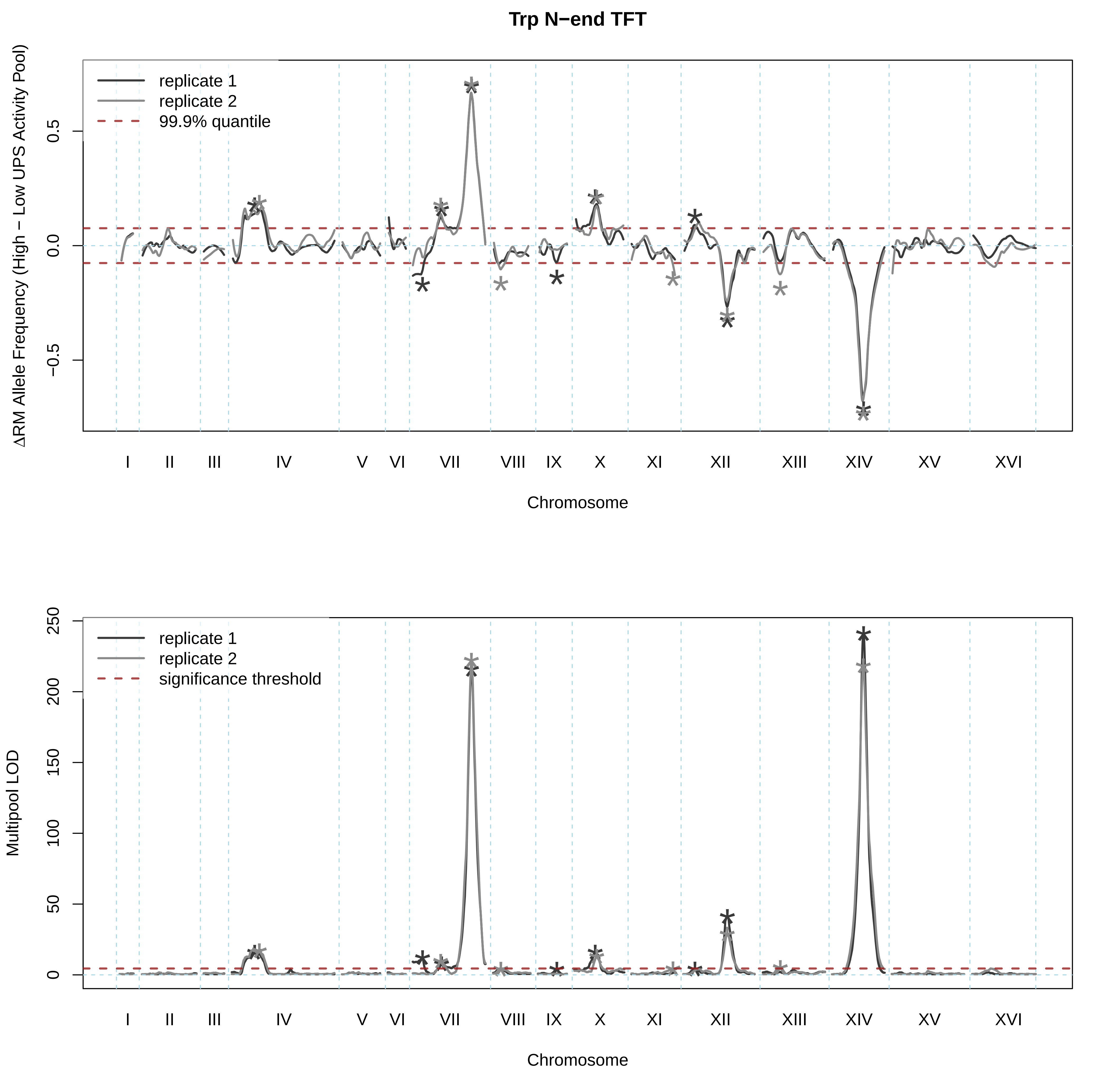

Trp N−end TFT
ΔRM Allele Frequency (High − Low UPS Activity Pool)
replicate 1
replicate 2
99.9% quantile
0.5
0.0
−0.5
I
II
III
IV
V
VI
VII
VIII
IX
X
XI
XII
XIII
XIV
XV
XVI
Chromosome
Multipool LOD
replicate 1
replicate 2
significance threshold
250
200
150
100
50
0
I
II
III
IV
V
VI
VII
VIII
IX
X
XI
XII
XIII
XIV
XV
XVI
Chromosome

### Tyr N−end TFT

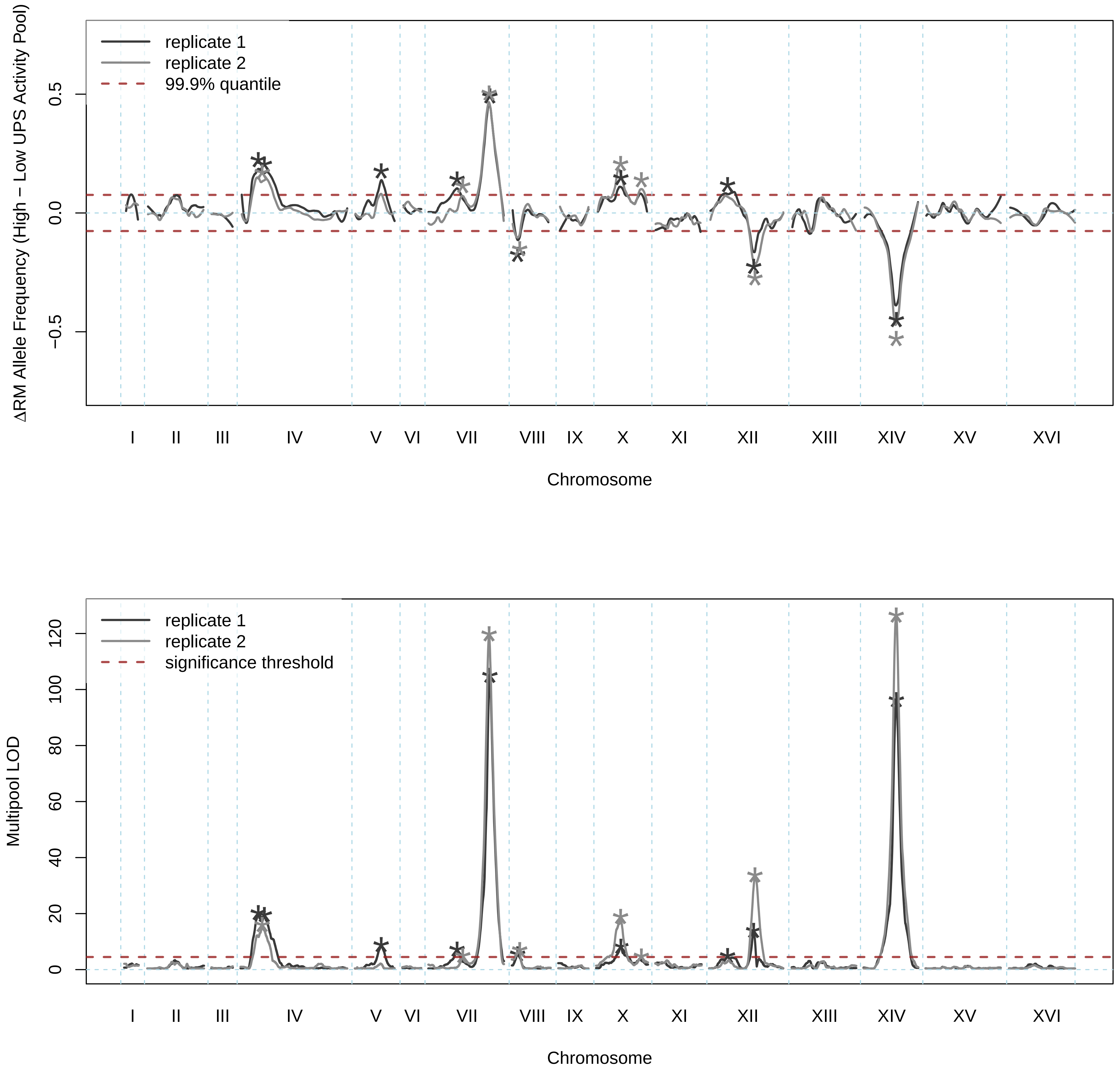
replicate 1
replicate 2
99.9% quantile
ΔRM Allele Frequency (High − Low UPS Activity Pool)
0.5
0.0
−0.5
I II III IV V VI VII VIII IX X XI XII XIII XIV XV XVI
Chromosome
replicate 1
replicate 2
significance threshold
Multipool LOD
0
20
40
60
80
100
120
I II III IV V VI VII VIII IX X XI XII XIII XIV XV XVI
Chromosome

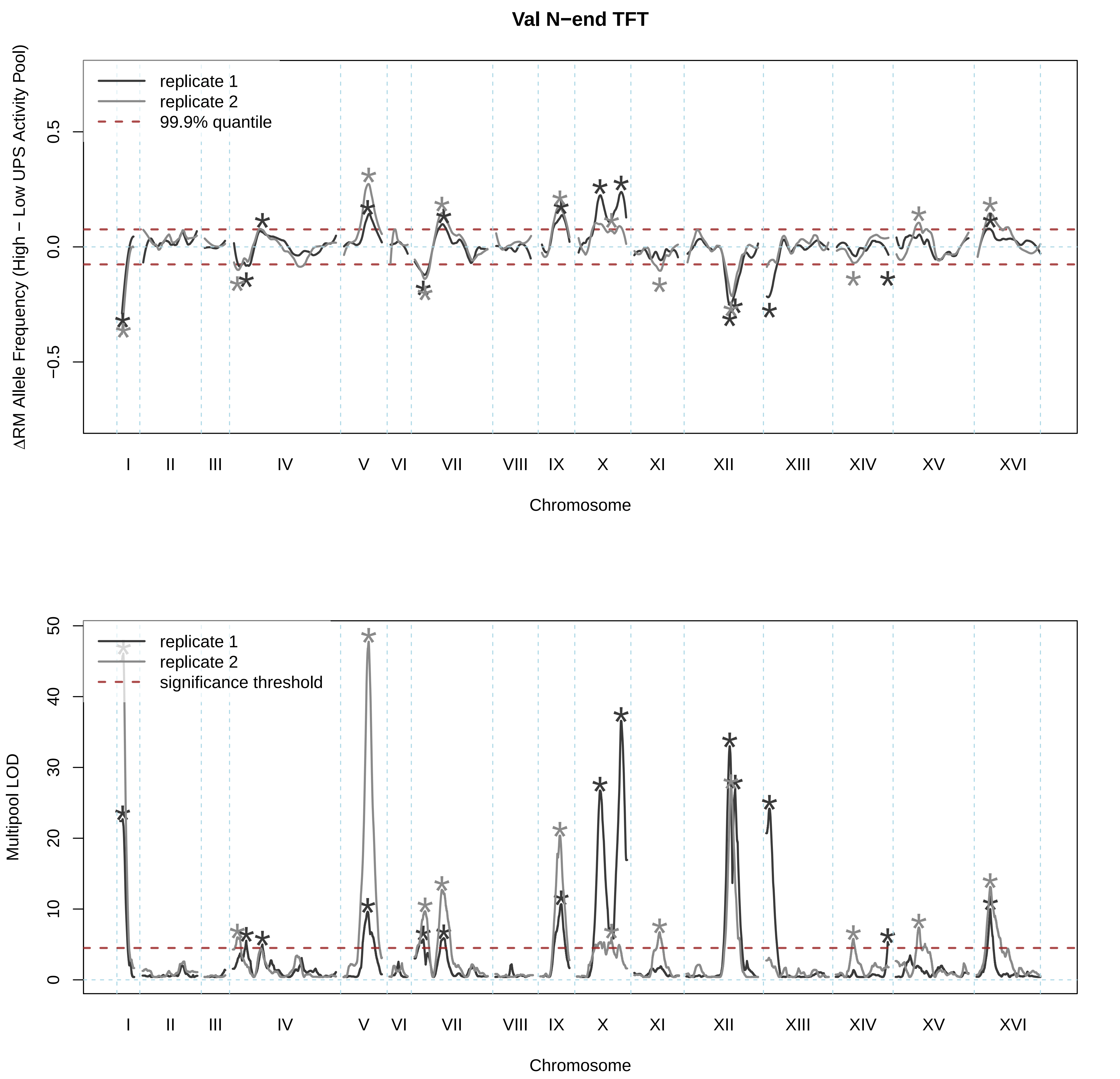

Val N−end TFT
replicate 1
replicate 2
99.9% quantile
ΔRM Allele Frequency (High − Low UPS Activity Pool)
0.5
0.0
−0.5
I
II
III
IV
V
VI
VII
VIII
IX
X
XI
XII
XIII
XIV
XV
XVI
Chromosome
replicate 1
replicate 2
significance threshold
Multipool LOD
50
40
30
20
10
0
I
II
III
IV
V
VI
VII
VIII
IX
X
XI
XII
XIII
XIV
XV
XVI
Chromosome
